# DeepCLIP: Predicting the effect of mutations on protein-RNA binding with Deep Learning

**DOI:** 10.1101/757062

**Authors:** Alexander Gulliver Bjørnholt Grønning, Thomas Koed Doktor, Simon Jonas Larsen, Ulrika Simone Spangsberg Petersen, Lise Lolle Holm, Gitte Hoffmann Bruun, Michael Birkerod Hansen, Anne-Mette Hartung, Jan Baumbach, Brage Storstein Andresen

## Abstract

Nucleotide variants can cause functional changes by altering protein-RNA binding in various ways that are not easy to predict. This can affect processes such as splicing, nuclear shuttling, and stability of the transcript. Therefore, correct modelling of protein-RNA binding is critical when predicting the effects of sequence variations. Many RNA-binding proteins recognize a diverse set of motifs and binding is typically also dependent on the genomic context, making this task particularly challenging. Here, we present DeepCLIP, the first method for context-aware modeling and predicting protein binding to nucleic acids using exclusively sequence data as input. We show that DeepCLIP outperforms existing methods for modelling RNA-protein binding. Importantly, we demonstrate that DeepCLIP is able to reliably predict the functional effects of contextually dependent nucleotide variants in independent wet lab experiments. Furthermore, we show how DeepCLIP binding profiles can be used in the design of therapeutically relevant antisense oligonucleotides, and to uncover possible position-dependent regulation in a tissue-specific manner. DeepCLIP can be freely used at http://deepclip.compbio.sdu.dk.

**Highlights:**

- We have designed DeepCLIP as a simple neural network that requires only CLIP binding sites as input. The architecture and parameter settings of DeepCLIP makes it an efficient classifier and robust to train, making high performing models easy to train and recreate.
- Using an extensive benchmark dataset, we demonstrate that DeepCLIP outperforms existing tools in classification. Furthermore, DeepCLIP provides direct information about the neural network’s decision process through visualization of binding motifs and a binding profile that directly indicates sequence elements contributing to the classification.
- To show that DeepCLIP models generalize to different datasets we have demonstrated that predictions correlate with *in vivo* and *in vitro* experiments using quantitative binding assays and minigenes.
- Identifying the binding sites for regulatory RNA-binding proteins is fundamental for efficient design of (therapeutic) antisense oligonucleotides. Employing a reported disease associated mutation, we demonstrate that DeepCLIP can be used for design of therapeutic antisense oligonucleotides that block regions important for binding of regulatory proteins and correct aberrant splicing.
- Using DeepCLIP binding profiles, we uncovered a possible position-dependent mechanism behind the reported tissue-specificity of a group of TDP-43 repressed pseudoexons.
- We have made DeepCLIP available as an online tool for training and application of proteinRNA binding deep learning models and prediction of the potential effects of clinically detected sequence variations (http://deepclip.compbio.sdu.dk/). We also provide DeepCLIP as a configurable stand-alone program (http://www.github.com/deepclip).

## INTRODUCTION

The massive technological progress in next generation sequencing (NGS) technologies has made sequencing affordable in the context of precision medicine and personalized health. NGS analysis enables identification of millions of sequence variants in each patient sample, increasing the need for *in silico* prediction of the functional consequences of a diverse range of variations. In particular, the effect of deep intronic sequence variants at the mRNA level through altered binding to RNA-binding proteins (RBPs) is difficult to predict *in silico* as existing tools’ predictions of functional outcomes of splicing are primarily based on the analysis of point mutations within or near exons(1–3). While some existing binding site prediction tools can work on sequences of any type, there is an unmet need for improved modelling of contextual dependencies other than structure that are important for correctly estimating the *in vivo* functionality of the binding sites. Extracted contextual information may form the basis for design of antisense oligonucleotide based therapies, which modulate RBP activity, such as splice-switching oligonucleotides (SSOs)(4–6). Thus, improving information on whether contexts act positively or negatively with regard to binding is an important area of research that will ultimately enable the development of novel therapeutic options in personalized medicine.

Sequencing technologies have also vastly expanded the wealth of information concerning protein binding to RNA when combined with cross-linking and immunoprecipitation (CLIP) techniques (7–9), which allow accurate mapping of protein binding sites in functional *in vivo* contexts. Classically, binding preferences or binding motifs have been represented by position frequency matrices (PFMs). Well-known *de novo* motif discovery tools such as MEME (10) and HOMER (11) output PFMs and base their motif detection and identification on the PFM concept. This approach to motif discovery implicitly assumes that such fixed-length motifs exist and that they function in a context-independent manner regarding the surrounding sequences. They further assume pairwise independence of the nucleotide frequencies within the motifs.

However, proteins that bind RNA typically do so in a context dependent manner. In particular, secondary structure may influence the binding of some RBPs(12). Information about double-stranded or single-stranded structure has been incorporated into MEMERIS(13), which is an extension of the MEME algorithm. Further structural dependencies have been incorporated into RNAcontext (12), which expands the information about secondary structure from simple double or single-stranded structures into paired, hairpin loops, bulges and internal or multi-loops, and unstructured contexts in order to further optimize the modeling of binding preference of RBPs. More recently, a graph-based modeling of structural and sequence binding preferences was introduced in the GraphProt (14) software, which out-performed RNAcontext on a set of diverse CLIP datasets using different CLIP methods. GraphProt uses RNAshapes (15) to predict the structures of RNA-sequences, which are then encoded into a hypergraph from which important structural features can be extracted. To improve the structure estimations, GraphProt extends the CLIP-derived sequences by 150 bp in each direction. Together with sequence features extracted only from the CLIP-derived binding sites, an overall model of binding preference is generated using support vector machines.

While inclusion of structural preferences may increase accuracy in prediction, these models still fail to capture other contextual dependencies affecting the *in vivo* functionality, such as a high density of protein binding sites nearby or localization within a specific functional region of the transcript, such as proximity to splice sites. For instance, exonic splicing enhancers (ESEs) that enhance splicing of exons by binding to SR proteins are enriched in exons, while exonic splicing silencers (ESSs) are underrepresented in exons. These observations have been used to generate ESE and ESS motifs from sequences enriched(16, 17) or depleted(16) in exons. Such contextual dependencies were recently introduced in the iONMF software(18), which uses integrative orthogonality-regularized nonnegative matrix factorization to incorporate multimodal information about CLIP-derived binding sites such as their position within the gene (5’UTR, CDS, exon, intron, 3’UTR), gene ontology, and presence of other protein binding sites determined by other CLIP studies, in addition to structural information, which improved performance for some datasets.

In recent years, deep learning techniques have been used to model protein binding. Deep learning has proven successful in various difficult classification tasks such as natural language processing (19), object recognition (20) and reconstructing brain circuits (21). Deep learning allows computational models composed of multiple processing layers to learn representations of data with multiple levels of abstraction (22). Deep learning models can identify dependencies and complex structures in very high-dimensional data - such as CLIP data - and have been used, for example, for predicting the effects of mutations in non-coding DNA on gene expression and disease (1, 23), predicting DNA function (24), mRNA coding potential (25), and prediction of subcellular locations of proteins (26). Starting with DeepBind (27), which uses convolutional neural networks (CNN) to classify bound sequences and non-bound, neural networks have also been used to directly model RBP preferences. Deepnet, a multimodal deep belief network incorporating 2D structure information (mDBN-) or both 2D and 3D structure information (mDBN+) along with a CNN architecture was introduced in the deepnet-rbp software (28), while more recently the iDeep framework combines the annotation data used by iONMF with a CNN into a multimodal neural network with increased accuracy in classification compared to iONMF(29). Even more recently, iDeepS was introduced as a replacement for iDeep to include analysis of 2D structural motifs much like GraphProt, using a combination of CNN and bidirectional LSTM layers akin to DeepCLIP’s architecture(30).

Previous models for RBP binding properties that consider contextual clues are focused either specifically on structural dependencies, which may fail to capture other important contextual dependencies, or on the presence of annotation data to aid in the task of classification. However, static annotations will not contribute to determining the effect of a mutation on the binding activity of proteins. For instance, a model, which relies heavily on clues from annotation data about the genomic region, such as location within an exon, will be unable to use this level of information to ascertain the effect of an exonic point mutation in which the context is maintained. Only iDeepS has a general-purpose LSTM layer able to model general context dependency, but it is supplemented with structural predictions from an external program and thus does not work on sequence data alone.

Understanding binding preferences is important for evaluation of the phenotypic impact of sequence variations. Mutations may alter the phenotype at several different levels, as in the case of missense mutations, which in addition to altering the amino-acid sequence, may also change the splicing pattern (31). Other mutations with less *visible* deleterious effects may abolish healthy splicing by altering the binding of RBPs, sometimes at somewhat distant sites. Splicing and the overall binding activity of RBPs is the result of a balance between positively and negatively acting elements that cooperate or compete for binding (32),(33), so even minor changes in RBP binding sites can change the outcome of splicing events.

Importantly, before they can be applied in predicting clinically important changes or functional elements to be targeted, binding models needs to be validated in the laboratory, using *in vitro* techniques such as RNA-protein affinity measurements and *in vivo* techniques such as minigene transfections and predictions need to be consistent with effects reported in clinically affected patients.

In this paper, we present DeepCLIP, a novel deep learning based tool for discovering protein-RNA binding sites and for characterizing binding preferences of RBPs. We demonstrate how it outperforms current state-of-the-art RBP binding analysis tools, and we show that DeepCLIP’s predictions provide information about high-affinity RBP binding sites and that it successfully predicts alterations of the RBP affinity for RNA-sequences when single nucleotide polymorphisms (SNP) or disease-causing mutations are introduced. This is reflected both in the binding profiles that show the region(s) important for RBP binding, and in the predictions of the sequences which indicate whether they are more similar to the “consensus” CLIP-sequence or to the genomic background. Last but not least, we have made DeepCLIP available as an online tool for training and application of protein-RNA binding deep learning models and prediction of the potential effects of clinically detected sequence variations (http://deepclip.compbio.sdu.dk/). We also provide DeepCLIP as a configurable stand-alone program (http://www.github.com/deepclip).

## MATERIALS AND METHODS

### DeepCLIP: more than just a motif discoverer

DeepCLIP is essentially a deep neural network that uses shallow 1D convolutional layers to find and enhance features of a set of presented sequences (26,34,35). This is followed by a Bidirectional Long Short Term Memory (BLSTM) layer (36, 37) which uses the extracted features and contextual information of the sequences to find areas of the RNA-sequences associated with RBP binding (Figure 1). Initially, the convolutional layers of DeepCLIP can be regarded as a collection of randomly generated PFMs of user-defined sizes that, as training progresses, learn to recognize important nucleotide patterns in the input data. When predicting, the convolutional layers score sequence segments according to their importance for the classification task. Pseudo-PFMs can be generated by collecting scored sequence patterns and counting the frequencies of different nucleotides at each possible position. We use the term pseudo-PFMs because each sequence used for the PFM generation is weighted by the squared output score given by the convolutional layers. In this way, the pseudo-PFMs will depict important class-specific nucleotide patterns. The BLSTM layer of DeepCLIP is used to generate a binding profile at the nucleotide level. The BLSTM layer consist of two LSTM layers that analyze “hidden” sequence representations (modified outputs of the convolutional layers) in a bidirectional manner (Figure S1).

**Figure 1.**
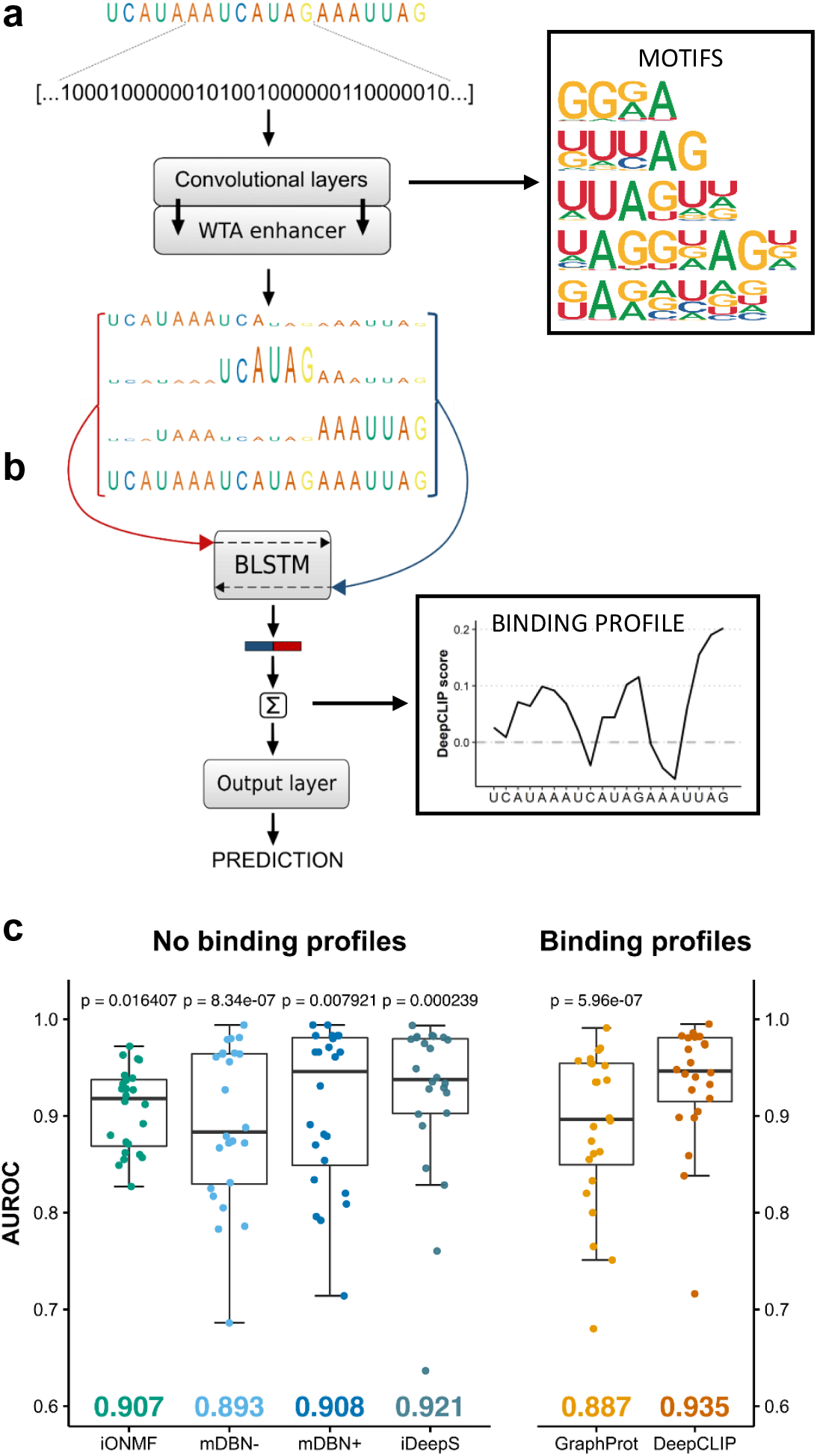
Classification performance of DeepCLIP surpasses competing methods. (a) One-hot embedded RNA-sequences function as input to the neural network and the 1D convolutional layers, which are further enhanced by a winner-takes-all (WTA) layer. (b) The outputs are concatenated with the input sequence and each segment of the concatenated arrays that corresponds to aligned bases is introduced to the following BLSTM layer as individual time-steps. The WTA-enhanced RNA-sequences are the three uppermost and the input sequence is in the bottom. DeepCLIP produces a prediction score, which is used during training, as well as binding motifs and a binding profile. (c) Boxplot of comparative analyses of DeepCLIP classification performance against other state-of-the-art tools. Area under receiver operator characteristic curve (AUROC) were measured in 10-fold cross-validation and statistical significance was computed through an exact Wilcoxon signed rank test. P-values above individual tools are from pair-wise comparisons with DeepCLIP. Mean AUROC score is indicated in the bottom.

The DeepCLIP tool takes a single or more RNA oligonucleotides (short RNA sequences) as input and predicts binding probability and calculates a binding profile. The main purpose of DeepCLIP is to identify binding sites of proteins in novel untested sequences using trained models that have extracted binding site information provided by CLIP data, to predict the effect of sequence variants on the binding, and to identify the importance of individual nucleotides for protein binding affinity. DeepCLIP is fast enough to run online on a web server (http://deepclip.compbio.sdu.dk/) and its Python code is also publicly available (http://www.github.com/deepclip).

### Training workflow of DeepCLIP

The core of DeepCLIP is a convolutional BLSTM network implemented in Theano (38) using the Lasagne library (39) and a few customized network layers and functions. DeepCLIP is a binary classifier that uses supervised learning to distinguish between unbound sequences and bound sequences derived from CLIP-experiments. The input to DeepCLIP consists of positive (bound) sequences, which are assigned to class 1, and negative (unbound) sequences assigned to class 0. These input sequences are converted into linearized one-hot encoded vectors, which serve as the actual input to the neural network layers. The DeepCLIP architecture is shown in Figure 1. By default, 80% of the input data is used for training while 10% is used for validation and the last 10% for testing (Figure S1d). DeepCLIP is trained by iterating over the training and validation sets several times (also called epochs). While training, performance is measured on the validation set after every epoch and the best performing model is saved. Early stopping can be applied to prevent training after a likely maximum performance has been obtained. The final performance of the saved model is measured on the test set, which contain data that have not previously been introduced to the model. When running in 10-fold cross-validation the input sequences are divided into 10 equal sized bins, and each bin is used once as a test set, once as a validation set and 8 times as part of the training set (Figure S1e).

### Encoding of sequence data

DeepCLIP processes sequence data as linearized one-hot encoded vectors. In the one-hot representation, the items of the vocabulary, *v* = *(A, C, G, U)*, are represented by vectors with lengths equal to the length of the vocabulary that each have a 1 in unique dimensions. The one-hot encoded bases are therefore independent of one another and are equally similar or dis-similar. It signals that no prior correlations between the bases are known. In this way, the network will determine correlations between the bases on its own (40). All input sequences are zero-padded until they have identical lengths and until the largest filter can conduct “full convolutions” as it is defined in the Lasagne documentation (39). Following vectorization of the bases of the sequences, the combined vectors are linearized, and the resulting one-dimensional data used as input to the neural network.

### Convolutional neural network layer

Convolutional layers consist of nodes that are only sensitive to a defined receptive field referred to as kernels or filters. Nodes of convolutional layers apply weight-sharing and sparse-connectivity, which means that the same filter can “view” all possible filter-sized segments of the input individually (41). As in previous work (26–29), the filters of the convolutional layers in DeepCLIP can be interpreted as motif detectors. The sizes of the filters of the convolutional layers are optional but we used ranges from 4-8 one-hot encoded bases. DeepCLIP only applies a single filter of each size. The strides of the filters are |𝑣|, so the filters only convolve patterns consisting of whole one-hot encoded bases. The convolutional layers apply the rectifier activation function (42). In this context, it means that only patterns that receive a score above zero can be assumed important for sequence identification.

The bias parameters of the convolutional layers are removed to ensure that only sequential areas containing one-hot encoded bases produce outputs above 0, which implies that these areas are deemed more important in the further processing. The initial weights of the convolutional nodes are set to 0.01. The filter sizes allow for diverse sequence patterns of various length to be incorporated into the model. The output vectors of convolutional layers in DeepCLIP are vectors containing values between 0 and ∞. Before the output vectors are passed to the BLSTM layer they undergo a so-called WTA-enhancement (Winner Take All-enhancement), which is described below.

The single highest values of the different output vectors are multiplied by 2 which is followed by a squaring of the vectors to enhance differences between high low values. These squared vectors have now been WTA-enhanced, where the “winners” are the highest values in the output vectors (see Figure 1 for a thorough explanation). The WTA-enhanced vectors are used for a recreation of the original one-hot embedded sequences where the one-hot values are defined by the WTA-enhanced vector elements. For each convolutional layer, a WTA-enhanced sequence is created and concatenated in a manner that makes it possible to process each numerical base representation of the sequences as individual steps in the following BLSTM layer (see Figure 1).

In this way, the convolutional layers help guide the attention of the BLSMT layer. The pseudo-PFMs created by the DeepCLIP tool that depict the important patterns in the RBP-bound sequences, are based on the specific sequential areas that only relate to class 1. Meaning, if an area of a sequence is, by the BLSTM layer, predicted as being associated with class 0, any convolutional output values in the sequential area will be zeroed out and therefore will not be a part of the pseudo-PFM calculation. The filters of the convolutional layers of the model with the best performance in the 10-fold cross validation were extracted and used for creation of pseudo-PFMs. The top 1000 sequences with respect to the predictions were used for the pseudo-PFM calculation.

### Bidirectional LSTM layer

Long short-term memory (LSTM) networks have already proven successful in biological sequence analysis (26, 43). DeepCLIP uses a single BLSTM layer that processes the WTA-enhanced sequences (Figure 1). In BLSTM layers, the input sequences are presented forwards and backwards in two separate LSTM layers that are connected to the same output layer (37). The implementation of a single LSTM layer in DeepCLIP is given by equations (1–9):

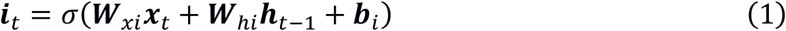

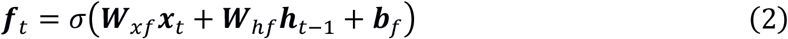

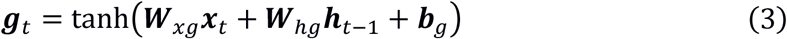

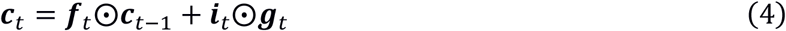

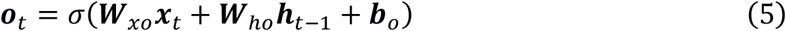

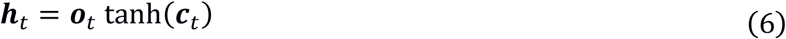

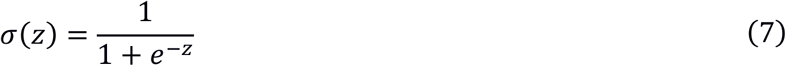

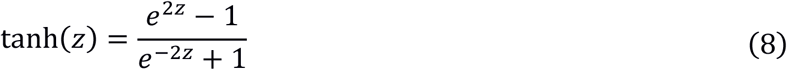

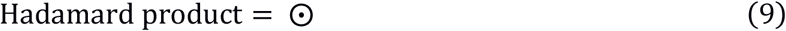

where ***i***, ***f***, ***g***, ***o*** and 𝒄 are the input gate, forget gate, modulatory gate, output gate and cell, respectively. ***x***_𝑡_is the input vector at timestep 𝑡, ***W***_𝑥𝑖_is the input-input gate weight matrix, ***W***_ℎ𝑖_ is the hidden-input gate weight matrix, 𝒃_𝑖_ is the bias of the input gate and ***h***_𝑡−1_ is the hidden output vector from timestep 𝑡 − 1. The same logic applies for the remaining gates. 𝜎 is the sigmoid activation function, 𝑡𝑎𝑛ℎ is the hyperbolic tangent activation function and the Hadamard product indicates elementwise multiplication. The hidden output vector of a LSTM memory block is 𝒉_𝑡_ at timestep 𝑡.

The forward LSTM reads an input sequence with length 𝑇 from ***x***_1_ to ***x***_𝑇_ and the backward LSTM reads the same input sequence from ***x***_𝑇_ to ***x***_1_. The forward LSTM layer produces forward hidden vectors, 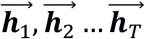 and the backward LSTM layer produces backward hidden vector 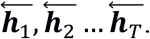 Here, the output of the backwards LSTM layer has been reversed, so the outputs of forward and backward LSTM layers go from ***x***_1_ to ***x***_𝑇_. The hidden vector of the BLSTM layer at time step 𝑡, ***h***_𝑡_, is given by the concatenation of the forward hidden vector and the backward hidden vector 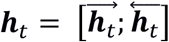 (44).

The output sequence of a BLSTM layer given an input sequences of length 𝑇 can be seen as a matrix, ***H*** = (***h***_1_, ***h***_2_, … ***h***_𝑇_), where each ***h***_𝑡_ is a row. Each ***h***_𝑡_ contains information about the whole input sequence with a strong focus on the parts surrounding the 𝑡^𝑡ℎ^ input vector (44). In DeepCLIP, each ***h***_𝑡_ represents a base of a sequence that has knowledge of the surrounding sequence. These context-aware representations of sequences function as input to the output layer where the final prediction is calculated. Dropout is applied on ***H***, which means that recurrent connections are not affected.

### Output layer, binding profile and prediction

The output layer consists of a single fixed node without a bias parameter that uses the sigmoid activation function. By “fixed” we mean that the parameters of the node do not update when training. The initial weight of the node is set to 1.0, which means that the node is forced to associate positive values with class 1 and negative values with class 0. The input to the output layer, given a single input sequence, is the vector that results from a row summation of the matrix ***H*** where segments based on zero-paddings are zeroed out. By zeroing out these segments it is ensured that only areas that contain one-hot embedded bases are used for the prediction of the given sequence. Basically, the prediction of a given RNA-sequence is the sum of the output values of the BLSTM layer inserted into a sigmoid activation function.

In terms of the BLSTM output, if a ***h***_𝑡_ is mainly positive the base at its specific position is associated with the “consensus patterns” of the sequences derived from the CLIP experiments. If a ***h***_𝑡_ is primarily negative, the base at its specific position is more associated with random background sequences derived from the genome. And if the values of a ***h***_𝑡_ sums to ∼0, the base at its position could belong to both classes. This approach makes DeepCLIP able to highlight sequential areas that differ from genomic background and thereby able to identify *in vivo* binding sites. The binding profiles are constructed using the input vectors of the output layer, where all the zero-padding has been removed.

### DeepCLIP default settings

DeepCLIP uses ADAM(45) (𝛼 = 0.0002, 𝛽_1_ = 0.9, 𝛽_2_ = 0.999, 𝜖 = 10^−8^) for gradient descent optimization. For the BLSTM layer, the parameters are sampled from a Gaussian distribution with 𝜇 = 0.0 and 𝑠𝑡𝑑 = 0.01. DeepCLIP uses binary cross entropy as loss function and employs dropout in order to avoid overfitting. The most optimal network weights based on AUROC performance on the validation set are saved during training. Dropout is applied to the BLSTM layer (10%).

### Generation of background sequences

Background sequences can be either supplied by the user as sequences or generated automatically by DeepCLIP from positive binding sites in one of two ways. The default way is to supply positive binding site in BED format and then generate a random set of identically sized genomic regions as the positive binding site and randomly placing each within the same gene as the corresponding positive binding site such that no background regions overlap either a positive binding site or another background region. Alternatively, background sequences can be generated from positive binding sites by scrambling the input sequences. When positive binding sites are supplied in BED format, they can optionally be expanded on each size, or fixed to a certain width, or a combination of these. In this paper, we have used the random genomic background method to most accurately obtain *in vivo* non-bound sites.

### Analysis of Area Under Receiver Operator Curve classification performance

Comparison of Area Under Receiver Operator Curve (AUROC) classification performance was performed on the dataset compiled in the GraphProt paper(14). In order to minimize computational complexity, we took the performance numbers of alternative models on this dataset as they were reported in the previous studies(14,18,28). We could not compare to iDeep(29), as they did not provide numbers for the same datasets, and did not provide any way of producing the multimodal data required as input. We ran iDeepS on 101 nt sequences from the GraphProt dataset by expanding the peak-areas on either side until the sequence was 101 nt. We ran iDeepS on these sequences with the same number of epochs that we used to train DeepCLIP models. We generated 10-fold cross-validation sets where one set was held out for testing one time, and used in training the other 9 times, in order to obtain comparable performance measures across the full datasets. We ran DeepCLIP on the peak-area sequences of the datasets in a 10-fold cross-validation, such that each site was held out exactly once for validation during training, and once for final testing, while being used for actual training the remaining eight times. Model performance was measured for each dataset using the performance measure tool “perf” as used by GraphProt on the combined predictions from all CV cycles. Additionally, AUROC confidence intervals were estimated using the DeLong algorithm as implemented in the “pROC” R package(46).

### Analysis of additional public CLIP data

Binding sites from an eCLIP study of SRSF1(47) in K562 cells were downloaded and the overlap between two replicates were extracted to create a set of non-redundant binding sites. These were used for constructing the bound dataset, with a matched genomic background as control. We then trained an SRSF1 DeepCLIP model on this dataset using same running parameters as previous models, with 50 training epochs and early stopping after 5 epochs. TDP-43 binding sites were downloaded from POSTAR2(48) and non-redundant input sites were used to train a DeepCLIP model, again using identical running parameters as previous models, but adjusting the number of training epochs to 50 and early stopping after 5 epochs to account for the much larger training set. Similarly, hnRNP A1 binding sites(49), were used to train a DeepCLIP model on binding sites with p-values below 0.01 using default parameters with 200 training epochs and early stopping after 20 epochs. In all cases, 10-fold cross-validation was used to identify the best performing model.

### Analysis of TDP-43 repressed pseudoexons

Pseudoexons activated by conditional knock-out of TDP-43 in mice(50) were analyzed with DeepCLIP by first extracting the sequence of the pseudoexon along with 100 nt of the neighboring intronic sequences. These sequences were then used to produce binding profiles by using a sliding window approach to produce raw DeepCLIP profiles of smaller segments, taking the value of the central nucleotide to build a binding profile covering the entire length of the sequence. Subsequently, regions corresponding to the 25 first and last nucleotides of the exons along with the 50 first and last nucleotides of the neighboring introns were extract in order to analyze TDP-43 binding to the acceptor and donor splice site regions.

### Minigene generation

*ACADM* exon 5 minigenes were identical to the previously used wt *ACADM* minigene (33), with the exception of nucleotide variants at positions corresponding to c.361, c.362, and c.363 with exon 5. These variants were introduced as previously described (33).

The *ACADM* exon 6 wt minigene was generated from genomic DNA by amplifying the complete exon 6 (81 bp) along with 864 bp of intron 5 and 603 bp of intron 6 and subsequent cloning into the pSPL3 vector (Gibco BRL) using the BamHI and XhoI restriction sites. For amplification we used the forward primer 5’-TCGAGAATTCAGGAGCA-3’ and the reverse primer 5’-CTCCACTAAATAGAGC-3’. The IVS6+7A>G mutation was introduced by GenScript (GenScript, Piscataway, NJ, USA).

### *ACADM* exon 5 minigene transfections and RT-PCR

HEK-293 cells were seeded in 3.5 cm^2^ 12-well plates (Nunc) at a density of 4×10^5^ cells/well 24 hours prior to transfection. In each well, cells were transiently transfected using X-tremeGENE 9 DNA Transfection Reagent (Merck): 0.3 µg of one of the *ACADM* exon 5 minigenes c.362C (wildtype), c.361C, c.361G, c.361T, c.362A, c.362G, c.362T, c.363A, c.363C, or c.363G. After 48 hours of incubation following minigene transfection, cells were harvested using QIAzol Lysis Reagent (Qiagen), followed by phenol/chloroform extraction of total RNA. Reverse transcription was performed using the High Capacity cDNA Reverse Transcription Kit (Thermo Scientific). Splicing patterns were analyzed by PCR amplification, using TEMPase Hot Start DNA Polymerase (Ampliqon), and agarose gel electrophoresis. We used the *ACADM* exon 5 minigene specific primers: MCTEST2AS (5′-AGACTCGAGTTACTATTAATTACACATC-3′) and MC242S (5’-CCTGGAACTTGGTTTAATG-3’). PCR products were quantified according to molar ratios by capillary gel electrophoresis on a Fragment Analyzer™ instrument (Agilent), and visualized on 1.5% agarose gels. Experiments were performed in triplicates.

### *ACADM* exon 6 minigene transfections with siRNA mediated knock-down of TDP-43 and RT-PCR

Knockdown of TDP-43 was obtained by performing reverse transfection during initial seeding of cells and another transfection 48 hours later. Both transfections were performed using Lipofectamine RNAiMAX Transfection Reagent (Thermo Fisher Scientific) and 40 nM of siRNA targeting *TARDBP* (L-012394-00-0020, Dharmacon) or non-targeting siRNA (D-001810-10-20, Dharmacon). HeLa cells were seeded in 3.5 cm^2^ 12-well plates (Nunc) at a density of 1.5×10^5^ cells/well 24 hours prior to minigene transfection. In each well, cells were transiently transfected using X-tremeGENE 9 DNA Transfection Reagent (Merck): 0.4 µg of one the two *ACADM* exon 6 minigenes: WT or +7A>G. After 48 hours of incubation following minigene transfection, cells were harvested using QIAzol Lysis Reagent (Qiagen), followed by phenol/chloroform extraction of total RNA. Reverse transcription was performed using the High Capacity cDNA Reverse Transcription Kit (Thermo Scientific). Splicing patterns were analyzed by PCR amplification, using TEMPase Hot Start DNA Polymerase (Ampliqon), and agarose gel electrophoresis. We used the minigene specific primers: SD6 (5’-TCTGAGTCACCTGGACAACC-3’) and SA2 (5’-ATCTCAGTGGTATTTGTGAGC-3’). PCR products were quantified according to molar ratios by capillary gel electrophoresis on a Fragment Analyzer™ instrument (Agilent), and visualized on 1.5% agarose gels. Knockdown of TDP-43 was validated by SDS-PAGE and Western Blotting and membranes were probed with antibodies anti-TDP-43 (10782-2-AP, ProteinTech) and as a loading control anti-HPRT (HPA006360, Merck). Experiments were performed in triplicates.

### SSO co-transfection with *ACADM* exon 6 minigenes

HeLa cells were reverse transfected in duplicates with 40 nM SSO using Lipofectamine RNAiMAX Transfection Reagent (Thermo Fisher Scientific) according to the manufacturer’s protocol, and seeded in 3.5 cm^2^ 12-well plates (Nunc) at a density of 2×10^5^ cells/well 24 hours prior to minigene transfection. SSOs were phosphorothioate oligonucleotides with 2′-O-methyl modifications on each sugar moiety (LGC Biosearch Technologies): SSO1 (5’-UAAGUGUGAAAUAAAGCGGCAGUUA-3’), SSO2 (5’-AGUGUGAAAUAAAGCGGCAGUUACA-3’), or a control SSO without any human target sites: 5’-GCUCAAUAUGCUACUGCCAUGCUUG-3’. Cells were transiently transfected using X-tremeGENE 9 DNA Transfection Reagent (Merck): 0.4 µg of one of the two *ACADM* exon 6 minigenes: WT, or +7A>G. After 24 hours of incubation following minigene transfection, cells were harvested using QIAzol Lysis Reagent (Qiagen), followed by phenol/chloroform extraction of total RNA. Reverse transcription was performed using the High Capacity cDNA Reverse Transcription Kit (Thermo Scientific). Splicing patterns were analyzed by PCR amplification, using TEMPase Hot Start DNA Polymerase (Ampliqon), and agarose gel electrophoresis. We used the minigene specific primers: SD6 (5’-TCTGAGTCACCTGGACAACC-3’) and SA2 (5’-ATCTCAGTGGTATTTGTGAGC-3’). PCR products were quantified according to molar ratios by capillary gel electrophoresis on a Fragment Analyzer™ instrument (Agilent), and visualized on 1.5% agarose gels. Experiments were performed in triplicates.

### Surface plasmon resonance imaging method

Biotinylated oligonucleotides were immobilized on a Senseye G strep (SSENS) sensorchip in a 2×4×12 array by continuous flow in a CFM 2.0 printer (Wasatch microfluidics). The oligonucleotides were diluted in 1XTBS to a concentration of 1 µM and spotted for 20 min followed by 5 minutes washing with TBS + 0.05% Tween-20. The sensor chip was transferred to the MX-96 (IBIS technologies), and the system was primed with SPR buffer (10 mM Hepes/KOH pH 7.9, 150 mM KCl, 10 mM MgCl_2_ and 0.075% Tween-80). Surface plasmon resonance imaging (SPRi) by IBIS MX-96 was used to measure the kinetics of recombinant hnRNP A1 (ab224866, Abcam), SRSF1 (GenScript, Piscataway, NJ, USA) and TDP-43 (R&Dsystems, AP-190) binding to the immobilized RNA oligonucleotides. Binding was measured in real time by following changes of the SPR angles at all printed positions of the array during 10 min. injections of recombinant protein over the entire surface. Seven injections of a 2-fold titration series from 6.25 to 400 nM protein was injected in sequence from the lowest concentration to the highest. Before adding protein to the chip, residual background binding was blocked by injecting 20mg/ml BSA in SPR buffer onto the chip for 10 minutes. A continuous flow of SPR buffer flowed over the surface before, between and after the protein injections, to measure baseline and dissociation kinetics. Dissociation was measured for 8 minutes, by injecting SPR buffer over the chip at a rate of 4 µl/sec. Responses for a calibration curve were created after the concentration series by measuring SPR responses from defined dilutions of glycerol in running buffer (ranging from 5 to 0 % glycerol) and of pure water as defined by the automated calibration routine of IBIS MX-96.

Data analysis: The SPRi data was imported into SPRINTX software (v. 2.1.1.0, IBIS technologies), calibrated, reference subtracted, and the baseline of the responses before all injections were zeroed. The time starting point was aligned at the beginning of each new injection. Then the data were exported to Scrubber 2 (v 2.0c, Biologics Inc.). Binding curves for all chip positions where binding was observed were fitted globally to the integrated rate equation that describes simple first order 1:1 binding kinetics to obtain kinetic association rate (ka), dissociation rate (kd) as well as the Rmax for the binding model. For hnRNPA1 and TDP-43 a 1:2 biphasic model was calculated and fitted. ClampXP (version 3.50, Biosensor Data Analysis) was used with a bimodal model to fit the binding data. The secondary Ka and Kd parameters were fixed to 1e-7 M, due to very low secondary association and dissociation. Primary binding parameters and ligand concentration (Rmax) was set to float. SPRi measurements were performed twice, with technical duplicates being used for model fitting each time.

### Statistical analyses

All statistical analyses were performed in R (version 3.5.3). We used the default wilcox.test() for two-tailed Wilcoxon rank sum and Wilcoxon signed rank tests. Linear regression was carried out using ggplot2 (version 3.2.1) and geom_smooth(method=lm, colour=“red”, se=TRUE), with correlation significance analysis using the default cor.test() with method=“spearman”.

## RESULTS

### DeepCLIP outperforms structural and multimodal models from sequence data alone

DeepCLIP is a neural network that combines shallow convolutional layers with a small bidirectional long short-term memory network to produce both a binding profile and a classification score ranging from 0 to 1 (Figure 1, S1a-c). Models are created by training a network on a set of known binding sites and a set of background genomic sequences (Figure S1d,e), which can optionally be generated by DeepCLIP by providing binding locations instead of raw binding sequences.

To ascertain DeepCLIP’s classification performance on a standardized dataset, we generated models from the curated CLIP datasets (8,9,51–59) used in the GraphProt publication (14), which has previously been used in other studies (18, 28). First, we trained DeepCLIP models in a 10-fold cross-validation scheme using 50-500 epochs depending on the size of the individual dataset with early stopping after 10% of the maximum number of epochs (Table S1). Next, we measured area under receiver operator characteristic curve (AUROC) using the standard method of 10-fold cross-validation and the combined performance across the 10 different sets (Figure S1e, Table S2). Importantly, DeepCLIP does not directly model structure. Consequently, we used the peak area alone, which is akin to the viewpoint mechanism as adopted in GraphProt. We compared the performance of our models with the performance numbers reported in the earlier studies describing GraphProt, iONMF, and deepnet-rbp (mDBN- and mDBN+). To obtain AUROC values for iDeepS, we performed a 10-fold cross-validation using the curated CLIP-datasets, as this was omitted in the iDeepS paper, by extending the peak area to 101nt per the input requirement for iDeepS. We found that DeepCLIP was the overall best classifier in every pair-wise comparison and when looking at the mean AUROC score, underscoring that DeepCLIP performs well on a broad set of data. Furthermore, DeepCLIP had more narrow distributions of scores with fewer low-scoring datasets and a majority of datasets scoring above 0.9 (Figure 1c). DeepCLIP consistently ranked among the best classifiers on individual datasets (Figure S2a, Table (S3)1), even without additional contextual data, i.e. working on the sequence input data alone. In total, DeepCLIP had the best performance for 14 of the 24 datasets, and second best for 6 datasets. For TAF15, mDBN+ and DeepCLIP had identical scores when rounding to the 3^rd^ decimal. DeepCLIP was also a better classifier than DeepBind ((pseudo)median 15.16 %-points (CI-95%: 7.76-24.17), Wilcoxon signed-rank p=0.0004883), on the 12 datasets for which DeepBind models are available (Figure S2c,d). Note, however, that DeepBind models were trained on very different input data, which makes it difficult to ascertain the true performance of DeepBind.

We did not observe any significant differences in DeepCLIP AUROC scores between the CLIP methods (Figure S3a) or a significant correlation to the number of bound sites (Figure S3b, p = 0.531), but a tendency towards improved performance on the nucleotide-resolution CLIP datasets (iCLIP and PAR-CLIP) versus HITS-CLIP did appear. Due to the limited number of HITS-CLIP and iCLIP datasets it is not clear whether this is a significant difference or not. DeepCLIP performed well on all datasets regardless of size and CLIP method, with the exception of ALKBH5, which is a problematic dataset for all methods that do not rely on additional metadata, presumably due to non-specific binding that may take place in cooperation with a number of other factors that target the factor to specific regions within the transcript. DeepCLIP is thus a robust classifier of *in vivo* binding sites using only sequence data. It compares favorably to models employing external structural information and annotation data in addition to sequence data.

### DeepCLIP models predict binding motifs

Although the classifications of DeepCLIP are based entirely on the values of the binding profiles, binding motifs can be assessed from the CNN filters incorporated in the network architecture. The motif of each filter was generated using the patterns from the 1,000 input sequences that produced the highest DeepCLIP classification score. We found that DeepCLIP produces binding motifs that are in agreement with previously published motifs (60–65), illustrating that DeepCLIP’s classification performance is not simply a result of learning how to recognize the background sequences, but depends on the binding preferences of the RBP in question (Figure S4). Additional model performance and CNN filter motifs are available (Table S2, Figure S5-S28).

### DeepCLIP predictions and binding profiles explain splicing mutations

Splicing of mRNA is regulated by binding of RBPs to the nascent pre-mRNA. To test DeepCLIP’s ability to predict effects of mutations on splicing we generated new models for the splicing factors hnRNP A1 and SRSF1 based on previous CLIP studies (47, 49) with DeepCLIP-generated background sequences in order to demonstrate DeepCLIP’s performance on novel datasets. We ran 10-fold cross-validation (Figure S29 and S30) using the same input parameters as previously, which produced CNN filters (Figure 2a) demonstrating binding motifs similar to previous studies (56,64,66,67). We then used the best performing model to predict binding of hnRNP A1 to a set of exonic point mutations (2), grouped into mutations known to cause skipping and mutations known to not cause skipping (Figure 2b, Table S4).

**Figure 2.**
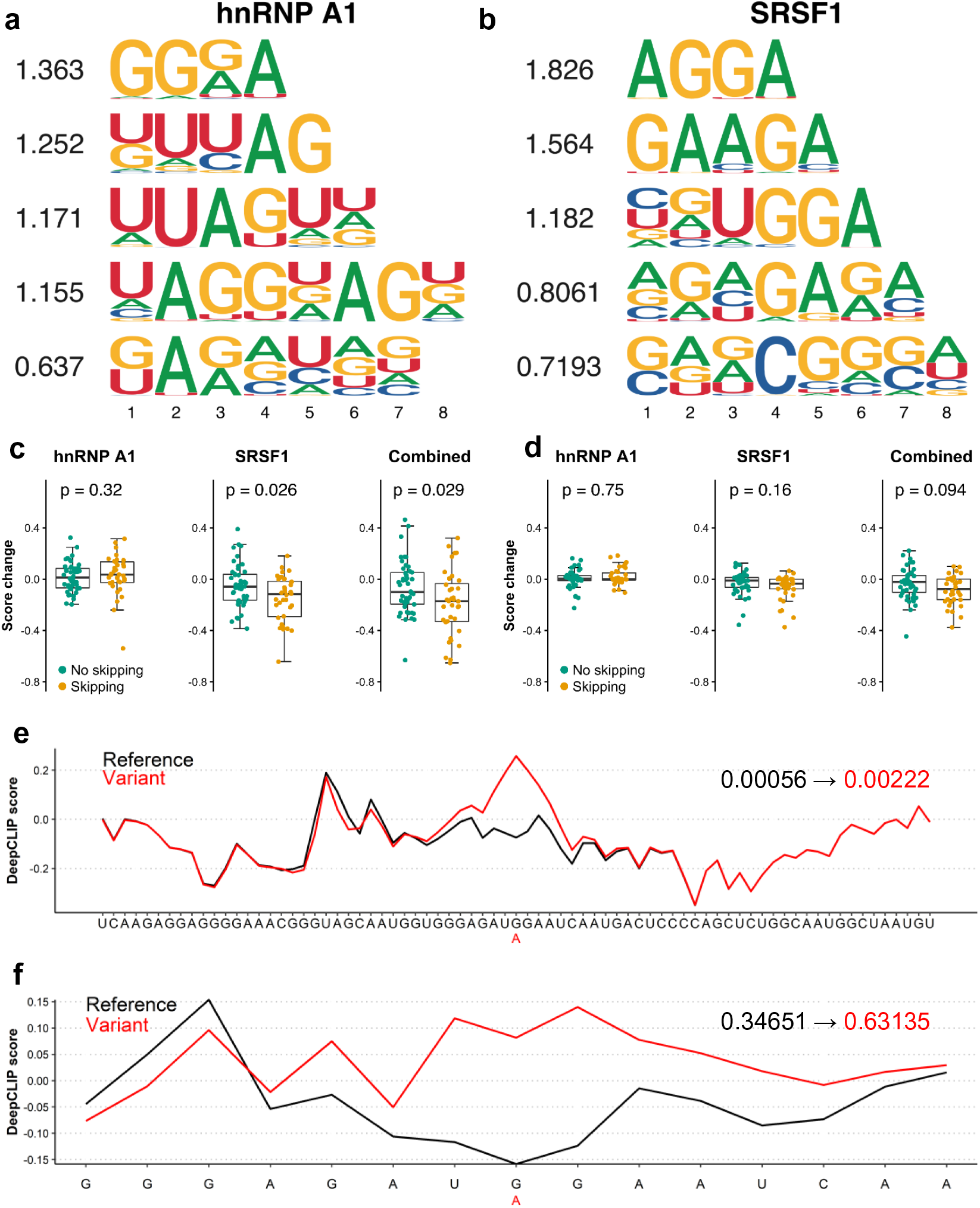
DeepCLIP models of hnRNPA1 and SRSF1 used to analyze splicing mutations. (a) The CNN filters trained in the hnRNP A1 DeepCLIP model. (b) The CNN filters trained in the SRSF1 model. (c) Box-plots of the prediction score change of DeepCLIP hnRNP A1 model (left), SRSF1 (middle) and combined (right, SRSF1 score minus hnRNP A1 score) predictions on 15 nt sequences representing wt and mutant versions exons that are skipped upon mutation (yellow, n = 37 sequence-pairs) and exons that remain included in the mutant version (green, n = 46 sequence-pairs). Two-tailed Wilcoxon rank sum test p-value is indicated above. Box-plot elements are defined as center line: median, box limits: upper and lower quartiles, whiskers: 1.5x interquartile range. All data points are shown, outliers are not highlighted. (d) Same as (c) but with 75 nt sequences. (e) DeepCLIP profiles of wt (black) and mutant (red) 75 nt sequence representing the *ATP7A* exon 20 +103G>A mutation. The overall DeepCLIP prediction scores are indicated in bold within the plot. (f) Same as (e), but with 15 nt input sequence.

We used the 15-mer sequences used by Raponi et al. in their work describing splicing because this represents the length of a typical RNA oligonucleotide used in affinity purification experiments to measure the binding of protein to RNA. Interestingly, we observe no significant difference in the overall change of hnRNP A1 scores for skipping and non-skipping mutations (Figure 2c, left, Wilcoxon signed rank p = 0.3152), suggesting that hnRNP A1 is not a general regulator of these splicing events. This is to be expected, since only a subset of these events is likely to be meditated by altered binding of hnRNP A1.

When scoring the same 15 nt oligonucleotides with the best performing SRSF1 model, we observe that the SRSF1 scores are significantly different between the two groups (Figure 2c, mid, Wilcoxon signed rank p = 0.02585). Also, when combining the hnRNP A1 scores with the SRSF1 scores, we find that the groups were significantly different (Figure 2c, right, Wilcoxon signed rank p = 0.02913). Importantly, the scores of the mutations known to cause skipping were decreased, consistent with the known role of SRSF1 as a positive regulator of exon inclusion.

To investigate whether the hnRNP A1 and SRSF1 models improve with extended sequence context, we expanded the 15-mer sequences from the middle and out to a length of 75 bp, the maximum sequence length used during training. This resulted in less pronounced changes that were not statistically significant (Figure 2d, Table S3), although the combined score indicated that the combined effect of losing SRSF1 binding and gaining hnRNP A1 binding was retained to a higher degree. This is likely caused by DeepCLIP’s classification being based on the total binding profile resulting in diminished differences as the sequence is expanded. This is exemplified by the *ATP7A* c.3904G>A exon skipping mutation located at +103 in exon 20 of *ATP7A,* which results in an overall score change of +0.00166 between the wt (0.00056) and mutant (0.00222) 75 nt long sequences (Figure 2e), but a score change of +0.28484 (from 0.34651 to 0.63135) when the 15 nt long sequence is used (Figure 2f). Importantly, both binding profiles predict a localized increase in hnRNP A1 binding to the mutant, show-casing the relevance of using importance profiles when analyzing sequence data and not simply an overall prediction score.

### DeepCLIP hnRNP A1 and SRSF1 prediction scores correlate with exon inclusion levels of a known SRSF1-dependent exon

We have previously characterized splicing of *ACADM* exon 5, which shares sequence similarity with *SMN1* exon 7 and we identified a similar regulation with splicing ultimately relying on the balance of SRSF1 and hnRNP A1 binding (33). A prevalent disease-causing c.362C>T mutation reduces the strength of a SRSF1 binding ESE, allows hnRNPA1 binding and causes exon 5 skipping. To test DeepCLIP models of hnRNP A1 and SRSF1 in relation to splicing of *ACADM* exon 5, we generated minigenes with all possible variants at positions c.361, c.362, and c.363 located down-stream of the CAG core motif (Figure 3a). DeepCLIP scores were obtained from the sequences using a window of 36 bp on each side of the 3 positions, totaling 75 nt. We then transfected the minigenes in HEK293 cells and measured exon inclusion levels (PSI) using RT-PCR and gel-electrophoresis (Figure 3b). We observe a strong negative correlation (Spearman’s *ρ* = −0.939, *p* < 2.2e-16, Figure 3c) between the hnRNP A1 prediction score and the observed inclusion of *ACADM* exon 5, and a strong correlation between the SRSF1 prediction score and the observed inclusions (Spearman’s *ρ* = 0.770, *p* = 0.01367, Figure 3d). The observed correlation is also very strong for the combined scores (Spearman’s *ρ* = 0.915, *p* = 0.0004667, Figure 3e), in agreement with the hypothesis that the overall inclusion level is a result of the balance between positive and negative factors. The same is true when we use the SRSF1 model generated from the GraphProt dataset (Figure S31), but not when we perform the same analysis with EX-SKIP (2) on the full exon (Spearman’s *ρ* = −0.177, *p* = 0.625, Fig S32a). When we use SPANR (1) we do observe a stronger positive correlation than with EX-SKIP (Spearman’s *ρ* = 0.552, *p* = 0.1043, Fig S32b), but all variants are predicted to have an inclusion level between 81.9% and 82.8%. This conflicts directly with the observed level of exon skipping induced by the disease-causing c.362C>T mutation in patient cells (33) as well as with the observed splicing pattern of the minigenes tested.

**Figure 3.**
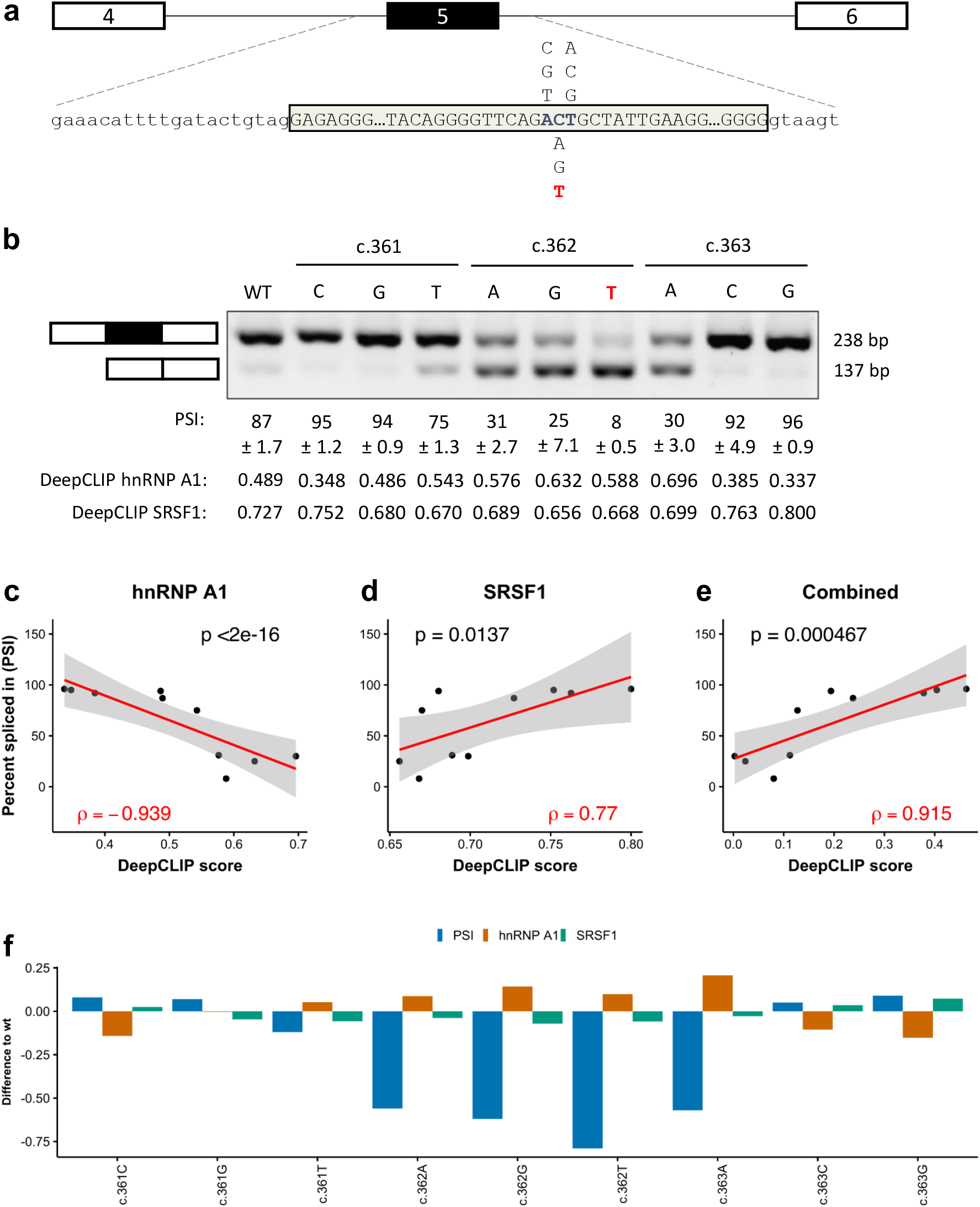
DeepCLIP models successfully model *ACADM* minigene splicing results. (a) Minigene schematic and location of variants tested, reference in blue. The disease-causing mutation is indicated in red. (b) Splicing of minigenes determined by RT-PCR. Estimates of mean PSI (n = 3) is indicated below, along with 95% CI size. (c) Scatter plot of PSI and DeepCLIP hnRNP A1 score with linear regression (red line, n = 10) and 95% confidence interval (shaded area). (d) Same as (c) but with DeepCLIP SRSF1 score instead. (e) Same as (c) and (d), but showing the DeepCLIP SRSF1 score minus the DeepCLIP hnRNP A1 score. (f) Barplot showing the difference to wt for the minigene PSI and DeepCLIP prediction scores for hnRNP A1 and SRSF1. Spearman’s correlation coefficient is indicated in (c), (d), and (e).

Overall, when the hnRNP A1 DeepCLIP model predicts an increase in hnRNP A1 binding, there was a decrease in exon 5 inclusion (Figure 3f). In particular, the high degree of exon skipping of all c.362 variants relative to wt were reflected by increases in hnRNP A1 scores (Figure 3b and f). The c.363A variant is predicted to abolish binding of SRSF1 and increase binding of hnRNP A1 and the minigene analysis demonstrates predominant skipping in agreement with this. Like hnRNP A1, many of the variants predicted to lose SRSF1 binding show increased skipping consistent with a loss of ESE activity.

The data indicate that while single DeepCLIP models capture binding preferences of individual proteins, the scores are additive and can be used to model effects of multiple proteins interacting in antagonistic and synergistic ways.

### DeepCLIP binding profiles can guide the design of therapeutic antisense oligonucleotides

DeepCLIP models produce binding profiles which are directly used for prediction calculations. We wanted to test how the binding profiles reflect *in vivo* sequence-protein binding dynamics and see if the profiles can help locating sites where splice-switching oligonucleotides (SSOs) can be applied for correction of splicing. Thus, we analyzed a known disease-causing mutation (68) in the *ACADM* gene, c.468+7A>G. The mutation is located outside the core U1 snRNP binding motif but is located within a larger GT-rich region, which is extended by the A>G mutation suggesting that it could be generating or strengthening a TDP-43 binding site. We selected wt and mutant sequences 75 bp of length by including the first 37 bp from either side of the locus of the A>G mutation. We then generated a TDP-43 DeepCLIP model based on publicly available binding sites from the POSTAR2 database (48) using the same model parameters as previously (Figure S33) and scored the wt and mutant sequences. We found that DeepCLIP predicts increased binding of TDP-43 to the mutant relative to the wt (Figure 4a).

**Figure 4.**
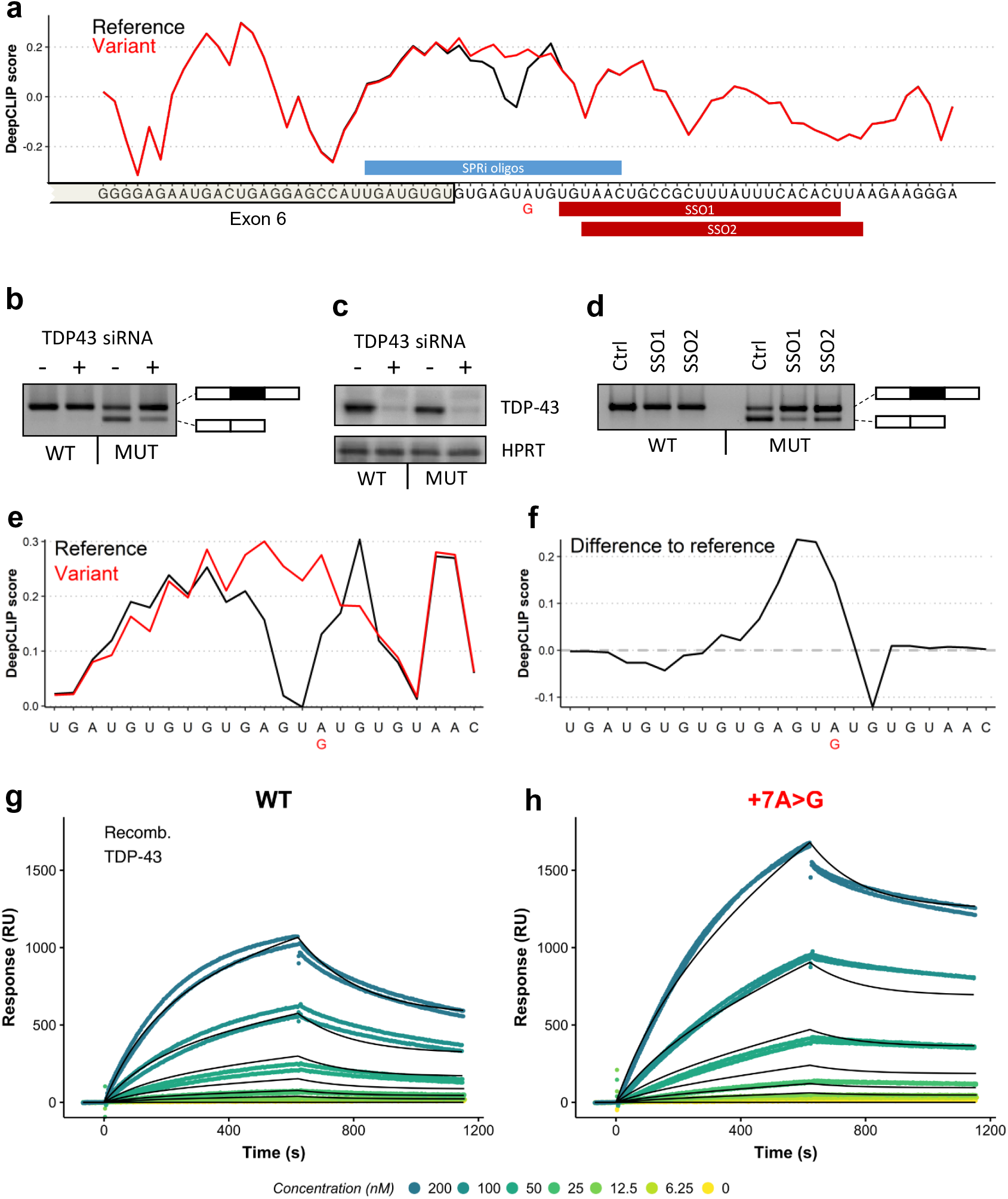
DeepCLIP predicts increased TDP-43 binding as mechanism behind *ACADM* exon 6 skipping. (a) DeepCLIP TDP43 profile across the 5’ss of *ACADM* exon 6 with wt indicated in black and patient mutation indicated in red. Along the first axis the sequence is shown and along the second axis the DeepCLIP BLSTM values are shown. SPRi oligo location and SSO locations are indicated in blue and red bars above and below the sequence, respectively. (b) Splicing of wt and mutant minigenes with either TDP-43 targeting siRNA or non-targeting siRNA determined by RT-PCR. (c) Western blot of TDP-43 and HPRT from siRNA and minigene transfected samples. (d) Splicing of wt and mutant minigenes treated with either a control SSO (Ctrl-SSO), SSO1, or SSO2 determined by RT-PCR. (e) DeepCLIP profile of short RNA oligos used in SPRi measurement, reference in black and +7A>G variant in red. (f) The difference in DeepCLIP binding profiles in (e) between reference and variant. Positive score indicates higher score in variant. (g) SPRi measurements of TDP-43 binding to the wt oligo in (e). (h) SPRi measurements of TDP-43 binding to the variant oligo in (e). In both (g) and (h) the black line indicates the fitted binding model.

Next, we designed a minigene harboring *ACADM* exon 6 and part of the flanking introns to test whether the c.468+7A>G mutation affects splicing of exon 6. We found that the mutation caused dramatic skipping of exon 6 from the minigene (Figure 4b). We hypothesized that this was caused by an increase of TDP-43 binding to the mutant sequence, and that exon skipping therefore could be reversed by treating the cells containing the minigenes with siRNA targeting TDP-43 mRNA. Indeed, TDP-43 siRNA treatment resulted in increased exon inclusion (Figure 4b), corroborating that the c.468+7A>G mutation generates a TDP-43 binding site that causes skipping of *ACADM* exon 6. Splice-switching oligonucleotides (SSOs) are a type of antisense oligonucleotide (ASO) that can be used to modulate splicing by sterically preventing binding of splicing regulatory factors to the RNA. Because of the close proximity of the mutation to the 5’ss, directly blocking the mutant position with an SSO most likely would not result in increased exon inclusion. Interestingly, DeepCLIP finds sites important for binding in a region downstream of the GT-rich core binding motif, which suggests that blocking these sites could prevent TDP-43 binding to the core motif and restore splicing of exon 6.

We tested this hypothesis using two different SSO molecules that targeted this downstream region and which had small overlaps with the end of the GT-rich region (Figure 4d). Strikingly, both SSOs proved very efficacious and almost completely restored splicing from the mutant minigene, indicating that blocking of the downstream motif prevented binding of TDP-43. This indicates that TDP-43 may exhibit context dependent binding modularity, and that the DeepCLIP model is able to detect these context-dependent signatures from the sequence alone.

To validate that TDP-43 binding is directly affected by the mutation, we first analyzed a set of 23 nt oligonucleotides with DeepCLIP (Figure 4e,f) showing that in this shorter context the mutation is still predicted to increase. We then used Surface Plasmon Resonance imaging (SPRi) to measure binding to the 3’ biotin labeled RNA oligonucleotides and observed a pronounced increase in TDP-43 binding to the mutant (Figure 4g-h) in agreement with DeepCLIP predictions.

### DeepCLIP binding scores correlate with *in vitro* binding affinities

One of the most important tasks of a model that predicts presence of RBP binding sites is to accurately estimate the effects of mutations on binding affinity. Therefore, we analyzed 6 sets of wt and mutant exonic variants from the Raponi et al 15-mer set employing SPRi using recombinant hnRNP A1 and SRSF1 as input-proteins (Figure S34-S39, Table S5). These measurements allow quantification of the binding to wt and mutant oligonucleotides, allowing confirmation of DeepCLIP predictions, such as the increase in hnRNP A1 binding to the *PTEN* exon 6 +19C>T mutant (Figure 5a,b). The maximum affinity values obtained by fitting binding models to the measured response by the different SPRi-models correlated well with both hnRNP A1 and SRSF1 DeepCLIP models (Figure 5c-d), across the diverse set of sequences in the dataset (hnRNP A1: Spearman correlation *ρ*=0.874, *p*=0.000309; SRSF1: Spearman correlation *ρ*=0.782, *p*=0.01165). This was also true when we compared DeepCLIP predictions with the Rmax value (Figure S40). This demonstrates that despite being trained on *in vivo* data, the modelling approach of DeepCLIP is also applicable with short *in vitro* sequences, which can be used to examine and validate specific changes in binding to target sites identified by DeepCLIP.

**Figure 5.**
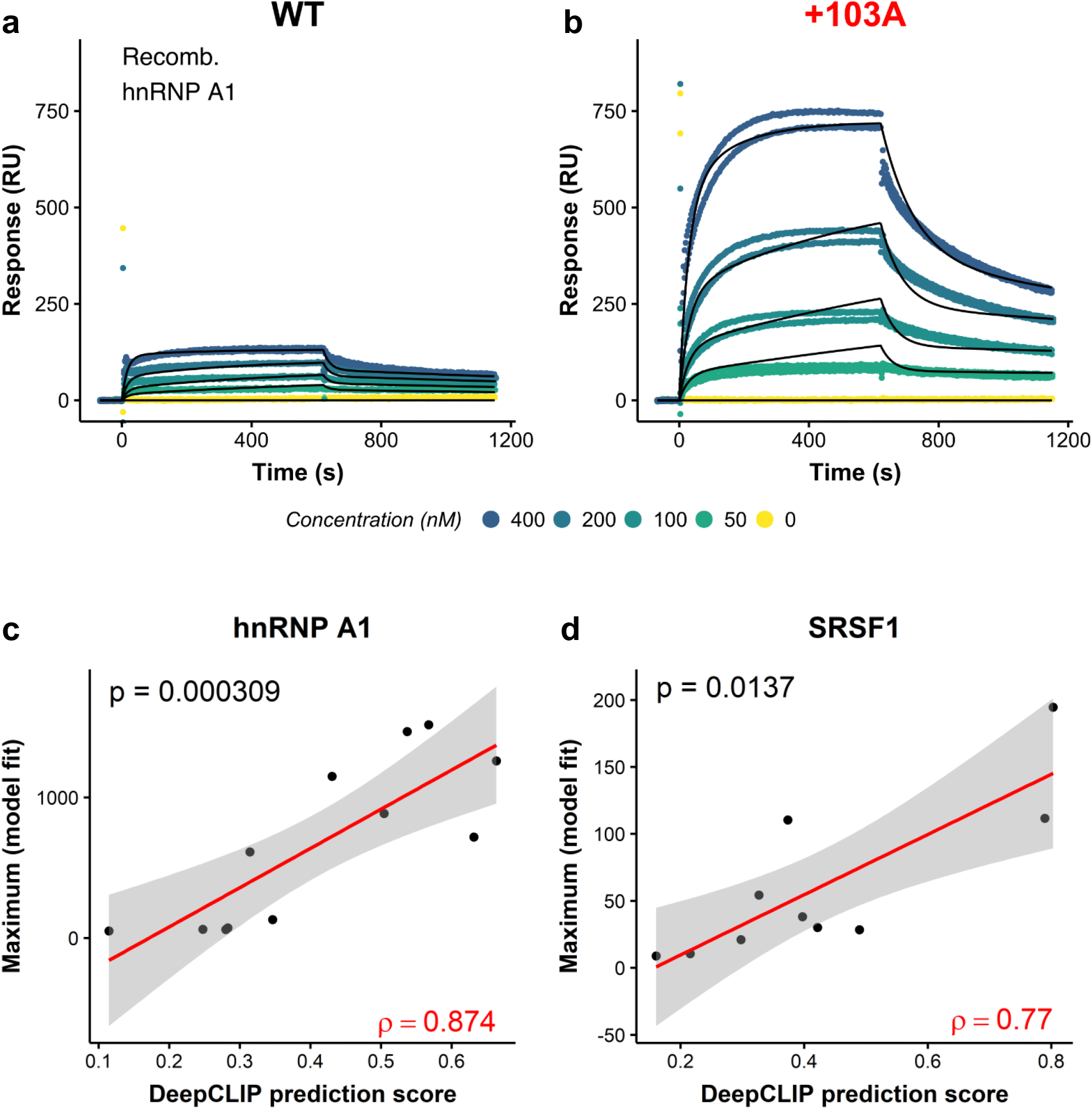
DeepCLIP predictions correlate with binding affinity studies. (a,b) Scatter plots of raw hnRNP A1 SPRi measurements (dots) and the fitted models (black lines) to wt (a) and mutant (b) *ATP7A* exon 20 15 nt oligonucleotide. (c,d) Scatter plots showing DeepCLIP predictions of hnRNP A1 binding (c, n = 12) and SRSF1 binding (d, n = 10) to 15 nt oligonucleotides corresponding to wt and mutant pairs from Raponi et al against the maximum of the binding model fitted to SPRi measurements. The 95% confidence intervals of fitted linear regression models (red line) are shown in grey. Spearman’s rho is show in red in lower right corner, and the p-value in the upper left.

### DeepCLIP analysis of TDP-43-repressed pseudoexons indicates that tissue-specificity is position-dependent

In addition to analyzing sequence variations, DeepCLIP can also be used on a global scale to conduct larger analyses of binding preferences of RBPs. TDP-43 is depleted in the nucleus of motor neurons in patients suffering from amyotrophic lateral sclerosis (ALS) (69, 70). TDP-43 has been reported to repress the inclusion of pseudoexons, and these are then erroneously activated following nuclear depletion, potentially leading to development of ALS symptoms(71). A conditional TDP-43 knock-out mouse-model displays increased pseudoexon inclusion, some of which are muscle and neuron-specific(50). Because these pseudoexons are not necessarily conserved in humans, they may not directly relate to ALS, but they may nevertheless improve our understanding of how some pseudoexons are selectively up-regulated in motor neurons. This can prove important to the understanding of the underlying molecular pathology of ALS. We therefore used DeepCLIP to analyze TDP-43-repressed pseudoexons in mice to examine the tissue specific differences in TDP-43 binding. We found that DeepCLIP overall predicted decreased binding to the region down-stream of the 5’ss of pseudoexons that are neuron specific compared to pseudoexons that are muscle-specific (Figure 6, Figure S41), while neuron-specific pseudoexons were predicted to bind more TDP-43 in the region covering the poly-pyrimidine tract compared to muscle-specific pseudoexons. This might reflect interplay between TDP-43 and tissue-specific factors interacting with these regions in a position-dependent manner. These results indicate that sequence analysis of known pseudoexons can lead to discovery of neuron-specific pseudoexons involved in ALS pathology in humans.

**Figure 6.**
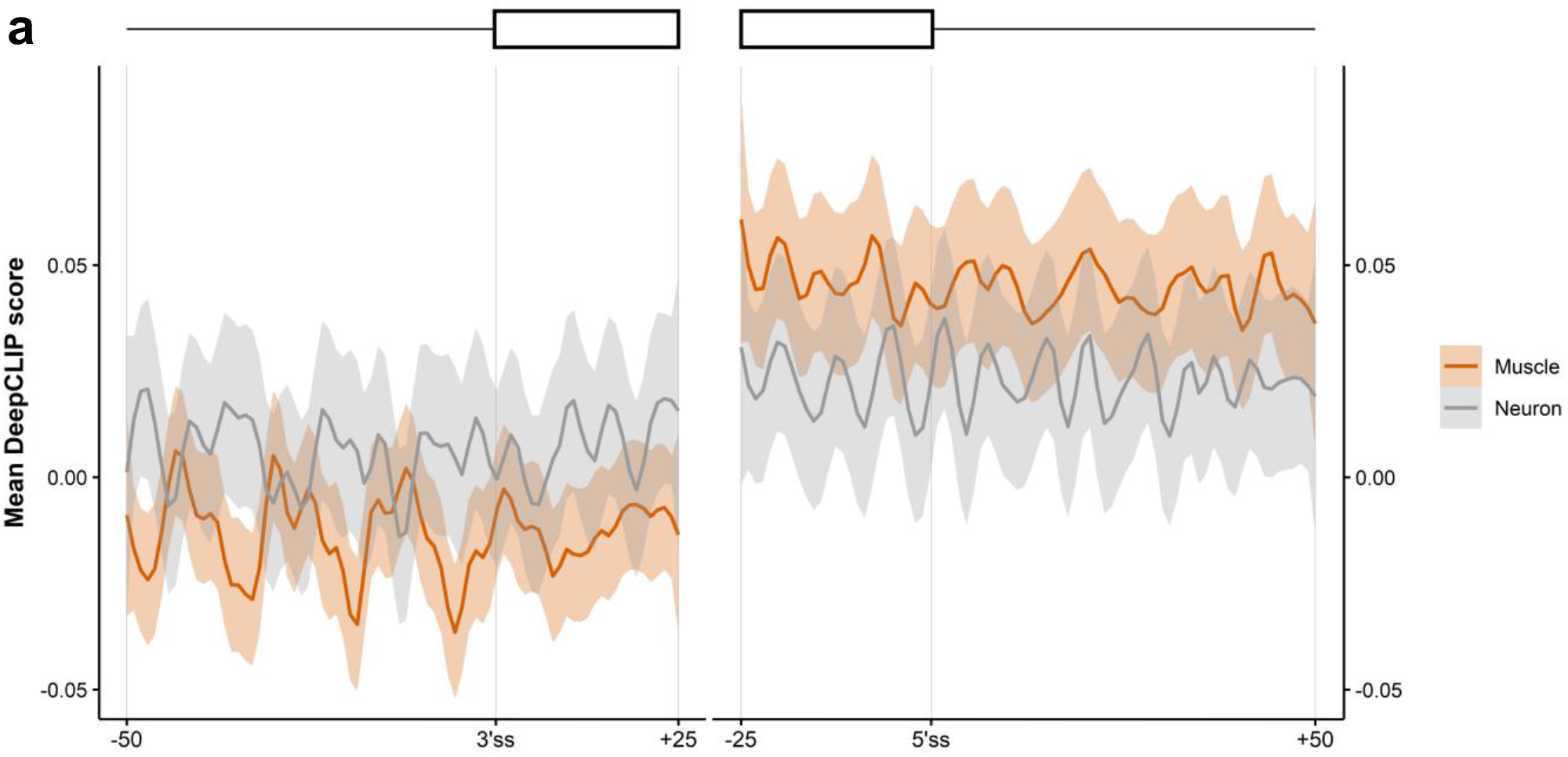
DeepCLIP analysis of TDP-43-repressed pseudoexons indicate position-dependent tissue-specificity. (a) The average DeepCLIP TDP43 profile scores of 58 neuron-specific and 79 muscle-specific pseudoexons activated in TDP43-null mice in the areas covering the 25 first and last nucleotides of the pseudoexon, and the 50 nt spanning intronic regions. 95%-confidence intervals are indicated by shaded areas.

## DISCUSSION

We present DeepCLIP, a novel deep learning approach to modelling RNA-binding protein sites using a shallow neural network composed of CNN and LSTM layers to capture context-dependent binding. DeepCLIP generalizes well across a diverse set of sequences in both *in vitro* and *in vivo* settings, and produces a profile of the sequence, which indicates sequence elements important for the binding of the RNA binding protein in question.

Previous RNA-protein binding classifiers attempted to improve their performance by incorporating context dependencies in a number of different ways, e.g. secondary and tertiary structure, known binding sites of other RNA-binding proteins, and annotated gene regions such as exons, introns, and UTRs.

With DeepCLIP, we demonstrate that a neural network, in which context dependency is not pre-defined, but modelled implicitly by a BLSTM layer, is competitive or outperforming existing classifiers that, in addition to the RNA-sequences, depend on one or more predefined data sets containing different categories of contextual information. This allows DeepCLIP to be agnostic with regard to other inputs and makes it robust towards any limitations in e.g. the modelling of the structure, or the level and quality of annotation of gene structure and other protein binding sites.

Structure modelling was previously shown to improve model accuracy for the datasets Ago1-4, CAPRIN1, IGF2BP1-3, MOV10 and ZC3H7B (14). While DeepCLIP does not directly include predictions of secondary structures when classifying, DeepCLIP AUROC measures for these proteins were the highest of all classifiers except for IGF2BP1-3, where iONMF, which also does not model structure, had a higher AUROC score. This indicates that the BLSTM layer of DeepCLIP captures sufficient structural information, or that other, potentially even hidden and unknown, contextual dependencies are as important as structure modelling.

DeepCLIP produces motifs of varying sizes ranked by the average information content (Figure 2a, b). The top-ranking motifs of DeepCLIP for the analyzed proteins were remarkably similar to core binding sites as described in literature (Figure S4). DeepCLIP does not generate motifs describing structural preferences of RBPs, however this may be obtained by using structure modelling software on high-scoring sites identified by DeepCLIP. DeepCLIP instead integrates contextual information in its models and uses this to output predictions and binding profiles, which indicate binding strength of the total sequence and shows areas of the input-sequence with high and low affinity for the protein in question, respectively.

When searching for binding sites in longer sequences, the information contained in the DeepCLIP binding profiles becomes invaluable, since it unravels interesting areas that are important for protein binding (Figure 2e-f). In the case where a longer sequence contains mainly strong background patterns and only a small segment with binding site potential, DeepCLIP and other tools will be prone to classify this sequence as a background sequence. However, DeepCLIP is able to identify the foreground segment and highlight it on the binding profile of the sequence.

DeepCLIP binding profiles can be used for estimating high- and low-affinity regions of sequences (Figure 4). The predictions, which are directly based on the binding profile values, display a strong correlation with affinity studies (Figure 5) suggesting that DeepCLIP successfully captures binding preferences of RBPs. To this end, the binding profiles produced by DeepCLIP can be used to identify splicing regulatory sites that can be targeted by SSOs (Figure 4a), which is an important novel and missing functionality of existing binding site discovery tools. Thus, DeepCLIP greatly facilitates the design of new drugs based on blocking protein-RNA binding sites, which is a very promising new therapeutic approach, as illustrated for instance by the recent success of the SpinrazaTM SSO in treating SMA (72, 73).

In summary, DeepCLIP models provide valuable insight into the functional consequences of sequence variants. Both *in vitro* binding assays and *in vivo* splicing assays as well as observed splicing of disease-causing mutations in patients cells correlate well with DeepCLIP predictions. This demonstrates that an *in silico* analysis with DeepCLIP can serve as a valuable tool for assessing the functional effects of potentially pathogenic sequence variants, providing an important tool for clinical diagnosis. Finally, we demonstrate that DeepCLIP can serve as tool for designing efficient SSOs for correcting aberrant splicing caused by disease-causing mutations.

## ACKNOWLEDGEMENTS

We are grateful to laboratory technicians Kasper Stonor Werner, Susanne Ruszczycka, and Gabriella Jensen for their assistance with experiments.

## AUTHOR CONTRIBUTIONS

TKD conceived and designed the project. The core algorithm (intrinsic neural network functionalities, layer inter-connectivities and general network composition) was designed and implemented by AGBG. DeepCLIP was implemented by AGBG and TKD. The DeepCLIP web-interface was designed and implemented by SJL. Experiments were designed by TKD, AGBG and BSA and performed by AGBG, USSP, LLH, GHB, MBH, and AMH. All computational analyses were designed by AGBG, TKD, BSA, and JB, and performed by AGBG and TKD. All authors read and contributed to the manuscript.

## DATA AVAILABILITY

All data, including raw data for all figures, in this study is available either through http://github.com/deepclip or http://deepclip.compbio.sdu.dk or direct communication with the authors.

## CODE AVAILABILITY

All code is available at http://github.com/deepclip

