## Supplementary data for "DeepCLIP: Predicting the effect of mutations on protein-RNA binding with Deep Learning"

### SUPPLEMENTARY TABLE LEGENDS

#### Table S1

This table shows the model parameters used when training DeepCLIP models on the curated dataset originally used by Maticzka et al, 2014.

#### Table S2

This table shows performance metrics for the DeepCLIP models trained on the curated dataset originally used by Maticzka et al, 2014.

#### Table S3

This table shows the raw data used in Figure 1 and Figure S2. The highest AUROC score per dataset is indicated in bold.

#### Table S4

This table shows DeepCLIP analysis results of exonic point mutations taken from the dataset previously published by Raponi et al, 2011. During preparation of this manuscript we uncovered some errors in the original dataset, which we corrected. This table contains the raw data for Figure 2c-h.

#### Table S5

This table contains information about the oligos used for SPRi and the model estimates produced by the software applications CLAMP and scrubber, as well as the DeepCLIP prediction scores. This table contains the raw data for Figure 5c-d and Figure S40.

#### SUPPLEMENTARY FIGURE LEGENDS

**Figure S1 | Conversion of the BLSTM output to binding profile and DeepCLIP training methodology.** (a) The output of the forward LSTM layer is shown as a red arrow with a color intensity that increases gradually with the time-step number. The higher the color intensity and the higher the time-step number, the more contextual information has been available to the LSTM layer. The same is shown for the backward LSTM layer (blue arrow). The hidden state produced at the last time-step, , could contain information about the entire sequence. The outputs from the forward and backward LSTM layers are concatenated so hidden-states that are based on the base are combined. This figure is equal to  from the BLSTM layer. Here, the time-step order is equal to the time-step order of the forward LSTM layer. (b) The contextual information that may present in BLSTM hidden-state that represents the highlighted cytosine (C in bold) is shown. The cytosine contains information about all the bases that surrounds it. (c) The binding profile is found via a summation of the vector elements in the BLSTM hidden-states. This results in a new vector that has a length equal to the input sequence. Every element of the vector indicates a class; positive value = class 1 (protein) and negative value = class 0 (genomic background). Every sequence has been divided by the vector element that has the highest absolute value. This results in a sequence that ranges from -1 to 1 and where it is easy to locate areas that are import for protein binding and areas that look like random genomic background. At the bottom, the prediction of the sequences in shown. (d) DeepCLIP is trained on input sequences belonging to either the bound class (assigned a score of 1) or the unbound (background) class (assigned a score of 0). The combined dataset is then divided into a training set, a validation set, and a test set. With default options they constitute 80%, 10%, and 10% of the dataset respectively. (e) When running in 10-fold cross-validation (CV) mode, the training set is divided into 10 different segmentations with a non-overlapping distribution of sequences in the validation and test set. These 10 different segmentations are then used to train 10 different models.

**Figure S2 | Comparison of DeepCLIP AUROC performance on a benchmark dataset with existing tools.** (a) Peak-sequences from the same input sequences used in Maticzka et al., 2014 were used to train DeepCLIP and iDeepS models using 10-fold cross-validation. Area under receiver operating characteristics (AUROC) for each protein were calculated using the combined predictions of all 10 models. AUROC for other methods were based on the reported scores from these studies. mDBN- indicates Deepnet with secondary structures, mDBN+ indicates Deepnet with secondary and tertiary structures. (b) AUROC metrics for the 12 proteins with DeepBind available were obtained by running DeepBind on the full dataset with all available models for the protein and using the model with the highest performance. All other values are identical to those in (a). Because deepnet with tertiary structures performed better than with just secondary structures, we show only this variant in this plot. (c) Boxplot of AUROC scores from each method for the 12 datasets. DeepBind produced an AUROC score of 0.319 for FUS, this datapoint is outside the plotting areas of both (b) and (c) plots.

**Figure S3 | DeepCLIP AUROC performance by CLIP method and dataset size.** (a) AUROC metrics for DeepCLIP grouped into CLIP methods. Significance of pairwise differences was calculated using Wilcoxon rank sum tests. (b) Correlation plot of DeepCLIP AUROC measures and log10 of the number of input binding sites. Correlation was estimated using Spearman’s rank correlation rho. 95% confidence interval of the linear model (red line) is shown in grey.

**Figure S4 | Pseudo-PFMs captured by DeepCLIP.** The column **Protein** contains the names of the analyzed proteins, which have binding preferences as described in the column **Known binding preference**. References to literature can be seen in column **Source**. In the outermost right column, **Pseudo-PFM**, motifs derived from DeepCLIPs convolutional nodes that correlates with known binding preferences are shown. H: A, C or U; N: A, C, G or U.

**Figure S5 | DeepCLIP model characteristics for AGO1-4 GraphProt dataset.** (a) Area under curve analysis of DeepCLIP models trained using 10-fold cross-validation. (b) Visualization of the CNN filters learned by the best performing model based on AUC. Score is equal to the mean information per base. (c-h) Visualizations of the combined model predictions of the 10-fold cross-validation. Scores are scaled to 0-100. (c) Density of background and bound prediction scores. (d) Combined density of prediction scores. (e) Cumulative predictive score of background and bound input sequences. (f) Barplot of cumulative scores of all input sequences. (g) Split prediction scores of background and bound input sequences. (h) Combined AUROC analysis using the pROC R package with DeLong estimation of 95% confidence interval.

**Figure S6 | DeepCLIP model characteristics for AGO2 GraphProt dataset.** (a) Area under curve analysis of DeepCLIP models trained using 10-fold cross-validation. (b) Visualization of the CNN filters learned by the best performing model based on AUC. Score is equal to the mean information per base. (c-h) Visualizations of the combined model predictions of the 10-fold cross-validation. Scores are scaled to 0-100. (c) Density of background and bound prediction scores. (d) Combined density of prediction scores. (e) Cumulative predictive score of background and bound input sequences. (f) Barplot of cumulative scores of all input sequences. (g) Split prediction scores of background and bound input sequences. (h) Combined AUROC analysis using the pROC R package with DeLong estimation of 95% confidence interval.

**Figure S7 | DeepCLIP model characteristics for ALKBH5 GraphProt dataset.** (a) Area under curve analysis of DeepCLIP models trained using 10-fold cross-validation. (b) Visualization of the CNN filters learned by the best performing model based on AUC. Score is equal to the mean information per base. (c-h) Visualizations of the combined model predictions of the 10-fold cross-validation. Scores are scaled to 0-100. (c) Density of background and bound prediction scores. (d) Combined density of prediction scores. (e) Cumulative predictive score of background and bound input sequences. (f) Barplot of cumulative scores of all input sequences. (g) Split prediction scores of background and bound input sequences. (h) Combined AUROC analysis using the pROC R package with DeLong estimation of 95% confidence interval.

**Figure S8 | DeepCLIP model characteristics for C17ORF85 GraphProt dataset.** (a) Area under curve analysis of DeepCLIP models trained using 10-fold cross-validation. (b) Visualization of the CNN filters learned by the best performing model based on AUC. Score is equal to the mean information per base. (c-h) Visualizations of the combined model predictions of the 10-fold cross-validation. Scores are scaled to 0-100. (c) Density of background and bound prediction scores. (d) Combined density of prediction scores. (e) Cumulative predictive score of background and bound input sequences. (f) Barplot of cumulative scores of all input sequences. (g) Split prediction scores of background and bound input sequences. (h) Combined AUROC analysis using the pROC R package with DeLong estimation of 95% confidence interval.

**Figure S9 | DeepCLIP model characteristics for C22ORF28 GraphProt dataset.** (a) Area under curve analysis of DeepCLIP models trained using 10-fold cross-validation. (b) Visualization of the CNN filters learned by the best performing model based on AUC. Score is equal to the mean information per base. (c-h) Visualizations of the combined model predictions of the 10-fold cross-validation. Scores are scaled to 0-100.

(c) Density of background and bound prediction scores. (d) Combined density of prediction scores. (e) Cumulative predictive score of background and bound input sequences. (f) Barplot of cumulative scores of all input sequences. (g) Split prediction scores of background and bound input sequences. (h) Combined AUROC analysis using the pROC R package with DeLong estimation of 95% confidence interval.

**Figure S10 | DeepCLIP model characteristics for CAPRIN1 GraphProt dataset.** (a) Area under curve analysis of DeepCLIP models trained using 10-fold cross-validation. (b) Visualization of the CNN filters learned by the best performing model based on AUC. Score is equal to the mean information per base. (c-h) Visualizations of the combined model predictions of the 10-fold cross-validation. Scores are scaled to 0-100. (c) Density of background and bound prediction scores. (d) Combined density of prediction scores. (e) Cumulative predictive score of background and bound input sequences. (f) Barplot of cumulative scores of all input sequences. (g) Split prediction scores of background and bound input sequences. (h) Combined AUROC analysis using the pROC R package with DeLong estimation of 95% confidence interval.

**Figure S11 | DeepCLIP model characteristics for ELAVL1 GraphProt dataset.** (a) Area under curve analysis of DeepCLIP models trained using 10-fold cross-validation. (b) Visualization of the CNN filters learned by the best performing model based on AUC. Score is equal to the mean information per base. (c-h) Visualizations of the combined model predictions of the 10-fold cross-validation. Scores are scaled to 0-100. (c) Density of background and bound prediction scores. (d) Combined density of prediction scores. (e) Cumulative predictive score of background and bound input sequences. (f) Barplot of cumulative scores of all input sequences. (g) Split prediction scores of background and bound input sequences. (h) Combined AUROC analysis using the pROC R package with DeLong estimation of 95% confidence interval.

**Figure S12 | DeepCLIP model characteristics for ELAVL1A GraphProt dataset.** (a) Area under curve analysis of DeepCLIP models trained using 10-fold cross-validation. (b) Visualization of the CNN filters learned by the best performing model based on AUC. Score is equal to the mean information per base. (c-h) Visualizations of the combined model predictions of the 10-fold cross-validation. Scores are scaled to 0-100. (c) Density of background and bound prediction scores. (d) Combined density of prediction scores. (e) Cumulative predictive score of background and bound input sequences. (f) Barplot of cumulative scores of all input sequences. (g) Split prediction scores of background and bound input sequences. (h) Combined AUROC analysis using the pROC R package with DeLong estimation of 95% confidence interval.

**Figure S13 | DeepCLIP model characteristics for ELAVL1B GraphProt dataset.** (a) Area under curve analysis of DeepCLIP models trained using 10-fold cross-validation. (b) Visualization of the CNN filters learned by the best performing model based on AUC. Score is equal to the mean information per base. (c-h) Visualizations of the combined model predictions of the 10-fold cross-validation. Scores are scaled to 0-100. (c) Density of background and bound prediction scores. (d) Combined density of prediction scores. (e) Cumulative predictive score of background and bound input sequences. (f) Barplot of cumulative scores of all input sequences. (g) Split prediction scores of background and bound input sequences. (h) Combined AUROC analysis using the pROC R package with DeLong estimation of 95% confidence interval.

**Figure S14 | DeepCLIP model characteristics for EWSR1 GraphProt dataset.** (a) Area under curve analysis of DeepCLIP models trained using 10-fold cross-validation. (b) Visualization of the CNN filters learned by the best performing model based on AUC. Score is equal to the mean information per base. (c-h) Visualizations of the combined model predictions of the 10-fold cross-validation. Scores are scaled to 0-100. (c) Density of background and bound prediction scores. (d) Combined density of prediction scores. (e)

Cumulative predictive score of background and bound input sequences. (f) Barplot of cumulative scores of all input sequences. (g) Split prediction scores of background and bound input sequences. (h) Combined AUROC analysis using the pROC R package with DeLong estimation of 95% confidence interval.

**Figure S15 | DeepCLIP model characteristics for FUS GraphProt dataset.** (a) Area under curve analysis of DeepCLIP models trained using 10-fold cross-validation. (b) Visualization of the CNN filters learned by the best performing model based on AUC. Score is equal to the mean information per base. (c-h) Visualizations of the combined model predictions of the 10-fold cross-validation. Scores are scaled to 0-100. (c) Density of background and bound prediction scores. (d) Combined density of prediction scores. (e) Cumulative predictive score of background and bound input sequences. (f) Barplot of cumulative scores of all input sequences. (g) Split prediction scores of background and bound input sequences. (h) Combined AUROC analysis using the pROC R package with DeLong estimation of 95% confidence interval.

**Figure S16 | DeepCLIP model characteristics for hnRNP C GraphProt dataset.** (a) Area under curve analysis of DeepCLIP models trained using 10-fold cross-validation. (b) Visualization of the CNN filters learned by the best performing model based on AUC. Score is equal to the mean information per base. (c-h) Visualizations of the combined model predictions of the 10-fold cross-validation. Scores are scaled to 0-100. (c) Density of background and bound prediction scores. (d) Combined density of prediction scores. (e) Cumulative predictive score of background and bound input sequences. (f) Barplot of cumulative scores of all input sequences. (g) Split prediction scores of background and bound input sequences. (h) Combined AUROC analysis using the pROC R package with DeLong estimation of 95% confidence interval.

**Figure S17 | DeepCLIP model characteristics for HuR GraphProt dataset.** (a) Area under curve analysis of DeepCLIP models trained using 10-fold cross-validation. (b) Visualization of the CNN filters learned by the best performing model based on AUC. Score is equal to the mean information per base. (c-h) Visualizations of the combined model predictions of the 10-fold cross-validation. Scores are scaled to 0-100. (c) Density of background and bound prediction scores. (d) Combined density of prediction scores. (e) Cumulative predictive score of background and bound input sequences. (f) Barplot of cumulative scores of all input sequences. (g) Split prediction scores of background and bound input sequences. (h) Combined AUROC analysis using the pROC R package with DeLong estimation of 95% confidence interval.

**Figure S18 | DeepCLIP model characteristics for IGF2BP1-3 GraphProt dataset.** (a) Area under curve analysis of DeepCLIP models trained using 10-fold cross-validation. (b) Visualization of the CNN filters learned by the best performing model based on AUC. Score is equal to the mean information per base. (c-h) Visualizations of the combined model predictions of the 10-fold cross-validation. Scores are scaled to 0-100. (c) Density of background and bound prediction scores. (d) Combined density of prediction scores. (e) Cumulative predictive score of background and bound input sequences. (f) Barplot of cumulative scores of all input sequences. (g) Split prediction scores of background and bound input sequences. (h) Combined AUROC analysis using the pROC R package with DeLong estimation of 95% confidence interval.

**Figure S19 | DeepCLIP model characteristics for MOV10 GraphProt dataset.** (a) Area under curve analysis of DeepCLIP models trained using 10-fold cross-validation. (b) Visualization of the CNN filters learned by the best performing model based on AUC. Score is equal to the mean information per base. (c-h) Visualizations of the combined model predictions of the 10-fold cross-validation. Scores are scaled to 0-100. (c) Density of background and bound prediction scores. (d) Combined density of prediction scores. (e) Cumulative predictive score of background and bound input sequences. (f) Barplot of cumulative scores of all input sequences. (g) Split prediction scores of background and bound input sequences. (h) Combined AUROC analysis using the pROC R package with DeLong estimation of 95% confidence interval.

**Figure S20 | DeepCLIP model characteristics for PTBP1 GraphProt dataset.** (a) Area under curve analysis of DeepCLIP models trained using 10-fold cross-validation. (b) Visualization of the CNN filters learned by the best performing model based on AUC. Score is equal to the mean information per base. (c-h) Visualizations of the combined model predictions of the 10-fold cross-validation. Scores are scaled to 0-100. (c) Density of background and bound prediction scores. (d) Combined density of prediction scores. (e) Cumulative predictive score of background and bound input sequences. (f) Barplot of cumulative scores of all input sequences. (g) Split prediction scores of background and bound input sequences. (h) Combined AUROC analysis using the pROC R package with DeLong estimation of 95% confidence interval.

**Figure S21 | DeepCLIP model characteristics for PUM2 GraphProt dataset.** (a) Area under curve analysis of DeepCLIP models trained using 10-fold cross-validation. (b) Visualization of the CNN filters learned by the best performing model based on AUC. Score is equal to the mean information per base. (c-h) Visualizations of the combined model predictions of the 10-fold cross-validation. Scores are scaled to 0-100. (c) Density of background and bound prediction scores. (d) Combined density of prediction scores. (e) Cumulative predictive score of background and bound input sequences. (f) Barplot of cumulative scores of all input sequences. (g) Split prediction scores of background and bound input sequences. (h) Combined AUROC analysis using the pROC R package with DeLong estimation of 95% confidence interval.

**Figure S22 | DeepCLIP model characteristics for QKI GraphProt dataset.** (a) Area under curve analysis of DeepCLIP models trained using 10-fold cross-validation. (b) Visualization of the CNN filters learned by the best performing model based on AUC. Score is equal to the mean information per base. (c-h) Visualizations of the combined model predictions of the 10-fold cross-validation. Scores are scaled to 0-100. (c) Density of background and bound prediction scores. (d) Combined density of prediction scores. (e) Cumulative predictive score of background and bound input sequences. (f) Barplot of cumulative scores of all input sequences. (g) Split prediction scores of background and bound input sequences. (h) Combined AUROC analysis using the pROC R package with DeLong estimation of 95% confidence interval.

**Figure S23 | DeepCLIP model characteristics for SRSF1 GraphProt dataset.** (a) Area under curve analysis of DeepCLIP models trained using 10-fold cross-validation. (b) Visualization of the CNN filters learned by the best performing model based on AUC. Score is equal to the mean information per base. (c-h) Visualizations of the combined model predictions of the 10-fold cross-validation. Scores are scaled to 0-100. (c) Density of background and bound prediction scores. (d) Combined density of prediction scores. (e) Cumulative predictive score of background and bound input sequences. (f) Barplot of cumulative scores of all input sequences. (g) Split prediction scores of background and bound input sequences. (h) Combined AUROC analysis using the pROC R package with DeLong estimation of 95% confidence interval.

**Figure S24 | DeepCLIP model characteristics for TAF15 GraphProt dataset.** (a) Area under curve analysis of DeepCLIP models trained using 10-fold cross-validation. (b) Visualization of the CNN filters learned by the best performing model based on AUC. Score is equal to the mean information per base. (c-h) Visualizations of the combined model predictions of the 10-fold cross-validation. Scores are scaled to 0-100. (c) Density of background and bound prediction scores. (d) Combined density of prediction scores. (e) Cumulative predictive score of background and bound input sequences. (f) Barplot of cumulative scores of all input sequences. (g) Split prediction scores of background and bound input sequences. (h) Combined AUROC analysis using the pROC R package with DeLong estimation of 95% confidence interval.

**Figure S25 | DeepCLIP model characteristics for TDP-43 GraphProt dataset.** (a) Area under curve analysis of DeepCLIP models trained using 10-fold cross-validation. (b) Visualization of the CNN filters learned by the best performing model based on AUC. Score is equal to the mean information per base. (c-h) Visualizations of the combined model predictions of the 10-fold cross-validation. Scores are scaled to 0-100. (c) Density of background and bound prediction scores. (d) Combined density of prediction scores. (e) Cumulative predictive score of background and bound input sequences. (f) Barplot of cumulative scores of all input sequences. (g) Split prediction scores of background and bound input sequences. (h) Combined AUROC analysis using the pROC R package with DeLong estimation of 95% confidence interval.

**Figure S26 | DeepCLIP model characteristics for TIA1 GraphProt dataset.** (a) Area under curve analysis of DeepCLIP models trained using 10-fold cross-validation. (b) Visualization of the CNN filters learned by the best performing model based on AUC. Score is equal to the mean information per base. (c-h) Visualizations of the combined model predictions of the 10-fold cross-validation. Scores are scaled to 0-100. (c) Density of background and bound prediction scores. (d) Combined density of prediction scores. (e) Cumulative predictive score of background and bound input sequences. (f) Barplot of cumulative scores of all input sequences. (g) Split prediction scores of background and bound input sequences. (h) Combined AUROC analysis using the pROC R package with DeLong estimation of 95% confidence interval.

**Figure S27 | DeepCLIP model characteristics for TIAL1 GraphProt dataset.** (a) Area under curve analysis of DeepCLIP models trained using 10-fold cross-validation. (b) Visualization of the CNN filters learned by the best performing model based on AUC. Score is equal to the mean information per base. (c-h) Visualizations of the combined model predictions of the 10-fold cross-validation. Scores are scaled to 0-100. (c) Density of background and bound prediction scores. (d) Combined density of prediction scores. (e) Cumulative predictive score of background and bound input sequences. (f) Barplot of cumulative scores of all input sequences. (g) Split prediction scores of background and bound input sequences. (h) Combined AUROC analysis using the pROC R package with DeLong estimation of 95% confidence interval.

**Figure S28 | DeepCLIP model characteristics for ZC3H7B GraphProt dataset.** (a) Area under curve analysis of DeepCLIP models trained using 10-fold cross-validation. (b) Visualization of the CNN filters learned by the best performing model based on AUC. Score is equal to the mean information per base. (c-h) Visualizations of the combined model predictions of the 10-fold cross-validation. Scores are scaled to 0-100. (c) Density of background and bound prediction scores. (d) Combined density of prediction scores. (e) Cumulative predictive score of background and bound input sequences. (f) Barplot of cumulative scores of all input sequences. (g) Split prediction scores of background and bound input sequences. (h) Combined AUROC analysis using the pROC R package with DeLong estimation of 95% confidence interval.

**Figure S29 | DeepCLIP model characteristics for hnRNP A1.** (a) Area under curve analysis of DeepCLIP models trained on hnRNP A1 iCLIP data from Bruun et al. using 10-fold cross-validation. (b) Visualization of the CNN filters learned by the best performing model based on AUC. Score is equal to the mean information per base. (c-h) Visualizations of the combined model predictions of the 10-fold cross-validation. Scores are scaled to 0-100. (c) Density of background and bound prediction scores. (d) Combined density of prediction scores. (e) Cumulative predictive score of background and bound input sequences. (f) Barplot of cumulative scores of all input sequences. (g) Split prediction scores of background and bound input sequences. (h) Combined AUROC analysis using the pROC R package with DeLong estimation of 95% confidence interval.

**Figure S30 | DeepCLIP model characteristics for SRSF1 (eCLIP).** (a) Area under curve analysis of DeepCLIP models trained on SRSF1 eCLIP data from Van Nostrand et al using 10-fold cross-validation. (b) Visualization of the CNN filters learned by the best performing model based on AUC. Score is equal to the mean information per base. (c-h) Visualizations of the combined model predictions of the 10-fold cross-validation. Scores are scaled to 0-100. (c) Density of background and bound prediction scores. (d) Combined density of prediction scores. (e) Cumulative predictive score of background and bound input sequences. (f) Barplot of cumulative scores of all input sequences. (g) Split prediction scores of background and bound input sequences. (h) Combined AUROC analysis using the pROC R package with DeLong estimation of 95% confidence interval.

**Figure S31 | DeepCLIP predictions of *ACADM* exon 5 mutations using SRSF1 (HITS-CLIP) model.** (a) Scatter plot of *ACADM* exon 5 minigene percent spliced in (PSI) values and DeepCLIP SRSF1 score with linear regression (red line) and 95% confidence interval (shaded area). (b) Same as (a) but with the DeepCLIP hnRNP A1 score subtracted from the SRSF1 score. Spearman’s rho is indicated in red for both plots in (a-b). (c) Barplot of the change relative to the wt of exon inclusion levels (blue), hnRNP A1 DeepCLIP scores (brown) and SRSF1 HITS-CLIP DeepCLIP model scores (green). The hnRNP A1 results are the same as presented in figure 5 but are included here for reference.

**Figure S32 | Correlation between observed exon skipping and exon skipping model predictions.** (a) Scatter plot of exon inclusion levels vs EX-SKIP predictions. (b) Scatter plot of the exon inclusions levels vs the SPANR prediction exon inclusion levels. Spearman’s rho is indicated in red for both plots in (a-b).

**Figure S33 | DeepCLIP model characteristics for TDP-43.** (a) Area under curve analysis of DeepCLIP models trained on TDP43 binding sites from the POSTAR2 database using 10-fold cross-validation. (b) Visualization of the CNN filters learned by the best performing model based on AUC. Score is equal to the mean information per base. (c-h) Visualizations of the combined model predictions of the 10-fold cross-validation. Scores are scaled to 0-100. (c) Density of background and bound prediction scores. (d) Combined density of prediction scores. (e) Cumulative predictive score of background and bound input sequences. (f) Barplot of cumulative scores of all input sequences. (g) Split prediction scores of background and bound input sequences. (h) Combined AUROC analysis using the pROC R package with DeLong estimation of 95% confidence interval.

**Figure S34-36 | Results of SPRi measurements of hnRNP A1 binding to a set of RNA wt and mutant oligos.** Plots showing measured response (RU) versus time of the wt oligo (left) and mutant (right). The combined model fit across concentrations is indicated in black and different concentrations are indicated green, brown, purple, pink, and olive in decreasing order. The fitted model’s maximum simulated value is indicated above each plot.

**Figure S37-39 | Results of SPRi measurements of SRSF1 binding to a set of RNA wt and mutant oligos.** Plots showing measured response (RU) versus time of the wt oligo (left) and mutant (right). The combined model fit across concentrations is indicated in black and different concentrations are indicated green, brown, purple, pink, and olive in decreasing order. The fitted model's maximum simulated value is indicated above each plot.

**Figure S40 | Correlation between observed *in vitro* binding and DeepCLIP hnRNP A1 and SRSF1 (GraphProt dataset) model predictions.** (a) Scatter plot of Rmax binding values from SPRi measurements of hnRNP A1 binding to oligos vs the predicted DeepCLIP hnRNP A1 scores. (b) Scatter plot of Rmax binding values from SPRi measurements of SRSF1 binding to oligos vs the predicted DeepCLIP SRSF1 scores. Spearman's rho is indicated in red in lower right corner and p-value in black in upper left corner for both plots in (a-b). The data for hnRNP A1 in this plot is the same as in Figure 3 and is included here for comparison.

**Figure S41 | DeepCLIP predicted binding at TDP-43 repressed pseudoexons.** (a-c) Heatmap of TDP-43 DeepCLIP binding profiles at the 3'ss and 5'ss regions of TDP-43 repressed pseudoexons in mice. In all heatmaps the plots show 25 nt into the exon indicated by blue bars at the top, and the 50 first and last nt of the introns. The heatmap in (a) shows pseudoexons up-relugated upon TDP-43 depletion in at least two of the examined tissues (stem cells, neurons, and muscle cells), while the heatmap in (b) show muscle specific pseudoexons and the heatmap in (c) shows neuron specific pseudoexons.

Fig. S1

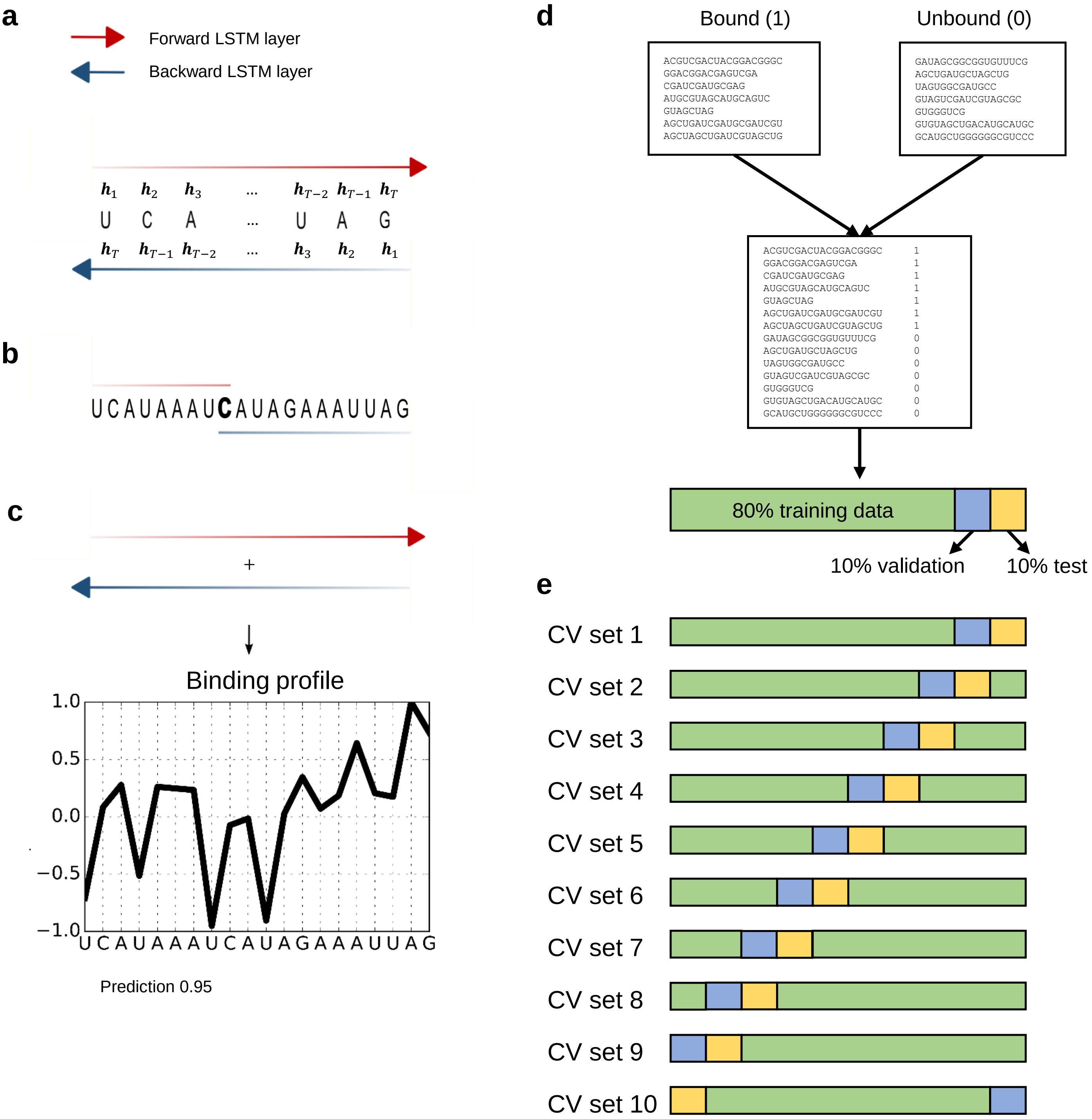
a
Forward LSTM layer
Backward LSTM layer
$h_1$ $h_2$ $h_3$ ... $h_{T-2}$ $h_{T-1}$ $h_T$
U C A ... U A G
$h_T$ $h_{T-1}$ $h_{T-2}$ ... $h_3$ $h_2$ $h_1$
b
UCAUAAAUCAUAGAAAUUAG
c
+
Binding profile
1.0
0.5
0.0
−0.5
−1.0
U C A U A A A U C A U A G A A A U U A G
Prediction 0.95
d
Bound (1)
Unbound (0)
ACGUCGACUACGGACGGGC
GGACGGACGAGUCGA
CGAUCGAUGCGAG
AUGCGUAGCAUGCAGUC
GUAGCUAG
AGCUGAUCGAUGCGAUCGU
AGCUAGCUGAUCGUAGCUG
GAUAGCGGCGGUGUUUCG
AGCUGAUGCUAGCUG
UAGUGGCGAUGCC
GUAGUCGAUCGUAGCGC
GUGGGUCG
GUGUAGCUGACAUGCAUGC
GCAUGCUGGGGGGCGUCCC
ACGUCGACUACGGACGGGC 1
GGACGGACGAGUCGA 1
CGAUCGAUGCGAG 1
AUGCGUAGCAUGCAGUC 1
GUAGCUAG 1
AGCUGAUCGAUGCGAUCGU 1
AGCUAGCUGAUCGUAGCUG 1
GAUAGCGGCGGUGUUUCG 0
AGCUGAUGCUAGCUG 0
UAGUGGCGAUGCC 0
GUAGUCGAUCGUAGCGC 0
GUGGGUCG 0
GUGUAGCUGACAUGCAUGC 0
GCAUGCUGGGGGGCGUCCC 0
80% training data
10% validation
10% test
e
CV set 1
CV set 2
CV set 3
CV set 4
CV set 5
CV set 6
CV set 7
CV set 8
CV set 9
CV set 10

Fig. S2

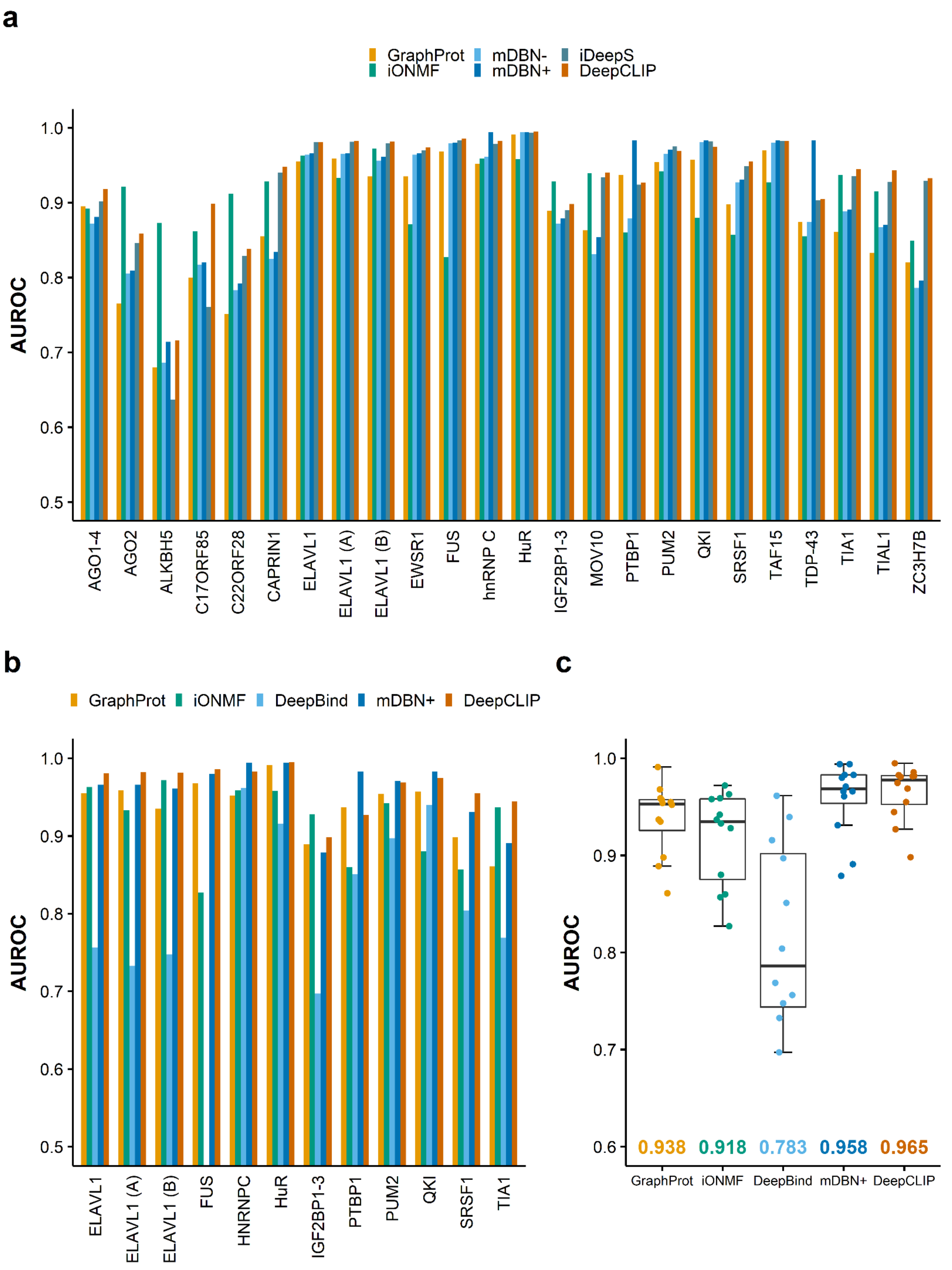

a
GraphProt
mDBN-
iDeepS
iONMF
mDBN+
DeepCLIP
AUROC
1.0
0.9
0.8
0.7
0.6
0.5
AGO1-4
AGO2
ALKBH5
C17ORF85
C22ORF28
CAPRIN1
ELAVL1
ELAVL1 (A)
ELAVL1 (B)
EWSR1
FUS
hnRNP C
HuR
IGF2BP1-3
MOV10
PTBP1
PUM2
QKI
SRSF1
TAF15
TDP-43
TIA1
TIAL1
ZC3H7B
b
GraphProt
iONMF
DeepBind
mDBN+
DeepCLIP
AUROC
ELAVL1
ELAVL1 (A)
ELAVL1 (B)
FUS
HNRNPC
HuR
IGF2BP1-3
PTBP1
PUM2
QKI
SRSF1
TIA1
c
AUROC
0.938
0.918
0.783
0.958
0.965
GraphProt
iONMF
DeepBind
mDBN+
DeepCLIP

Fig. S3

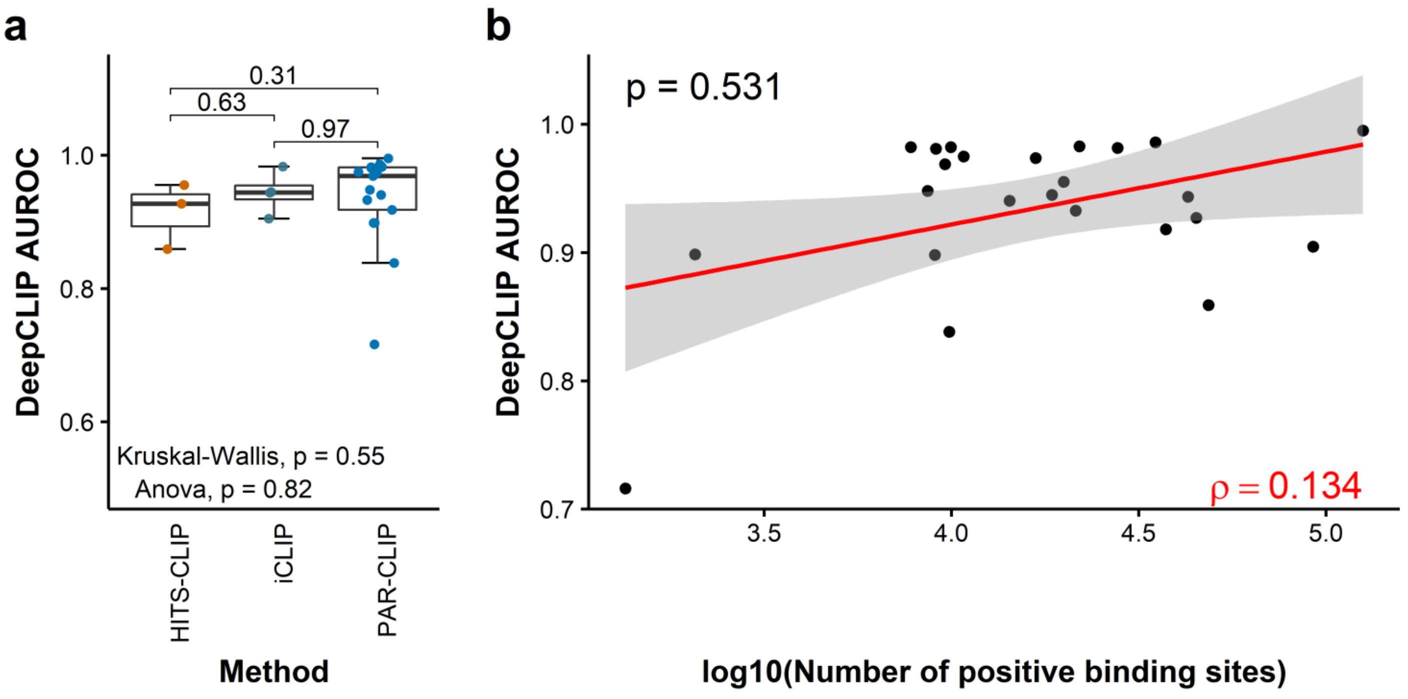
a
0.31
0.63
0.97
1.0
0.8
0.6
DeepCLIP AUROC
Kruskal-Wallis, p = 0.55
Anova, p = 0.82
HITS-CLIP
iCLIP
PAR-CLIP
Method
b
p = 0.531
1.0
0.9
0.8
0.7
DeepCLIP AUROC
ρ = 0.134
3.5
4.0
4.5
5.0
log10(Number of positive binding sites)

### Fig. S4

| Protein (dataset) | Known binding preference | Add. binding preference source | CLIP data source | CLIP method | Top-2 CNN filters |
|---|---|---|---|---|---|
| AGO1-4 | Mainly binds coding regions and 3'UTR | - | Hafner et al., 2010 | PAR-CLIP | 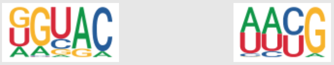 |
| AGO2 | Mainly binds coding regions and 3'UTR | - | Kishore et al., 2011 | HITS-CLIP | 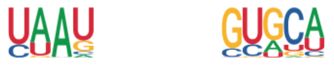 |
| ALKBH5 | Mainly binds coding regions and 3'UTR | - | Baltz et al., 2012 | PAR-CLIP | 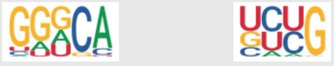 |
| C17ORF85 | Mainly binds coding regions | - | Baltz et al., 2012 | PAR-CLIP | 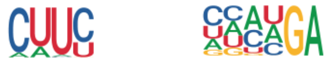 |
| C22ORF28 | Mainly binds coding regions | - | Baltz et al., 2012 | PAR-CLIP | 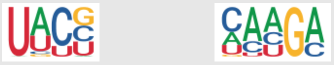 |
| CAPRIN1 | Mainly binds coding regions and 3'UTR | - | Baltz et al., 2012 | PAR-CLIP | 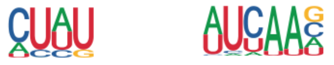 |
| ELAVL1 | Binds single-stranded AU or U-rich regions | Barker et al., 2012 | Kishore et al., 2011 | HITS-CLIP | 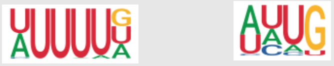 |
| ELAVL1 (A) | Binds single-stranded AU or U-rich regions | Barker et al., 2012 | Kishore et al., 2011 | PAR-CLIP | 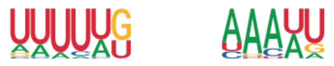 |
| ELAVL1 (B) | Binds single-stranded AU or U-rich regions | Barker et al., 2012 | Lebedeva et al., 2011 | PAR-CLIP | 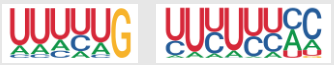 |
| EWSR1 | Binds AU-rich loop structures | - | Hoell et al., 2011 | PAR-CLIP | 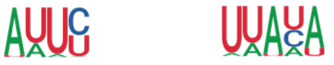 |
| FUS | Binds AU-rich loop structures | - | Hoell et al., 2011 | PAR-CLIP | 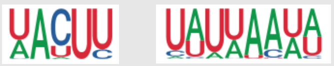 |
| hnRNP C | Binds U-rich regions | - | König et al., 2010 | iCLIP | 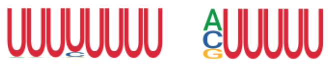 |
| HuR | Binds single-stranded AU or U-rich regions | Barker et al., 2012 | Mukherjee et al., 2011 | PAR-CLIP | 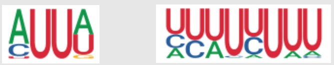 |
| IGF2BP1-3 | Binds strongly to CAUH-patterns | - | Hafner et al., 2010 | PAR-CLIP | 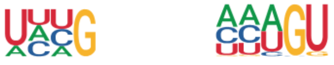 |
| MOV10 | Binds AU-rich 3’ UTR regions | Gregersen et al., 2014 | Sievers et al., 2012 | PAR-CLIP | 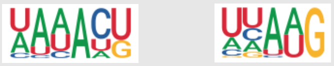 |
| PTBP1 | Binds CU-rich regions | Perez et al., 1997 | Xue et al., 2009 | HITS-CLIP | 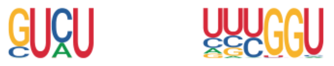 |
| PUM2 | High-affinity binding sites contain UGUANAU | - | Hafner et al., 2010 | PAR-CLIP | 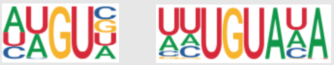 |
| QKI | High-affinity binding sites contain YUAAY | - | Hafner et al., 2010 | PAR-CLIP | 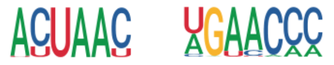 |
| SRSF1 | Single-stranded AGAAGAAG motif | Wang et al., 2011 | Sanford et al., 2009 | HITS-CLIP | 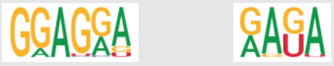 |
| TAF15 | Binds AU-rich loop structures | - | Hoell et al., 2011 | PAR-CLIP | 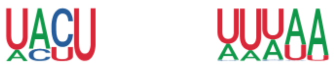 |
| TDP43 | Binds GU-rich regions | Buratti et al., 2004 | Tollervey et al., 2011 | iCLIP | 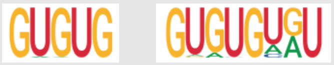 |
| TIA1 | Binds U-rich regions near 5’ss | Dember et al., 1996 | Wang et al., 2010 | iCLIP | 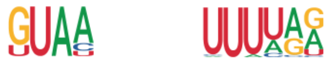 |
| TIAL1 | Binds U-rich regions near 5’ss | Dember et al., 1996 | Wang et al., 2010 | iCLIP | 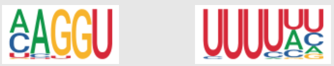 |
| ZC3H7B | Mainly binds coding regions, 3'UTR and intronic regions | - | Baltz et al., 2012 | PAR-CLIP | 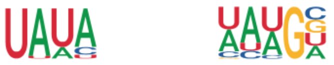 |

Fig. S5

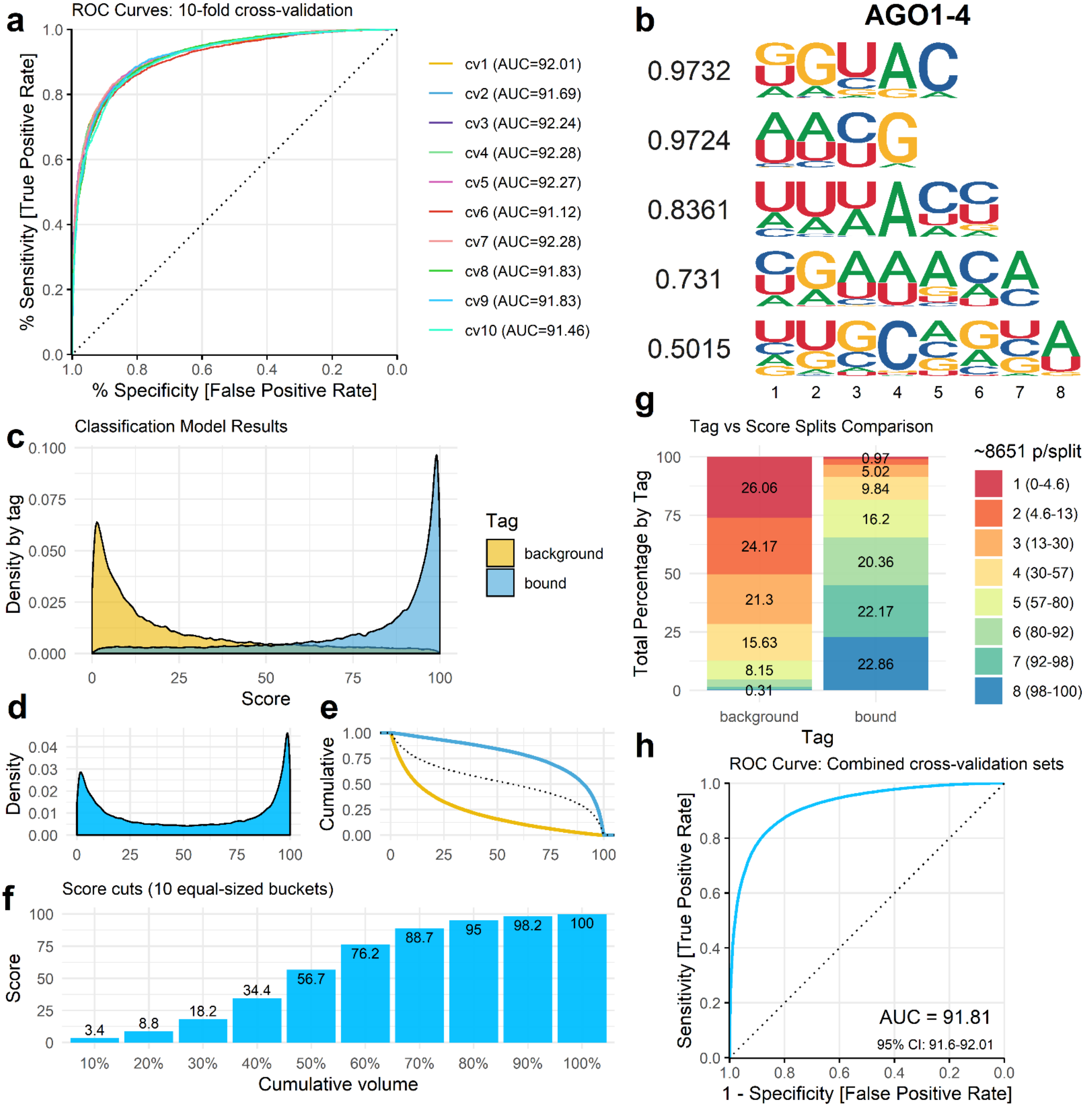
a
ROC Curves: 10-fold cross-validation
% Sensitivity [True Positive Rate]
% Specificity [False Positive Rate]
cv1 (AUC=92.01)
cv2 (AUC=91.69)
cv3 (AUC=92.24)
cv4 (AUC=92.28)
cv5 (AUC=92.27)
cv6 (AUC=91.12)
cv7 (AUC=92.28)
cv8 (AUC=91.83)
cv9 (AUC=91.83)
cv10 (AUC=91.46)
b
AGO1-4
0.9732
0.9724
0.8361
0.731
0.5015
c
Classification Model Results
Density by tag
Score
Tag
background
bound
d
Density
e
Cumulative
f
Score cuts (10 equal-sized buckets)
Score
Cumulative volume
3.4
8.8
18.2
34.4
56.7
76.2
88.7
95
98.2
100
g
Tag vs Score Splits Comparison
Total Percentage by Tag
~8651 p/split
1 (0-4.6)
2 (4.6-13)
3 (13-30)
4 (30-57)
5 (57-80)
6 (80-92)
7 (92-98)
8 (98-100)
26.06
24.17
21.3
15.63
8.15
0.31
0.97
5.02
9.84
16.2
20.36
22.17
22.86
background
bound
Tag
h
ROC Curve: Combined cross-validation sets
Sensitivity [True Positive Rate]
1 - Specificity [False Positive Rate]
AUC = 91.81
95% CI: 91.6-92.01

Fig. S6

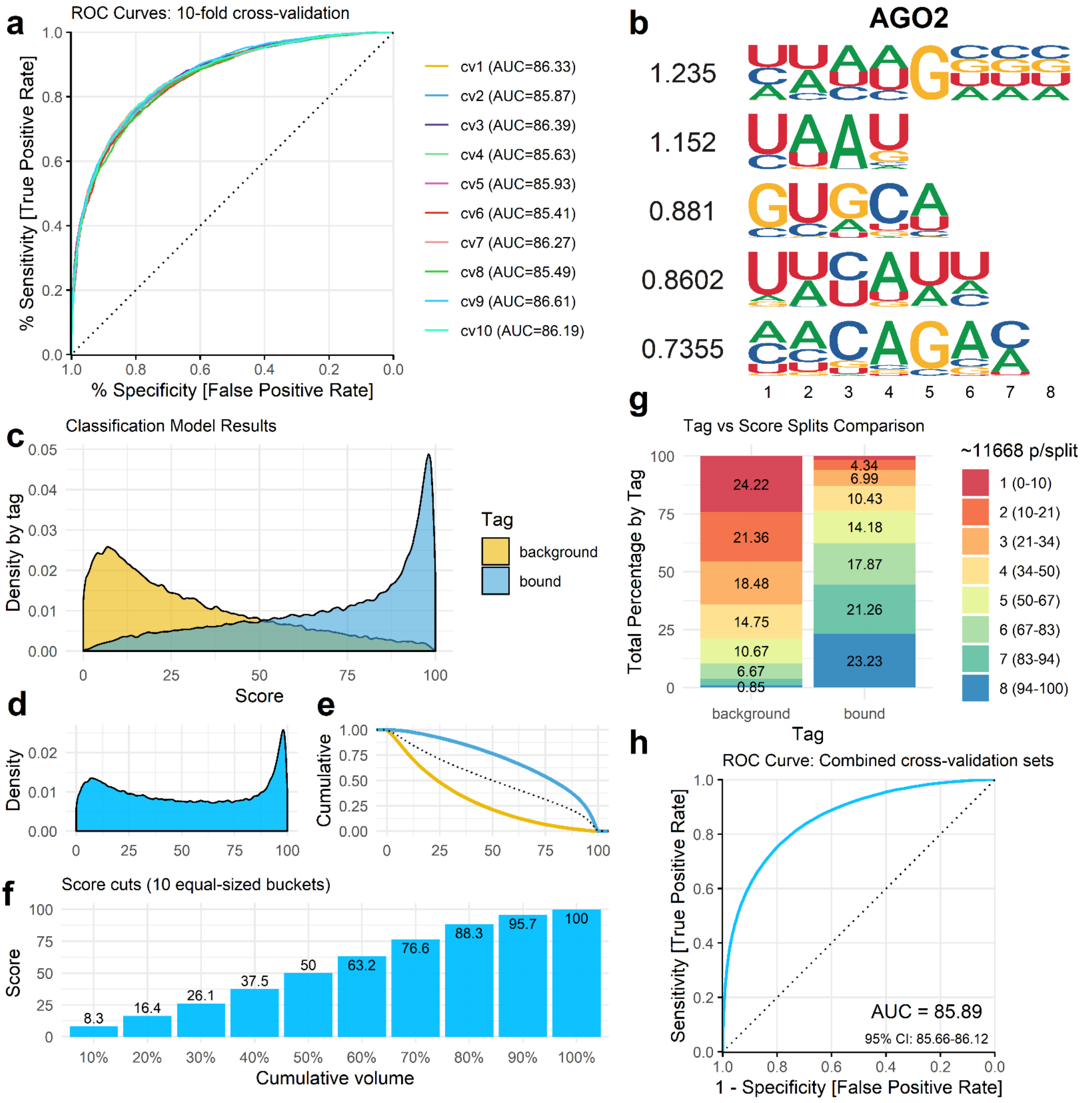
a
ROC Curves: 10-fold cross-validation
% Sensitivity [True Positive Rate]
% Specificity [False Positive Rate]
cv1 (AUC=86.33)
cv2 (AUC=85.87)
cv3 (AUC=86.39)
cv4 (AUC=85.63)
cv5 (AUC=85.93)
cv6 (AUC=85.41)
cv7 (AUC=86.27)
cv8 (AUC=85.49)
cv9 (AUC=86.61)
cv10 (AUC=86.19)
b
AGO2
1.235
1.152
0.881
0.8602
0.7355
c
Classification Model Results
Density by tag
Score
Tag
background
bound
d
Density
e
Cumulative
f
Score cuts (10 equal-sized buckets)
Score
Cumulative volume
8.3
16.4
26.1
37.5
50
63.2
76.6
88.3
95.7
100
g
Tag vs Score Splits Comparison
Total Percentage by Tag
24.22
21.36
18.48
14.75
10.67
6.67
0.85
4.34
6.99
10.43
14.18
17.87
21.26
23.23
background
bound
Tag
~11668 p/split
1 (0-10)
2 (10-21)
3 (21-34)
4 (34-50)
5 (50-67)
6 (67-83)
7 (83-94)
8 (94-100)
h
ROC Curve: Combined cross-validation sets
Sensitivity [True Positive Rate]
1 - Specificity [False Positive Rate]
AUC = 85.89
95% CI: 85.66-86.12

Fig. S7

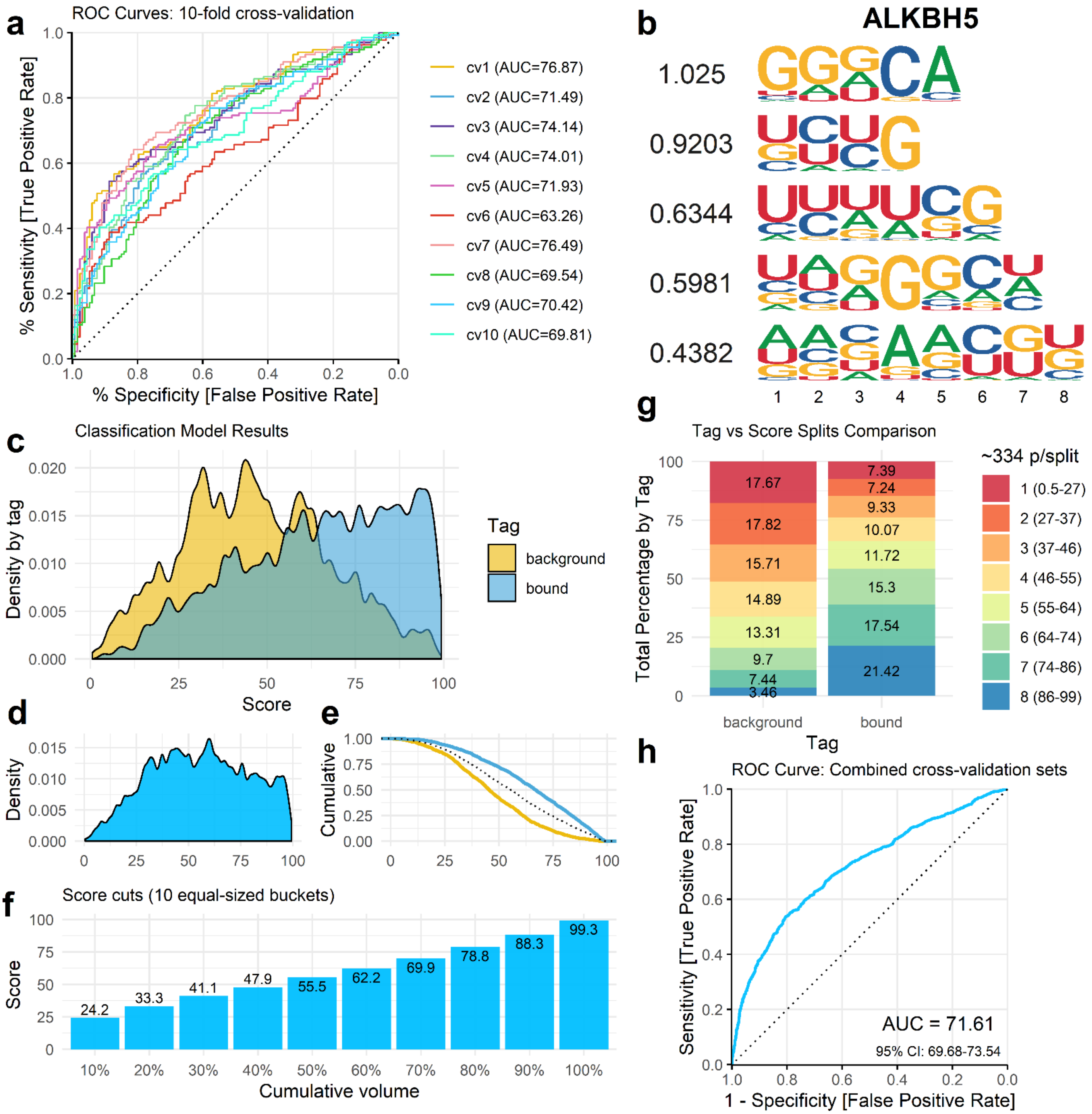
a
ROC Curves: 10-fold cross-validation
% Sensitivity [True Positive Rate]
% Specificity [False Positive Rate]
cv1 (AUC=76.87)
cv2 (AUC=71.49)
cv3 (AUC=74.14)
cv4 (AUC=74.01)
cv5 (AUC=71.93)
cv6 (AUC=63.26)
cv7 (AUC=76.49)
cv8 (AUC=69.54)
cv9 (AUC=70.42)
cv10 (AUC=69.81)
b
ALKBH5
1.025
0.9203
0.6344
0.5981
0.4382
c
Classification Model Results
Density by tag
Score
Tag
background
bound
d
Density
e
Cumulative
f
Score cuts (10 equal-sized buckets)
Score
Cumulative volume
24.2
33.3
41.1
47.9
55.5
62.2
69.9
78.8
88.3
99.3
g
Tag vs Score Splits Comparison
Total Percentage by Tag
~334 p/split
1 (0.5-27)
2 (27-37)
3 (37-46)
4 (46-55)
5 (55-64)
6 (64-74)
7 (74-86)
8 (86-99)
background: 17.67, 17.82, 15.71, 14.89, 13.31, 9.7, 7.44, 3.46
bound: 7.39, 7.24, 9.33, 10.07, 11.72, 15.3, 17.54, 21.42
Tag
h
ROC Curve: Combined cross-validation sets
Sensitivity [True Positive Rate]
1 - Specificity [False Positive Rate]
AUC = 71.61
95% CI: 69.68-73.54

Fig. S8

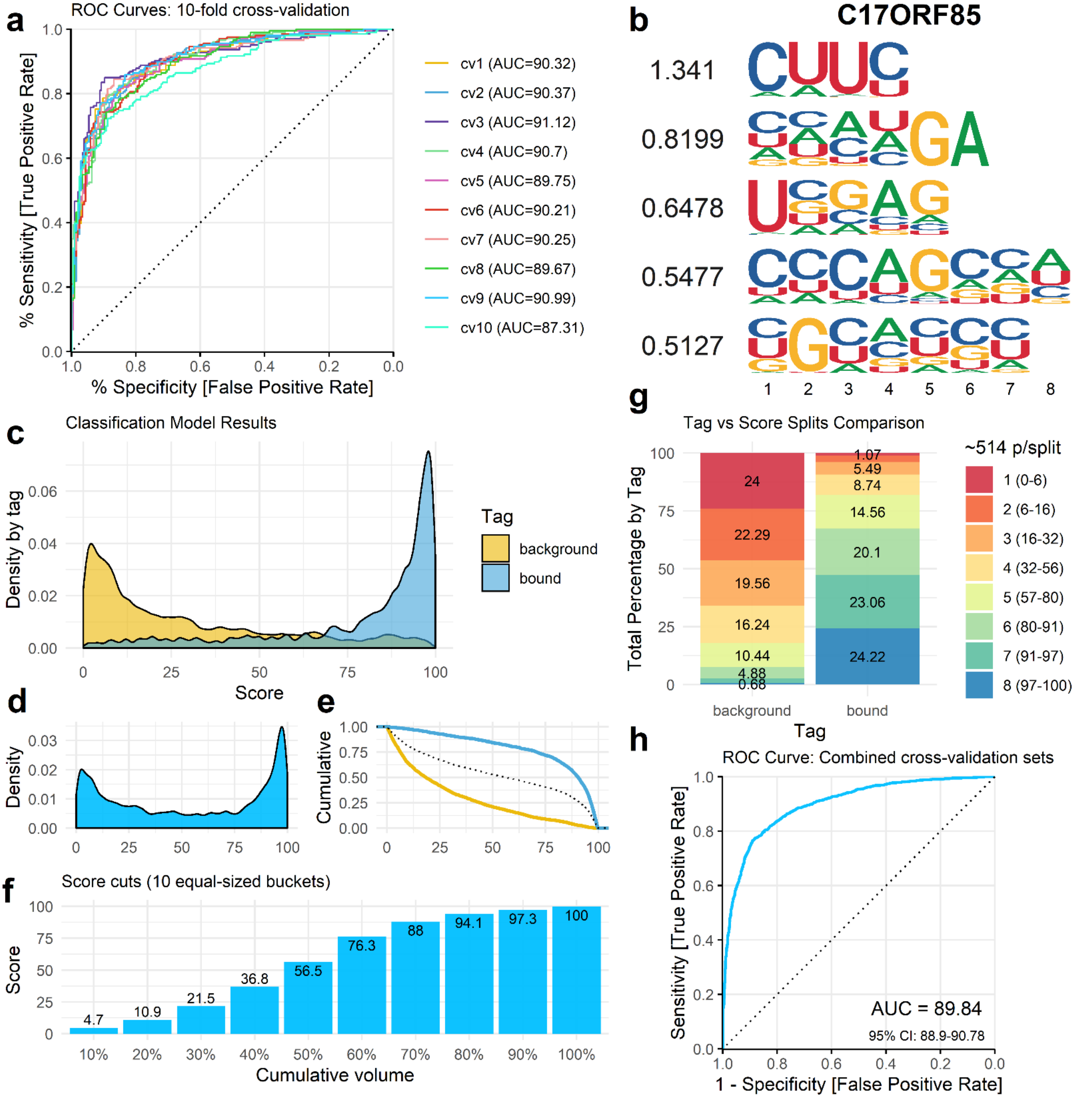
a
ROC Curves: 10-fold cross-validation
% Sensitivity [True Positive Rate]
% Specificity [False Positive Rate]
cv1 (AUC=90.32)
cv2 (AUC=90.37)
cv3 (AUC=91.12)
cv4 (AUC=90.7)
cv5 (AUC=89.75)
cv6 (AUC=90.21)
cv7 (AUC=90.25)
cv8 (AUC=89.67)
cv9 (AUC=90.99)
cv10 (AUC=87.31)
b
C17ORF85
1.341
0.8199
0.6478
0.5477
0.5127
c
Classification Model Results
Density by tag
Score
Tag
background
bound
d
Density
e
Cumulative
f
Score cuts (10 equal-sized buckets)
Score
Cumulative volume
4.7
10.9
21.5
36.8
56.5
76.3
88
94.1
97.3
100
g
Tag vs Score Splits Comparison
Total Percentage by Tag
Tag
~514 p/split
1 (0-6)
2 (6-16)
3 (16-32)
4 (32-56)
5 (57-80)
6 (80-91)
7 (91-97)
8 (97-100)
24
22.29
19.56
16.24
10.44
4.88
0.68
1.07
5.49
8.74
14.56
20.1
23.06
24.22
h
ROC Curve: Combined cross-validation sets
Sensitivity [True Positive Rate]
1 - Specificity [False Positive Rate]
AUC = 89.84
95% CI: 88.9-90.78

### Fig. S9

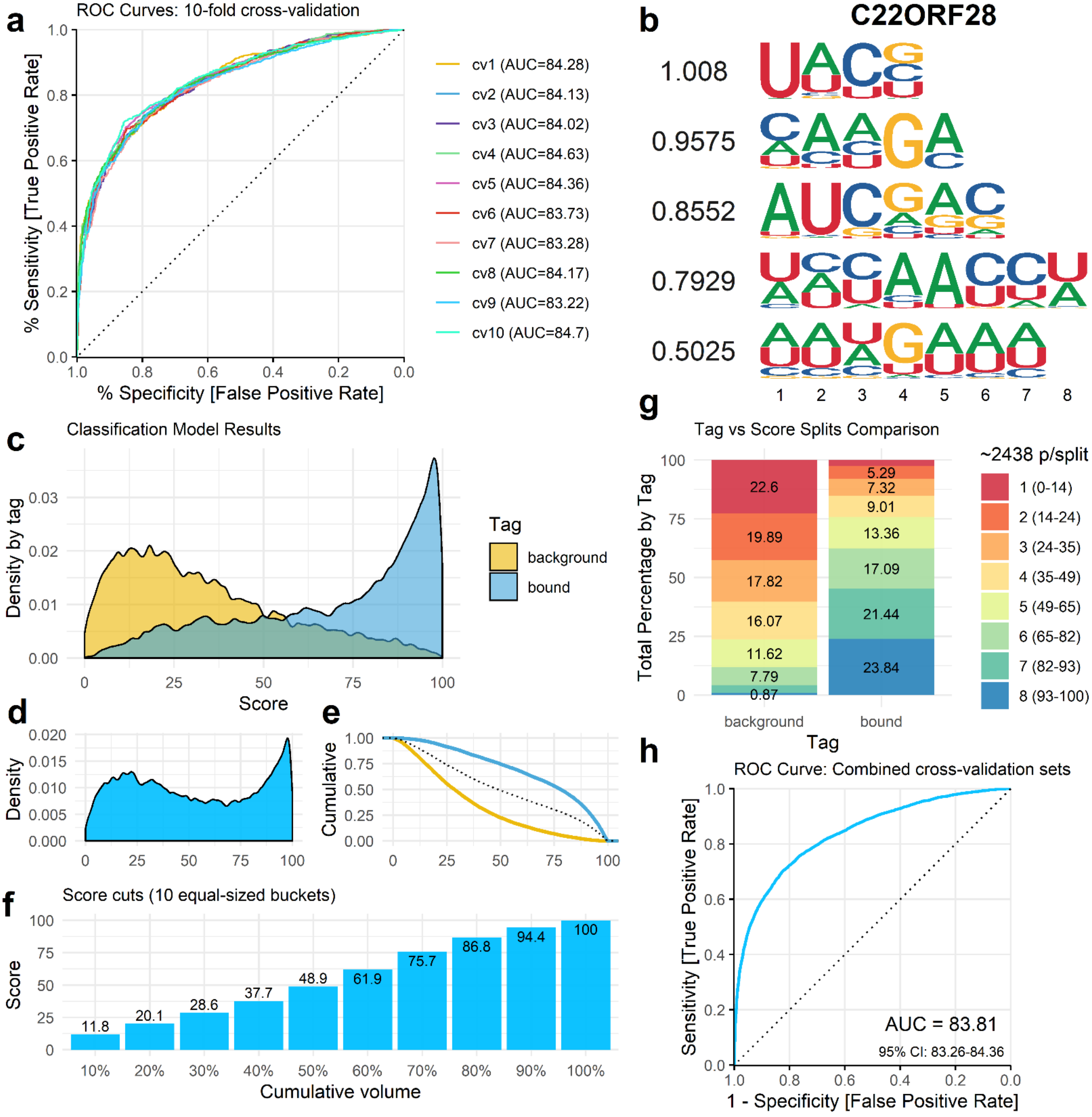

a
ROC Curves: 10-fold cross-validation
% Sensitivity [True Positive Rate]
% Specificity [False Positive Rate]
cv1 (AUC=84.28)
cv2 (AUC=84.13)
cv3 (AUC=84.02)
cv4 (AUC=84.63)
cv5 (AUC=84.36)
cv6 (AUC=83.73)
cv7 (AUC=83.28)
cv8 (AUC=84.17)
cv9 (AUC=83.22)
cv10 (AUC=84.7)
b
C22ORF28
1.008
0.9575
0.8552
0.7929
0.5025
c
Classification Model Results
Density by tag
Score
Tag
background
bound
d
Density
e
Cumulative
f
Score cuts (10 equal-sized buckets)
Score
Cumulative volume
11.8
20.1
28.6
37.7
48.9
61.9
75.7
86.8
94.4
100
g
Tag vs Score Splits Comparison
Total Percentage by Tag
Tag
~2438 p/split
1 (0-14)
2 (14-24)
3 (24-35)
4 (35-49)
5 (49-65)
6 (65-82)
7 (82-93)
8 (93-100)
22.6
19.89
17.82
16.07
11.62
7.79
0.87
5.29
7.32
9.01
13.36
17.09
21.44
23.84
h
ROC Curve: Combined cross-validation sets
Sensitivity [True Positive Rate]
1 - Specificity [False Positive Rate]
AUC = 83.81
95% CI: 83.26-84.36

Fig. S10

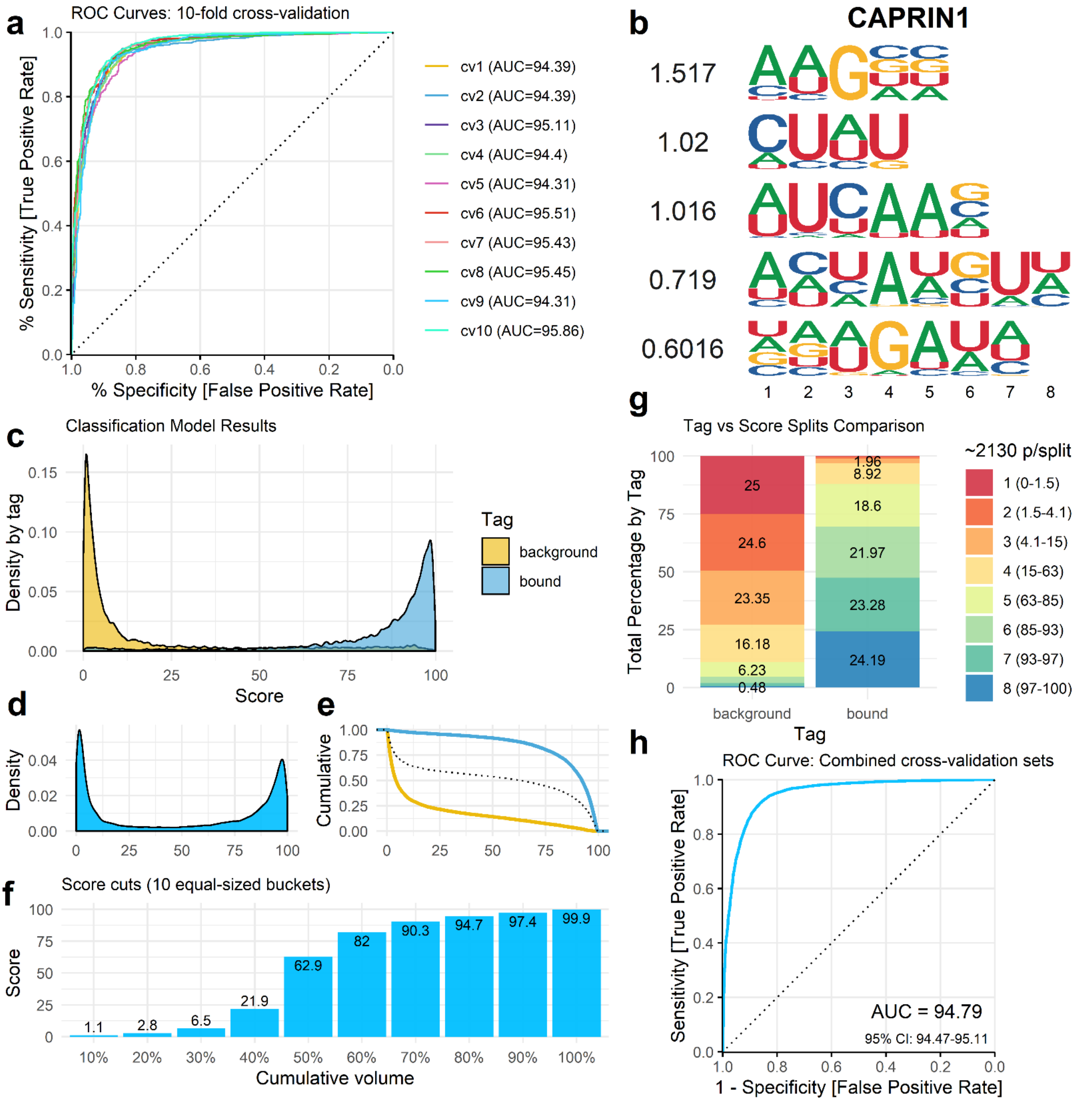

a
ROC Curves: 10-fold cross-validation
% Sensitivity [True Positive Rate]
% Specificity [False Positive Rate]
cv1 (AUC=94.39)
cv2 (AUC=94.39)
cv3 (AUC=95.11)
cv4 (AUC=94.4)
cv5 (AUC=94.31)
cv6 (AUC=95.51)
cv7 (AUC=95.43)
cv8 (AUC=95.45)
cv9 (AUC=94.31)
cv10 (AUC=95.86)
b
CAPRIN1
1.517
1.02
1.016
0.719
0.6016
c
Classification Model Results
Density by tag
Score
Tag
background
bound
d
Density
e
Cumulative
f
Score cuts (10 equal-sized buckets)
Score
Cumulative volume
1.1
2.8
6.5
21.9
62.9
82
90.3
94.7
97.4
99.9
g
Tag vs Score Splits Comparison
Total Percentage by Tag
~2130 p/split
1 (0-1.5)
2 (1.5-4.1)
3 (4.1-15)
4 (15-63)
5 (63-85)
6 (85-93)
7 (93-97)
8 (97-100)
25
24.6
23.35
16.18
6.23
0.48
1.96
8.92
18.6
21.97
23.28
24.19
background
bound
Tag
h
ROC Curve: Combined cross-validation sets
Sensitivity [True Positive Rate]
1 - Specificity [False Positive Rate]
AUC = 94.79
95% CI: 94.47-95.11

Fig. S11

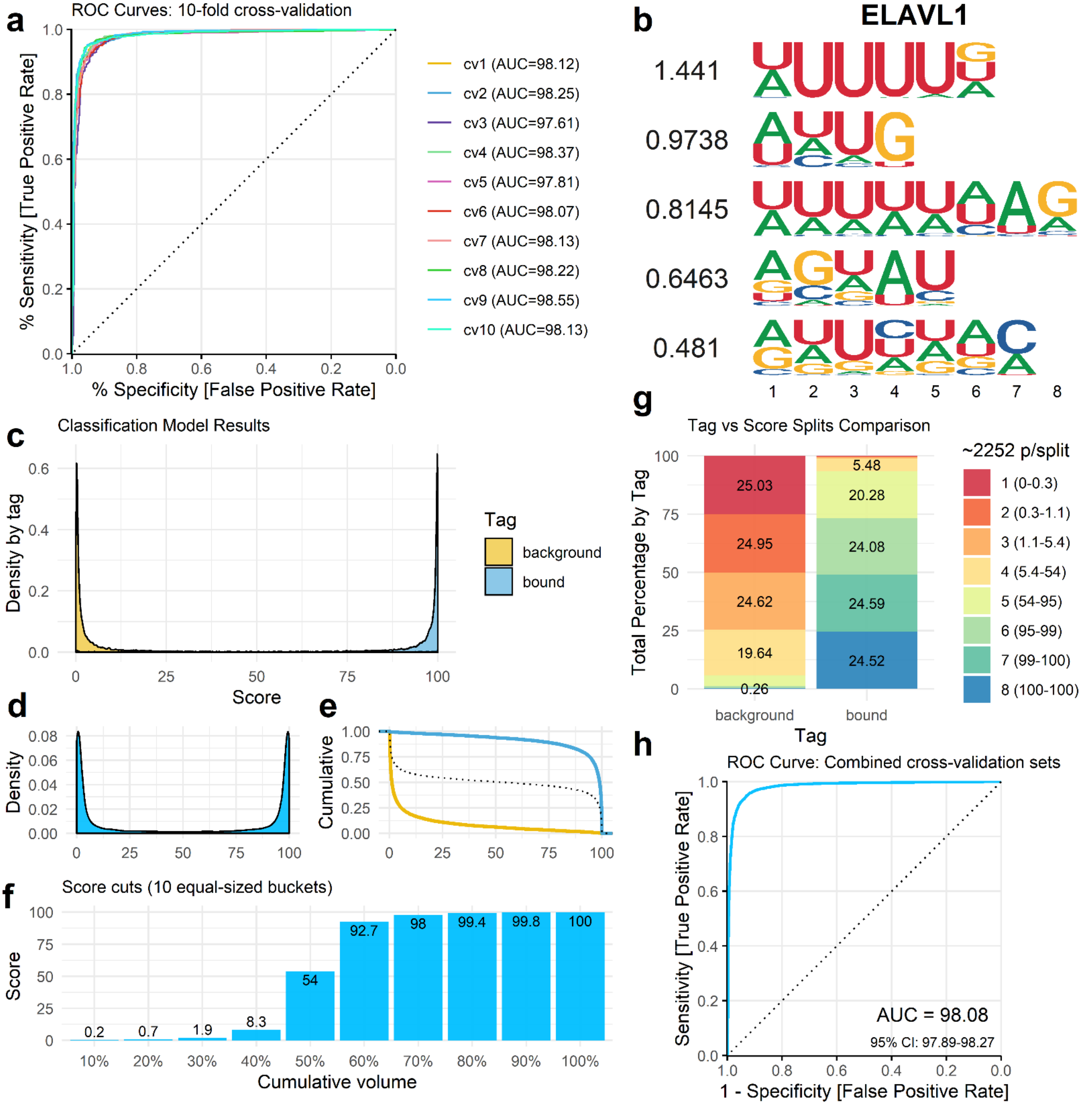

a
ROC Curves: 10-fold cross-validation
% Sensitivity [True Positive Rate]
% Specificity [False Positive Rate]
cv1 (AUC=98.12)
cv2 (AUC=98.25)
cv3 (AUC=97.61)
cv4 (AUC=98.37)
cv5 (AUC=97.81)
cv6 (AUC=98.07)
cv7 (AUC=98.13)
cv8 (AUC=98.22)
cv9 (AUC=98.55)
cv10 (AUC=98.13)
b
ELAVL1
1.441
0.9738
0.8145
0.6463
0.481
c
Classification Model Results
Density by tag
Score
Tag
background
bound
d
Density
e
Cumulative
f
Score cuts (10 equal-sized buckets)
Score
Cumulative volume
0.2
0.7
1.9
8.3
54
92.7
98
99.4
99.8
100
10%
20%
30%
40%
50%
60%
70%
80%
90%
100%
g
Tag vs Score Splits Comparison
Total Percentage by Tag
~2252 p/split
1 (0-0.3)
2 (0.3-1.1)
3 (1.1-5.4)
4 (5.4-54)
5 (54-95)
6 (95-99)
7 (99-100)
8 (100-100)
25.03
24.95
24.62
19.64
0.26
5.48
20.28
24.08
24.59
24.52
background
bound
Tag
h
ROC Curve: Combined cross-validation sets
Sensitivity [True Positive Rate]
1 - Specificity [False Positive Rate]
AUC = 98.08
95% CI: 97.89-98.27

Fig. S12

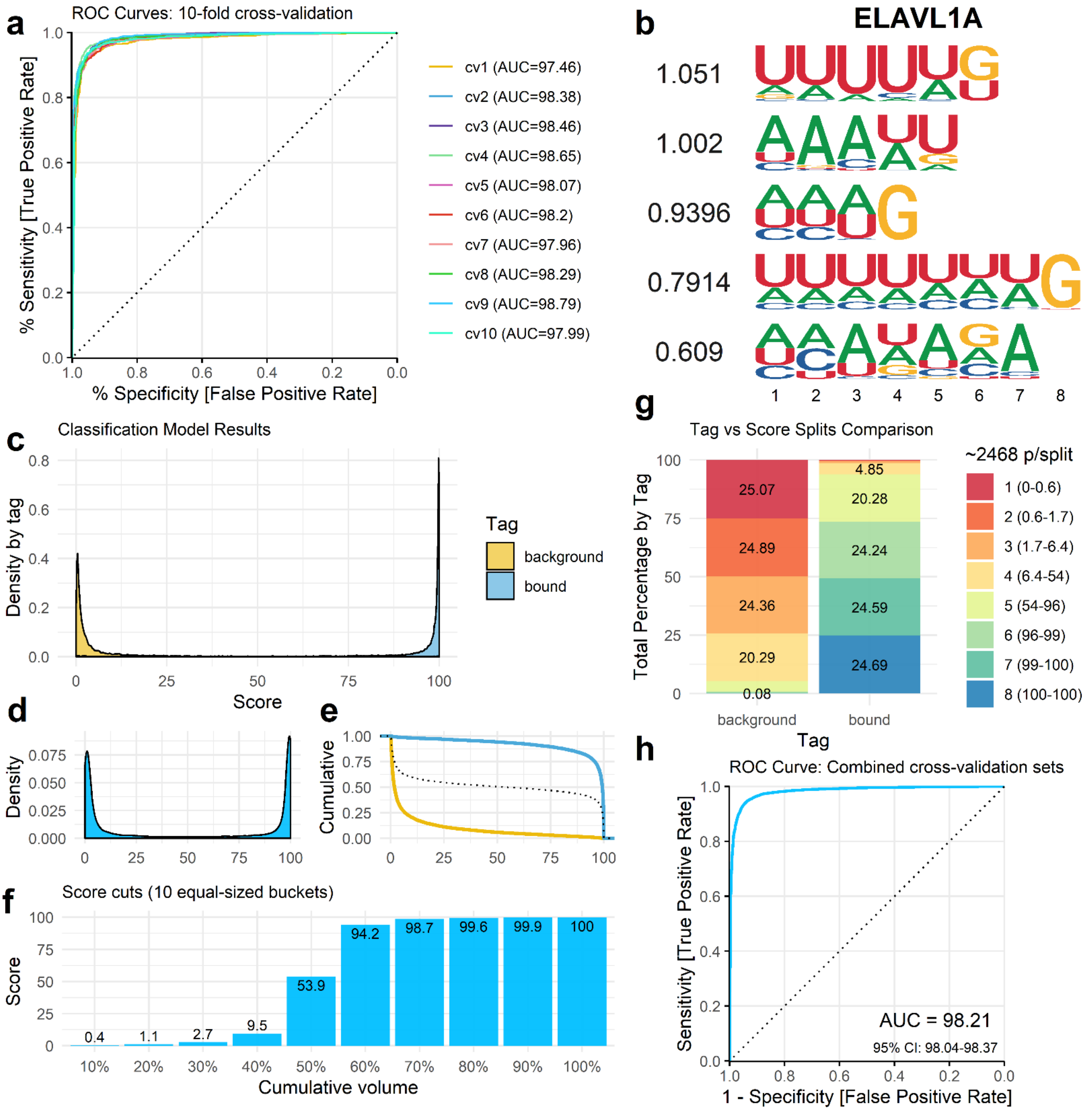
a
ROC Curves: 10-fold cross-validation
% Sensitivity [True Positive Rate]
% Specificity [False Positive Rate]
1.0
0.8
0.6
0.4
0.2
0.0
cv1 (AUC=97.46)
cv2 (AUC=98.38)
cv3 (AUC=98.46)
cv4 (AUC=98.65)
cv5 (AUC=98.07)
cv6 (AUC=98.2)
cv7 (AUC=97.96)
cv8 (AUC=98.29)
cv9 (AUC=98.79)
cv10 (AUC=97.99)
b
ELAVL1A
1.051
1.002
0.9396
0.7914
0.609
1 2 3 4 5 6 7 8
c
Classification Model Results
Density by tag
Score
Tag
background
bound
0 25 50 75 100
d
Density
0.075
0.050
0.025
0.000
e
Cumulative
1.00
0.75
0.50
0.25
0.00
f
Score cuts (10 equal-sized buckets)
Score
Cumulative volume
0.4
1.1
2.7
9.5
53.9
94.2
98.7
99.6
99.9
100
10% 20% 30% 40% 50% 60% 70% 80% 90% 100%
g
Tag vs Score Splits Comparison
Total Percentage by Tag
Tag
background
bound
25.07
24.89
24.36
20.29
0.08
4.85
20.28
24.24
24.59
24.69
~2468 p/split
1 (0-0.6)
2 (0.6-1.7)
3 (1.7-6.4)
4 (6.4-54)
5 (54-96)
6 (96-99)
7 (99-100)
8 (100-100)
h
ROC Curve: Combined cross-validation sets
Sensitivity [True Positive Rate]
1 - Specificity [False Positive Rate]
AUC = 98.21
95% CI: 98.04-98.37

Fig. S13

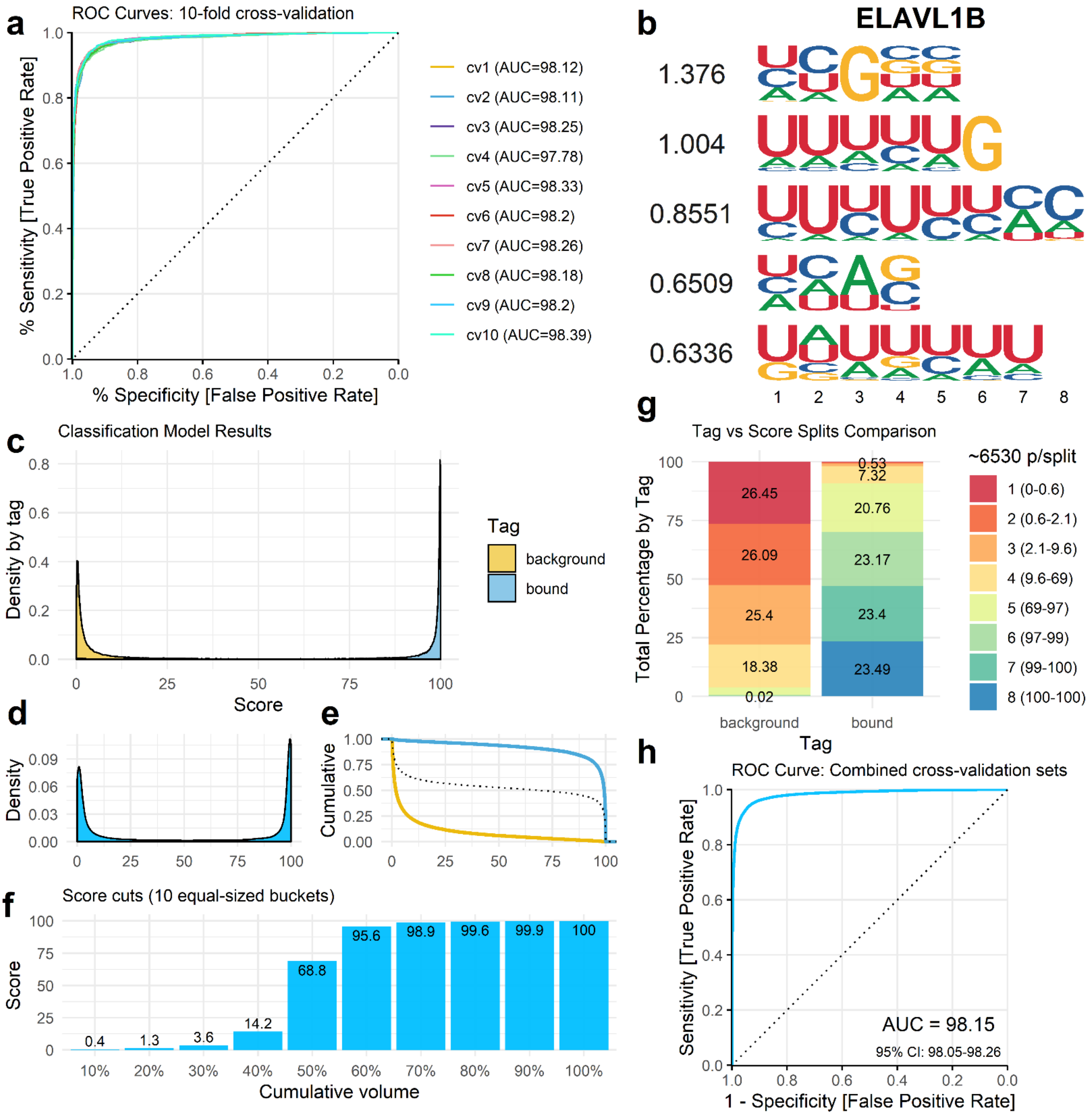

a
ROC Curves: 10-fold cross-validation
% Sensitivity [True Positive Rate]
% Specificity [False Positive Rate]
cv1 (AUC=98.12)
cv2 (AUC=98.11)
cv3 (AUC=98.25)
cv4 (AUC=97.78)
cv5 (AUC=98.33)
cv6 (AUC=98.2)
cv7 (AUC=98.26)
cv8 (AUC=98.18)
cv9 (AUC=98.2)
cv10 (AUC=98.39)
b
ELAVL1B
1.376
1.004
0.8551
0.6509
0.6336
c
Classification Model Results
Density by tag
Score
Tag
background
bound
d
Density
e
Cumulative
f
Score cuts (10 equal-sized buckets)
Score
Cumulative volume
0.4
1.3
3.6
14.2
68.8
95.6
98.9
99.6
99.9
100
g
Tag vs Score Splits Comparison
Total Percentage by Tag
~6530 p/split
1 (0-0.6)
2 (0.6-2.1)
3 (2.1-9.6)
4 (9.6-69)
5 (69-97)
6 (97-99)
7 (99-100)
8 (100-100)
26.45
26.09
25.4
18.38
0.02
0.53
7.32
20.76
23.17
23.4
23.49
background
bound
Tag
h
ROC Curve: Combined cross-validation sets
Sensitivity [True Positive Rate]
1 - Specificity [False Positive Rate]
AUC = 98.15
95% CI: 98.05-98.26

Fig. S14

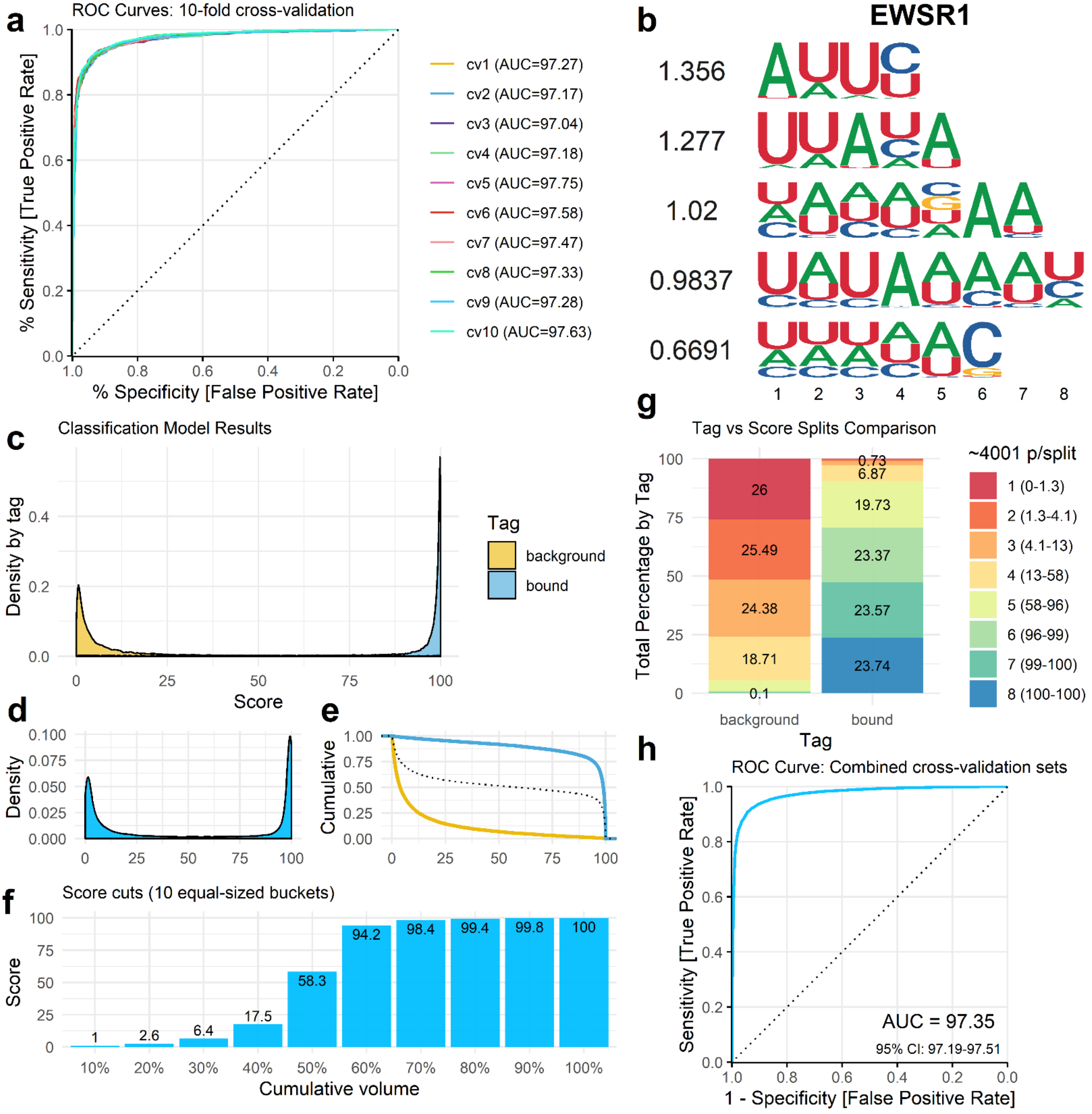

a
ROC Curves: 10-fold cross-validation
% Sensitivity [True Positive Rate]
% Specificity [False Positive Rate]
cv1 (AUC=97.27)
cv2 (AUC=97.17)
cv3 (AUC=97.04)
cv4 (AUC=97.18)
cv5 (AUC=97.75)
cv6 (AUC=97.58)
cv7 (AUC=97.47)
cv8 (AUC=97.33)
cv9 (AUC=97.28)
cv10 (AUC=97.63)
b
EWSR1
1.356
1.277
1.02
0.9837
0.6691
c
Classification Model Results
Density by tag
Score
Tag
background
bound
d
Density
e
Cumulative
f
Score cuts (10 equal-sized buckets)
Score
Cumulative volume
1
2.6
6.4
17.5
58.3
94.2
98.4
99.4
99.8
100
g
Tag vs Score Splits Comparison
Total Percentage by Tag
Tag
~4001 p/split
1 (0-1.3)
2 (1.3-4.1)
3 (4.1-13)
4 (13-58)
5 (58-96)
6 (96-99)
7 (99-100)
8 (100-100)
26
25.49
24.38
18.71
0.1
0.73
6.87
19.73
23.37
23.57
23.74
h
ROC Curve: Combined cross-validation sets
Sensitivity [True Positive Rate]
1 - Specificity [False Positive Rate]
AUC = 97.35
95% CI: 97.19-97.51

### Fig. S15

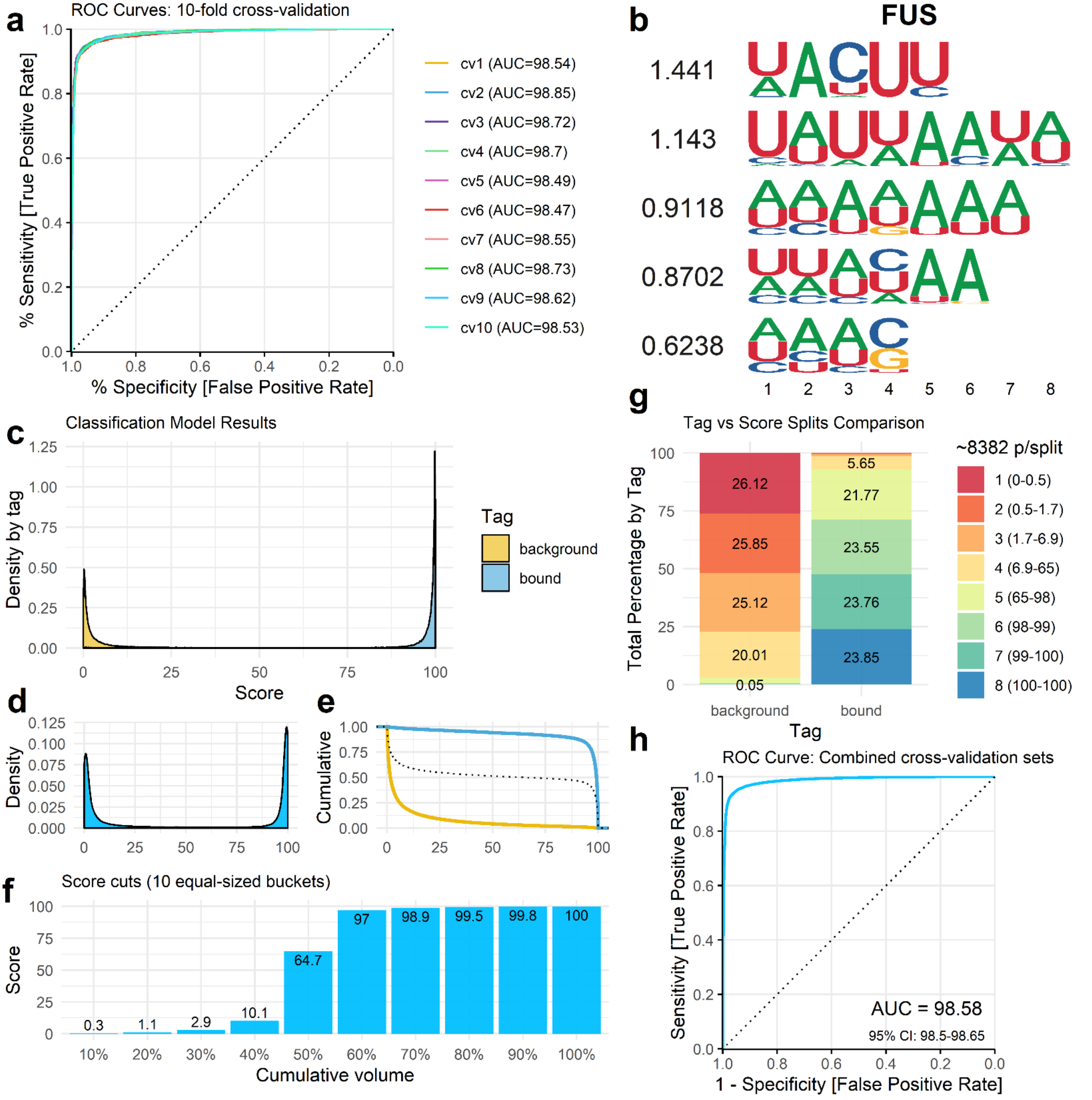
a
ROC Curves: 10-fold cross-validation
% Sensitivity [True Positive Rate]
% Specificity [False Positive Rate]
cv1 (AUC=98.54)
cv2 (AUC=98.85)
cv3 (AUC=98.72)
cv4 (AUC=98.7)
cv5 (AUC=98.49)
cv6 (AUC=98.47)
cv7 (AUC=98.55)
cv8 (AUC=98.73)
cv9 (AUC=98.62)
cv10 (AUC=98.53)
b
FUS
1.441
1.143
0.9118
0.8702
0.6238
c
Classification Model Results
Density by tag
Score
Tag
background
bound
d
Density
e
Cumulative
f
Score cuts (10 equal-sized buckets)
Score
Cumulative volume
0.3
1.1
2.9
10.1
64.7
97
98.9
99.5
99.8
100
g
Tag vs Score Splits Comparison
Total Percentage by Tag
~8382 p/split
1 (0-0.5)
2 (0.5-1.7)
3 (1.7-6.9)
4 (6.9-65)
5 (65-98)
6 (98-99)
7 (99-100)
8 (100-100)
26.12
25.85
25.12
20.01
0.05
5.65
21.77
23.55
23.76
23.85
background
bound
Tag
h
ROC Curve: Combined cross-validation sets
Sensitivity [True Positive Rate]
1 - Specificity [False Positive Rate]
AUC = 98.58
95% CI: 98.5-98.65

Fig. S16

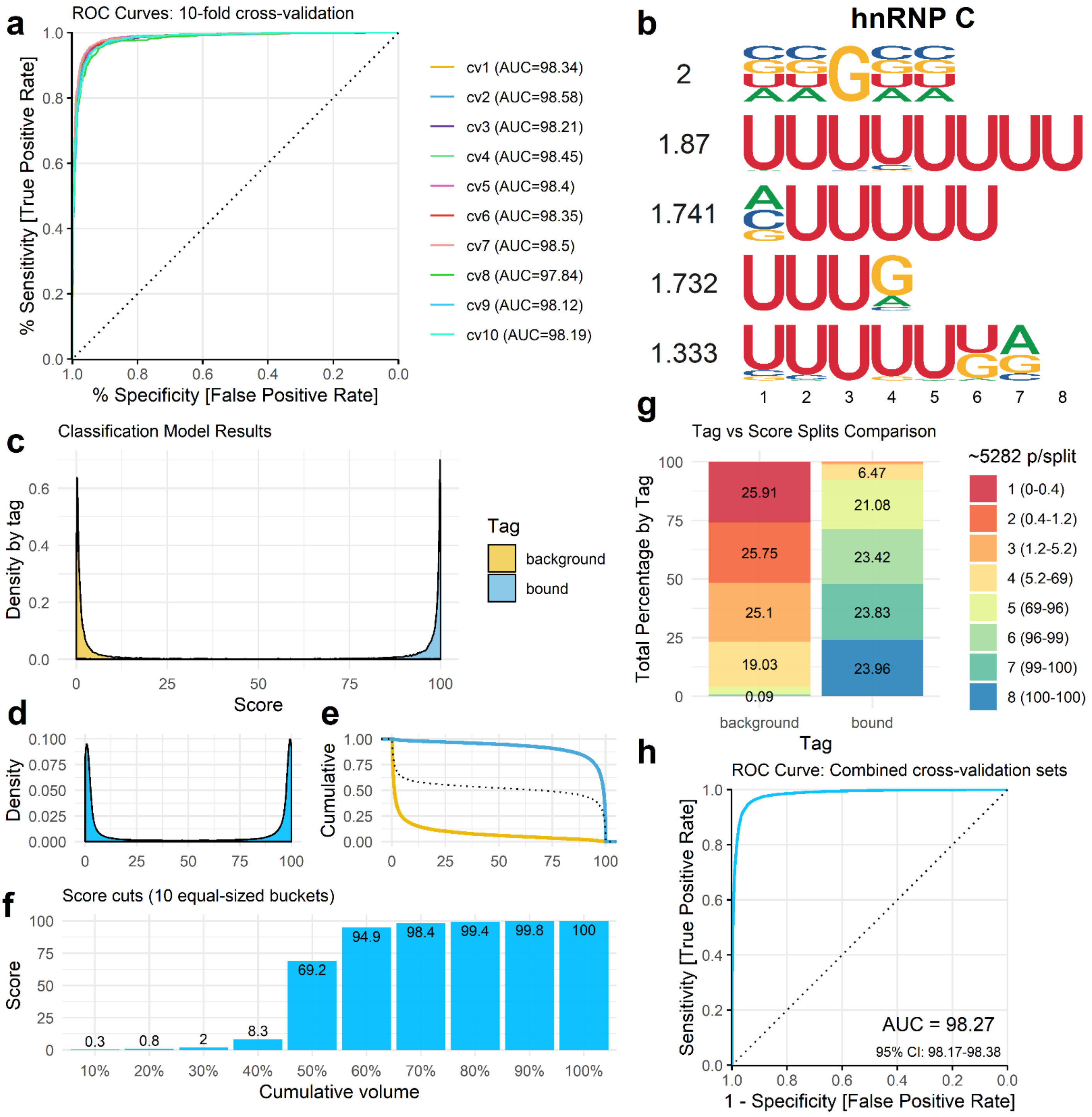
a
ROC Curves: 10-fold cross-validation
% Sensitivity [True Positive Rate]
% Specificity [False Positive Rate]
cv1 (AUC=98.34)
cv2 (AUC=98.58)
cv3 (AUC=98.21)
cv4 (AUC=98.45)
cv5 (AUC=98.4)
cv6 (AUC=98.35)
cv7 (AUC=98.5)
cv8 (AUC=97.84)
cv9 (AUC=98.12)
cv10 (AUC=98.19)
b
hnRNP C
2
1.87
1.741
1.732
1.333
c
Classification Model Results
Density by tag
Score
Tag
background
bound
d
Density
e
Cumulative
f
Score cuts (10 equal-sized buckets)
Score
Cumulative volume
0.3
0.8
2
8.3
69.2
94.9
98.4
99.4
99.8
100
10%
20%
30%
40%
50%
60%
70%
80%
90%
100%
g
Tag vs Score Splits Comparison
Total Percentage by Tag
25.91
25.75
25.1
19.03
0.09
6.47
21.08
23.42
23.83
23.96
background
bound
Tag
~5282 p/split
1 (0-0.4)
2 (0.4-1.2)
3 (1.2-5.2)
4 (5.2-69)
5 (69-96)
6 (96-99)
7 (99-100)
8 (100-100)
h
ROC Curve: Combined cross-validation sets
Sensitivity [True Positive Rate]
1 - Specificity [False Positive Rate]
AUC = 98.27
95% CI: 98.17-98.38

Fig. S17

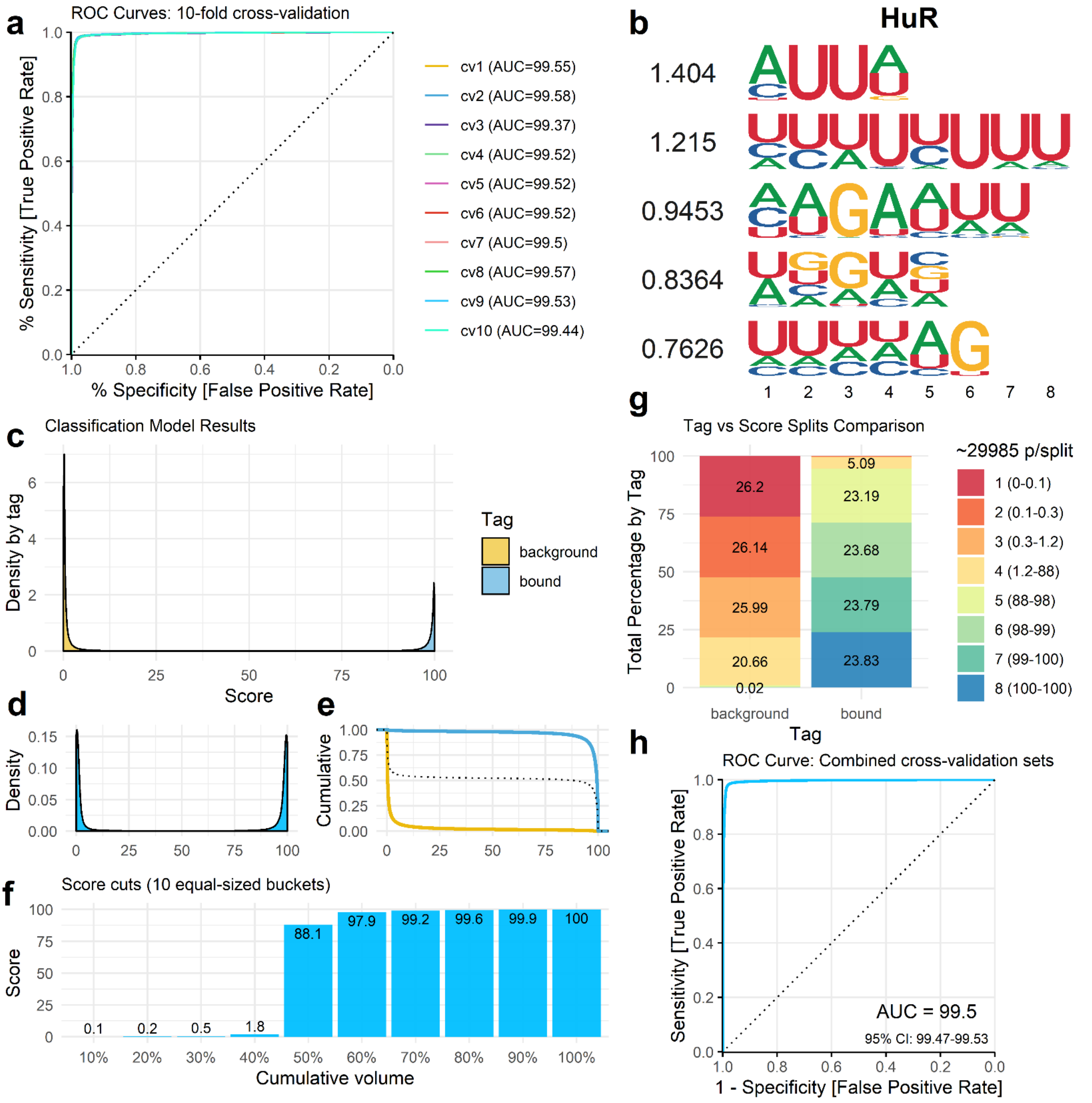

a
ROC Curves: 10-fold cross-validation
% Sensitivity [True Positive Rate]
% Specificity [False Positive Rate]
cv1 (AUC=99.55)
cv2 (AUC=99.58)
cv3 (AUC=99.37)
cv4 (AUC=99.52)
cv5 (AUC=99.52)
cv6 (AUC=99.52)
cv7 (AUC=99.5)
cv8 (AUC=99.57)
cv9 (AUC=99.53)
cv10 (AUC=99.44)
b
HuR
1.404
1.215
0.9453
0.8364
0.7626
c
Classification Model Results
Density by tag
Score
Tag
background
bound
d
Density
e
Cumulative
f
Score cuts (10 equal-sized buckets)
Score
Cumulative volume
0.1
0.2
0.5
1.8
88.1
97.9
99.2
99.6
99.9
100
g
Tag vs Score Splits Comparison
Total Percentage by Tag
Tag
~29985 p/split
1 (0-0.1)
2 (0.1-0.3)
3 (0.3-1.2)
4 (1.2-88)
5 (88-98)
6 (98-99)
7 (99-100)
8 (100-100)
26.2
26.14
25.99
20.66
0.02
5.09
23.19
23.68
23.79
23.83
h
ROC Curve: Combined cross-validation sets
Sensitivity [True Positive Rate]
1 - Specificity [False Positive Rate]
AUC = 99.5
95% CI: 99.47-99.53

Fig. S18

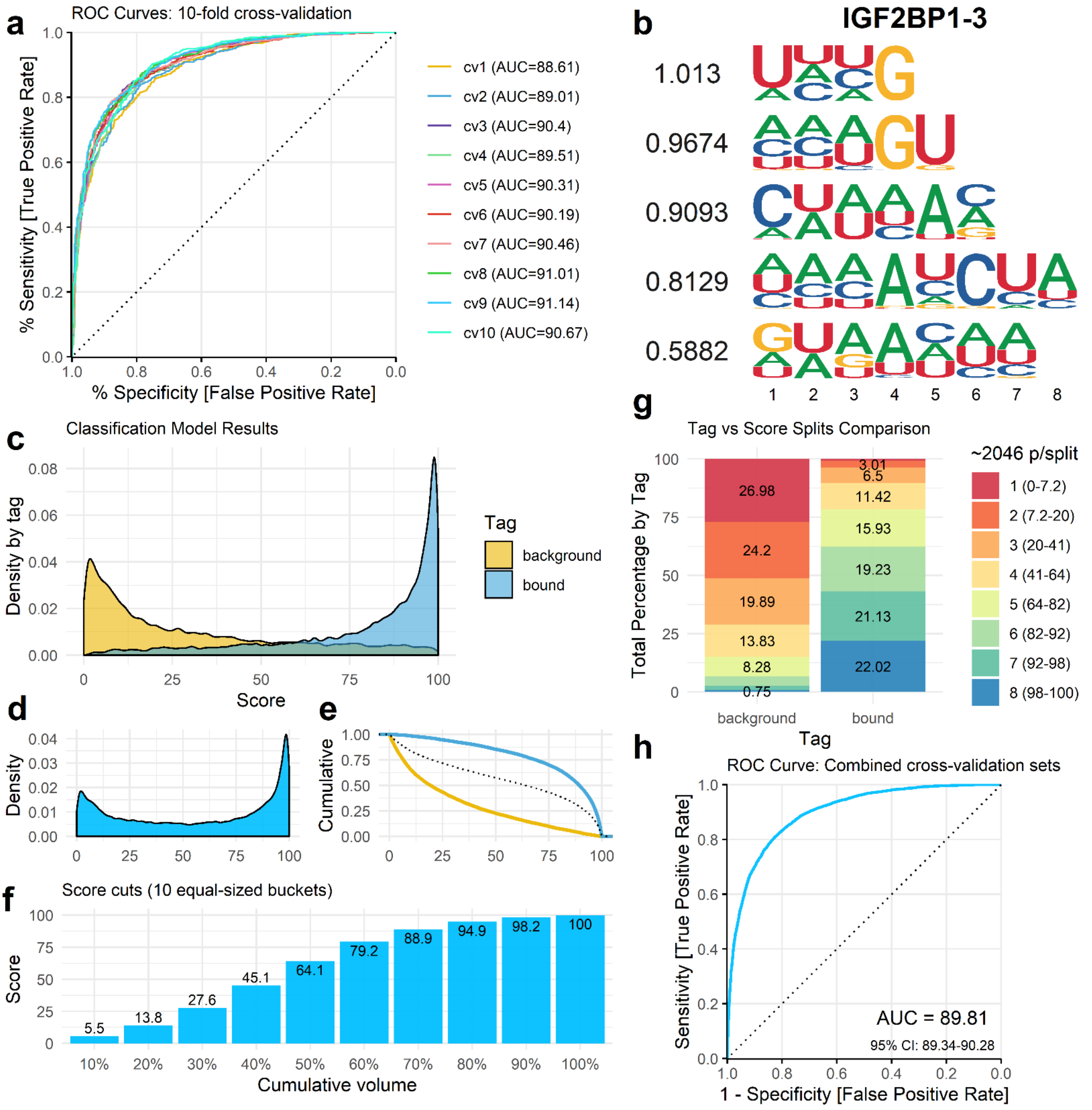

a
ROC Curves: 10-fold cross-validation
% Sensitivity [True Positive Rate]
% Specificity [False Positive Rate]
cv1 (AUC=88.61)
cv2 (AUC=89.01)
cv3 (AUC=90.4)
cv4 (AUC=89.51)
cv5 (AUC=90.31)
cv6 (AUC=90.19)
cv7 (AUC=90.46)
cv8 (AUC=91.01)
cv9 (AUC=91.14)
cv10 (AUC=90.67)
b
IGF2BP1-3
1.013
0.9674
0.9093
0.8129
0.5882
c
Classification Model Results
Density by tag
Score
Tag
background
bound
d
Density
e
Cumulative
f
Score cuts (10 equal-sized buckets)
Score
Cumulative volume
5.5
13.8
27.6
45.1
64.1
79.2
88.9
94.9
98.2
100
g
Tag vs Score Splits Comparison
Total Percentage by Tag
Tag
~2046 p/split
1 (0-7.2)
2 (7.2-20)
3 (20-41)
4 (41-64)
5 (64-82)
6 (82-92)
7 (92-98)
8 (98-100)
background: 26.98, 24.2, 19.89, 13.83, 8.28, 0.75
bound: 3.01, 6.5, 11.42, 15.93, 19.23, 21.13, 22.02
h
ROC Curve: Combined cross-validation sets
Sensitivity [True Positive Rate]
1 - Specificity [False Positive Rate]
AUC = 89.81
95% CI: 89.34-90.28

Fig. S19

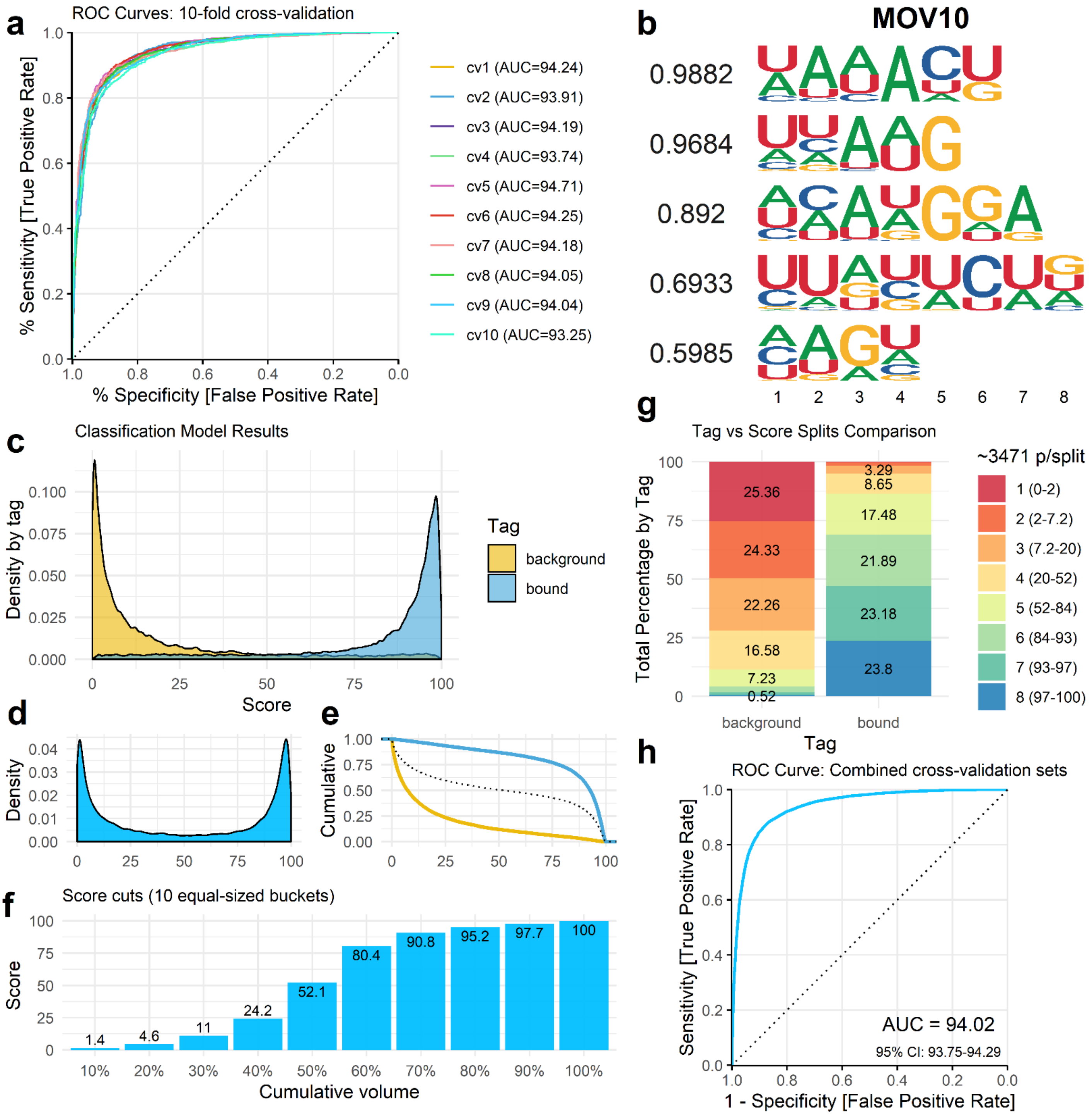
a
ROC Curves: 10-fold cross-validation
% Sensitivity [True Positive Rate]
% Specificity [False Positive Rate]
cv1 (AUC=94.24)
cv2 (AUC=93.91)
cv3 (AUC=94.19)
cv4 (AUC=93.74)
cv5 (AUC=94.71)
cv6 (AUC=94.25)
cv7 (AUC=94.18)
cv8 (AUC=94.05)
cv9 (AUC=94.04)
cv10 (AUC=93.25)
b
MOV10
0.9882
0.9684
0.892
0.6933
0.5985
1 2 3 4 5 6 7 8
c
Classification Model Results
Density by tag
Score
Tag
background
bound
d
Density
e
Cumulative
f
Score cuts (10 equal-sized buckets)
Score
Cumulative volume
1.4
4.6
11
24.2
52.1
80.4
90.8
95.2
97.7
100
10% 20% 30% 40% 50% 60% 70% 80% 90% 100%
g
Tag vs Score Splits Comparison
Total Percentage by Tag
~3471 p/split
25.36
24.33
22.26
16.58
7.23
0.52
3.29
8.65
17.48
21.89
23.18
23.8
background
bound
Tag
1 (0-2)
2 (2-7.2)
3 (7.2-20)
4 (20-52)
5 (52-84)
6 (84-93)
7 (93-97)
8 (97-100)
h
ROC Curve: Combined cross-validation sets
Sensitivity [True Positive Rate]
1 - Specificity [False Positive Rate]
AUC = 94.02
95% CI: 93.75-94.29

Fig. S20

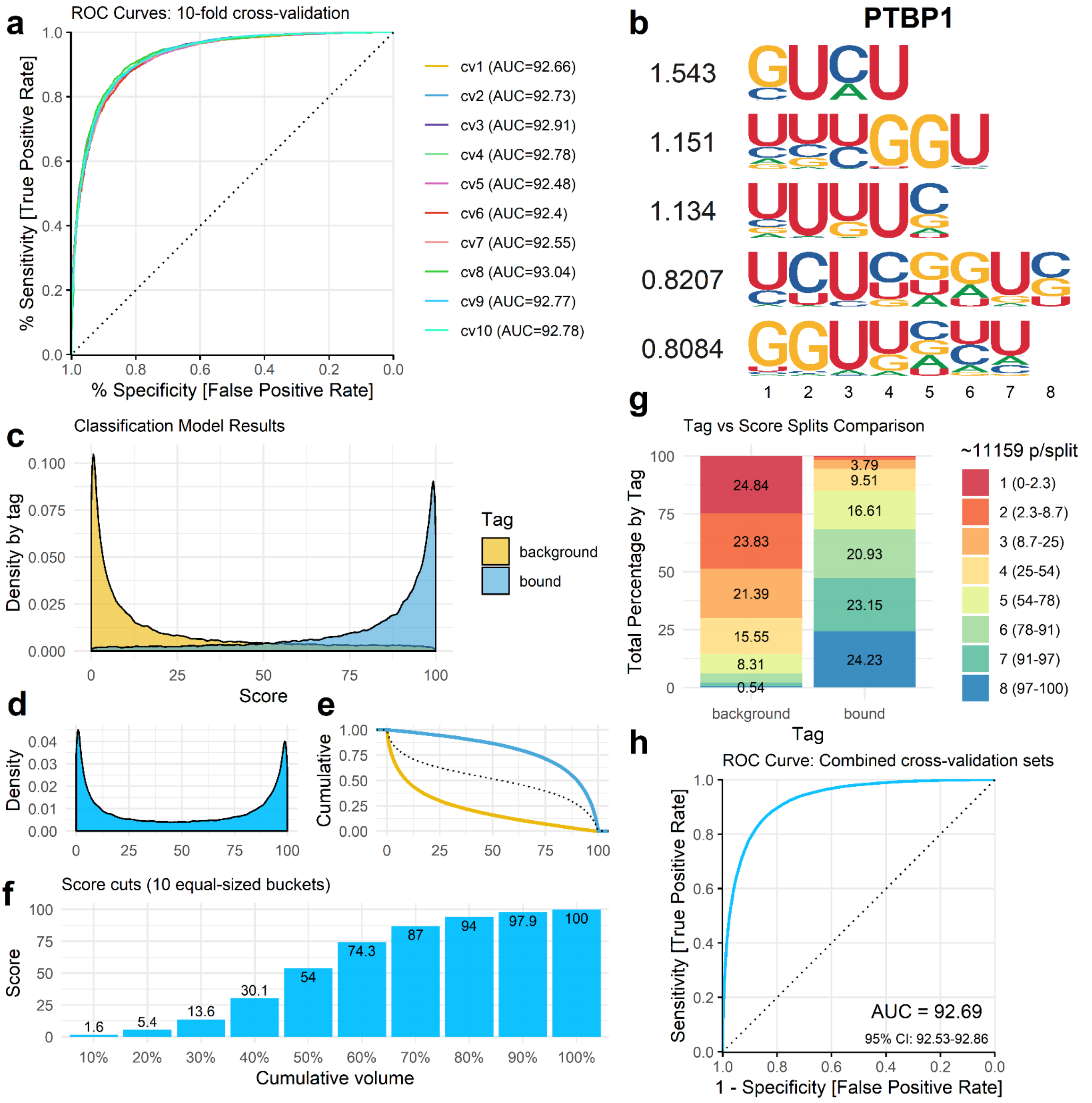

a
ROC Curves: 10-fold cross-validation
% Sensitivity [True Positive Rate]
% Specificity [False Positive Rate]
cv1 (AUC=92.66)
cv2 (AUC=92.73)
cv3 (AUC=92.91)
cv4 (AUC=92.78)
cv5 (AUC=92.48)
cv6 (AUC=92.4)
cv7 (AUC=92.55)
cv8 (AUC=93.04)
cv9 (AUC=92.77)
cv10 (AUC=92.78)
b
PTBP1
1.543
1.151
1.134
0.8207
0.8084
c
Classification Model Results
Density by tag
Score
Tag
background
bound
d
Density
e
Cumulative
f
Score cuts (10 equal-sized buckets)
Score
Cumulative volume
1.6
5.4
13.6
30.1
54
74.3
87
94
97.9
100
g
Tag vs Score Splits Comparison
Total Percentage by Tag
~11159 p/split
1 (0-2.3)
2 (2.3-8.7)
3 (8.7-25)
4 (25-54)
5 (54-78)
6 (78-91)
7 (91-97)
8 (97-100)
24.84
23.83
21.39
15.55
8.31
0.54
3.79
9.51
16.61
20.93
23.15
24.23
background
bound
Tag
h
ROC Curve: Combined cross-validation sets
Sensitivity [True Positive Rate]
1 - Specificity [False Positive Rate]
AUC = 92.69
95% CI: 92.53-92.86

Fig. S21

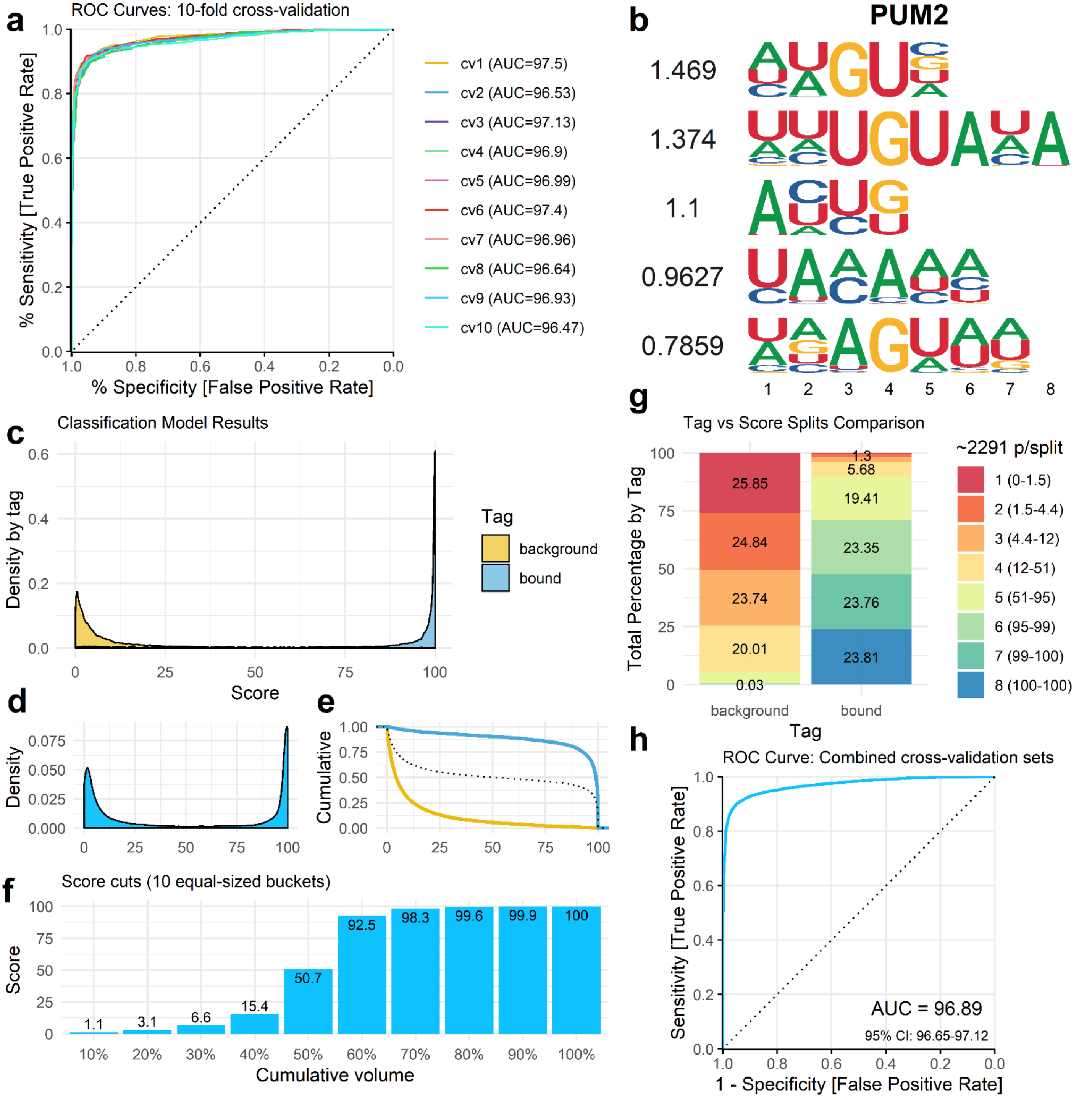
a
ROC Curves: 10-fold cross-validation
% Sensitivity [True Positive Rate]
% Specificity [False Positive Rate]
cv1 (AUC=97.5)
cv2 (AUC=96.53)
cv3 (AUC=97.13)
cv4 (AUC=96.9)
cv5 (AUC=96.99)
cv6 (AUC=97.4)
cv7 (AUC=96.96)
cv8 (AUC=96.64)
cv9 (AUC=96.93)
cv10 (AUC=96.47)
b
PUM2
1.469
1.374
1.1
0.9627
0.7859
c
Classification Model Results
Density by tag
Score
Tag
background
bound
d
Density
e
Cumulative
f
Score cuts (10 equal-sized buckets)
Score
Cumulative volume
1.1
3.1
6.6
15.4
50.7
92.5
98.3
99.6
99.9
100
g
Tag vs Score Splits Comparison
Total Percentage by Tag
~2291 p/split
1 (0-1.5)
2 (1.5-4.4)
3 (4.4-12)
4 (12-51)
5 (51-95)
6 (95-99)
7 (99-100)
8 (100-100)
25.85
24.84
23.74
20.01
0.03
1.3
5.68
19.41
23.35
23.76
23.81
background
bound
Tag
h
ROC Curve: Combined cross-validation sets
Sensitivity [True Positive Rate]
1 - Specificity [False Positive Rate]
AUC = 96.89
95% CI: 96.65-97.12

### Fig. S22

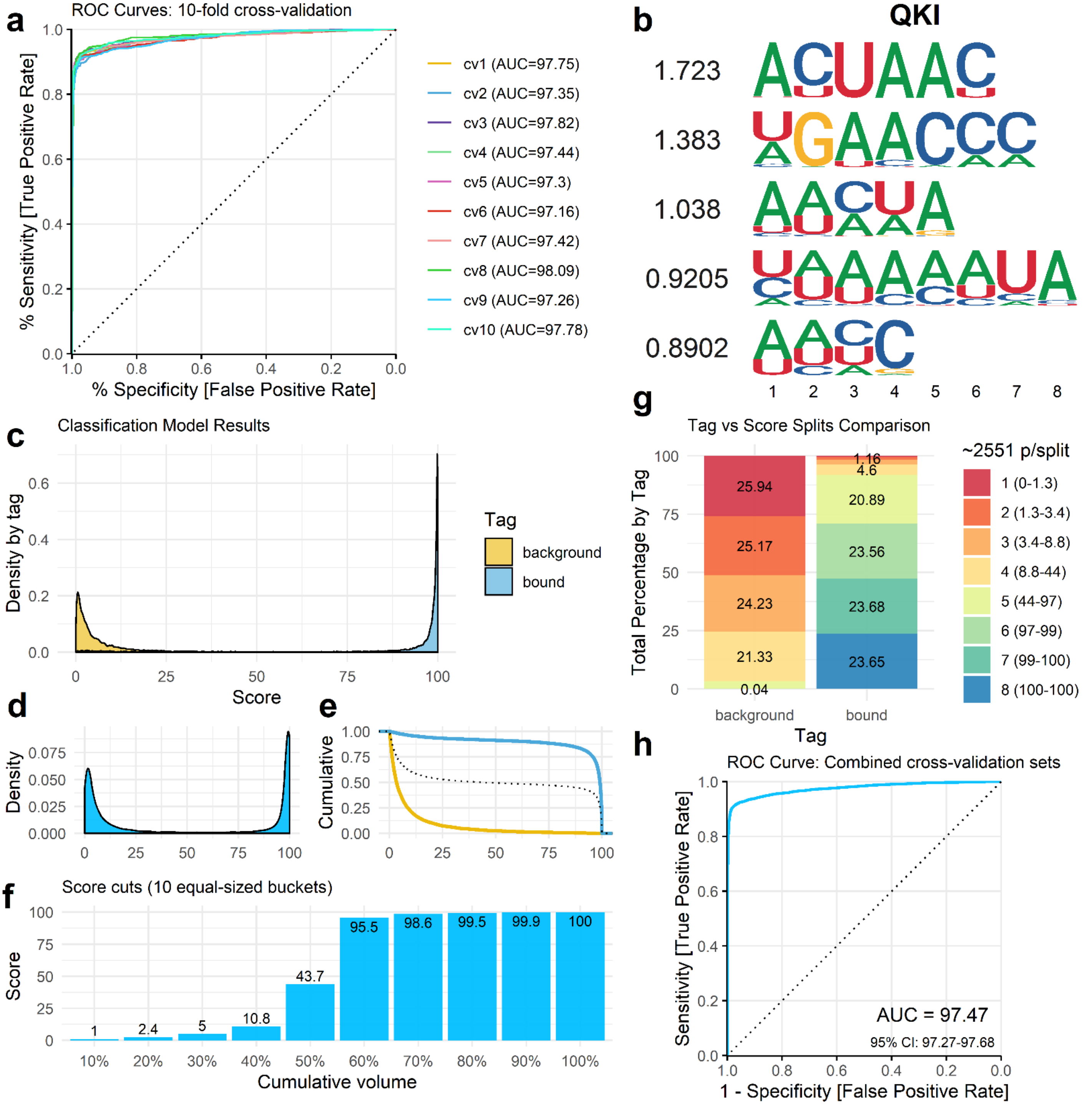

a
ROC Curves: 10-fold cross-validation
% Sensitivity [True Positive Rate]
% Specificity [False Positive Rate]
cv1 (AUC=97.75)
cv2 (AUC=97.35)
cv3 (AUC=97.82)
cv4 (AUC=97.44)
cv5 (AUC=97.3)
cv6 (AUC=97.16)
cv7 (AUC=97.42)
cv8 (AUC=98.09)
cv9 (AUC=97.26)
cv10 (AUC=97.78)
b
QKI
1.723
1.383
1.038
0.9205
0.8902
c
Classification Model Results
Density by tag
Score
Tag
background
bound
d
Density
e
Cumulative
f
Score cuts (10 equal-sized buckets)
Score
Cumulative volume
1
2.4
5
10.8
43.7
95.5
98.6
99.5
99.9
100
g
Tag vs Score Splits Comparison
Total Percentage by Tag
~2551 p/split
1 (0-1.3)
2 (1.3-3.4)
3 (3.4-8.8)
4 (8.8-44)
5 (44-97)
6 (97-99)
7 (99-100)
8 (100-100)
25.94
25.17
24.23
21.33
0.04
1.16
4.6
20.89
23.56
23.68
23.65
background
bound
Tag
h
ROC Curve: Combined cross-validation sets
Sensitivity [True Positive Rate]
1 - Specificity [False Positive Rate]
AUC = 97.47
95% CI: 97.27-97.68

Fig. S23

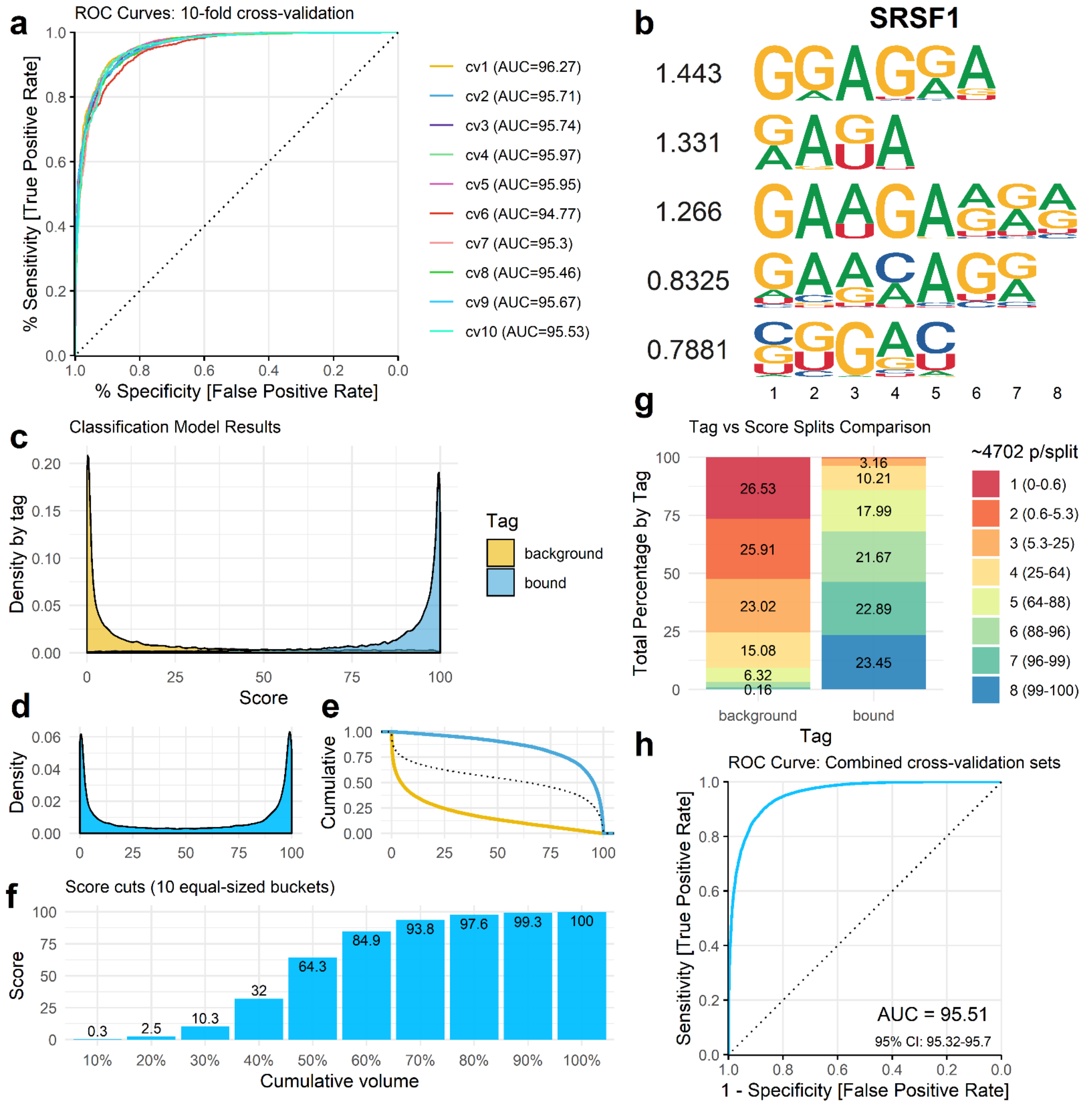

a
ROC Curves: 10-fold cross-validation
% Sensitivity [True Positive Rate]
% Specificity [False Positive Rate]
cv1 (AUC=96.27)
cv2 (AUC=95.71)
cv3 (AUC=95.74)
cv4 (AUC=95.97)
cv5 (AUC=95.95)
cv6 (AUC=94.77)
cv7 (AUC=95.3)
cv8 (AUC=95.46)
cv9 (AUC=95.67)
cv10 (AUC=95.53)
b
SRSF1
1.443
1.331
1.266
0.8325
0.7881
1 2 3 4 5 6 7 8
c
Classification Model Results
Density by tag
Score
Tag
background
bound
d
Density
e
Cumulative
f
Score cuts (10 equal-sized buckets)
Score
Cumulative volume
0.3
2.5
10.3
32
64.3
84.9
93.8
97.6
99.3
100
10% 20% 30% 40% 50% 60% 70% 80% 90% 100%
g
Tag vs Score Splits Comparison
Total Percentage by Tag
~4702 p/split
1 (0-0.6)
2 (0.6-5.3)
3 (5.3-25)
4 (25-64)
5 (64-88)
6 (88-96)
7 (96-99)
8 (99-100)
26.53
25.91
23.02
15.08
6.32
0.16
3.16
10.21
17.99
21.67
22.89
23.45
background
bound
Tag
h
ROC Curve: Combined cross-validation sets
Sensitivity [True Positive Rate]
1 - Specificity [False Positive Rate]
AUC = 95.51
95% CI: 95.32-95.7

Fig. S24

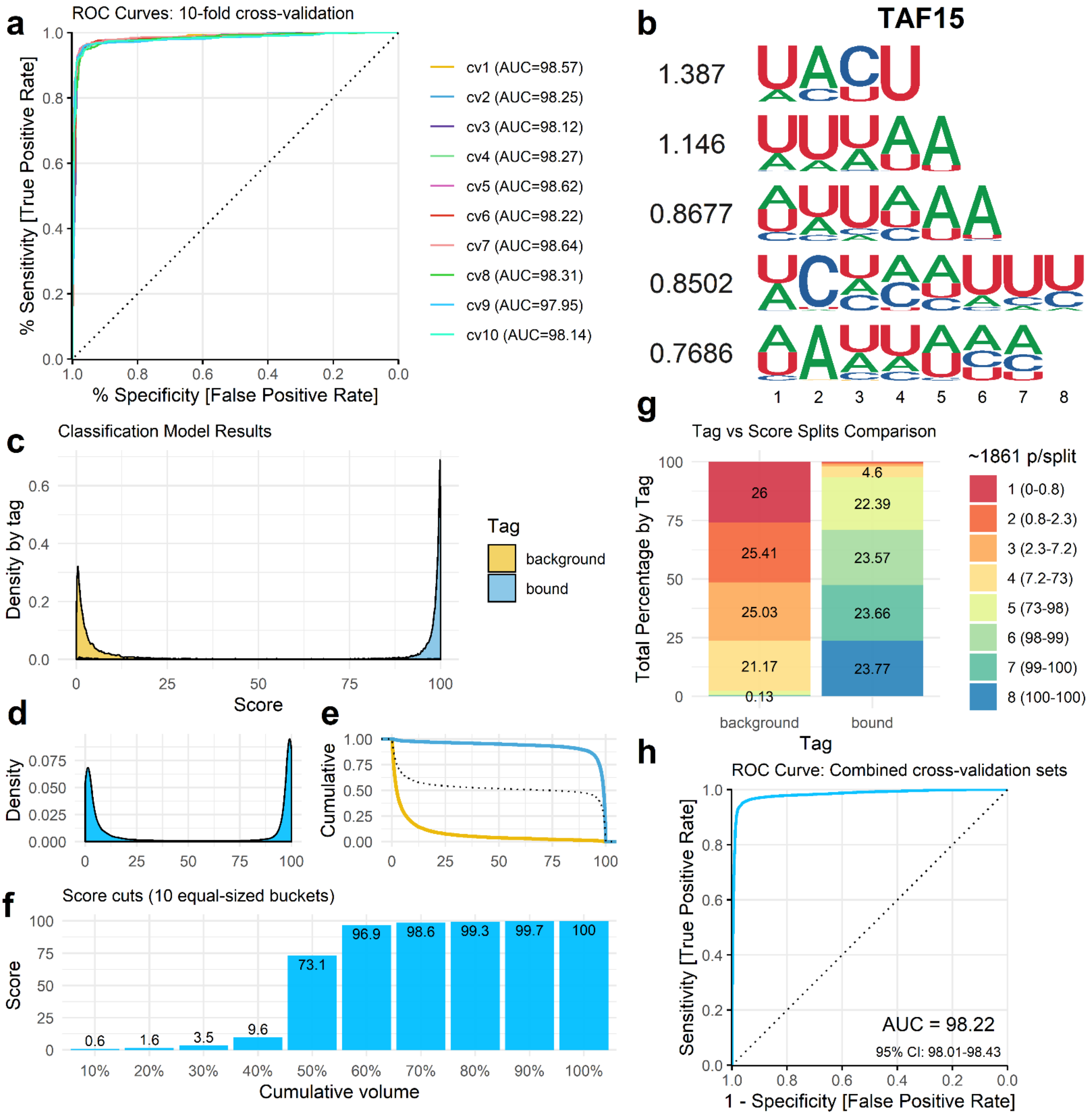

a
ROC Curves: 10-fold cross-validation
% Sensitivity [True Positive Rate]
% Specificity [False Positive Rate]
cv1 (AUC=98.57)
cv2 (AUC=98.25)
cv3 (AUC=98.12)
cv4 (AUC=98.27)
cv5 (AUC=98.62)
cv6 (AUC=98.22)
cv7 (AUC=98.64)
cv8 (AUC=98.31)
cv9 (AUC=97.95)
cv10 (AUC=98.14)
b
TAF15
1.387
1.146
0.8677
0.8502
0.7686
c
Classification Model Results
Density by tag
Score
Tag
background
bound
d
Density
e
Cumulative
f
Score cuts (10 equal-sized buckets)
Score
Cumulative volume
g
Tag vs Score Splits Comparison
Total Percentage by Tag
~1861 p/split
1 (0-0.8)
2 (0.8-2.3)
3 (2.3-7.2)
4 (7.2-73)
5 (73-98)
6 (98-99)
7 (99-100)
8 (100-100)
Tag
background
bound
h
ROC Curve: Combined cross-validation sets
Sensitivity [True Positive Rate]
1 - Specificity [False Positive Rate]
AUC = 98.22
95% CI: 98.01-98.43

Fig. S25

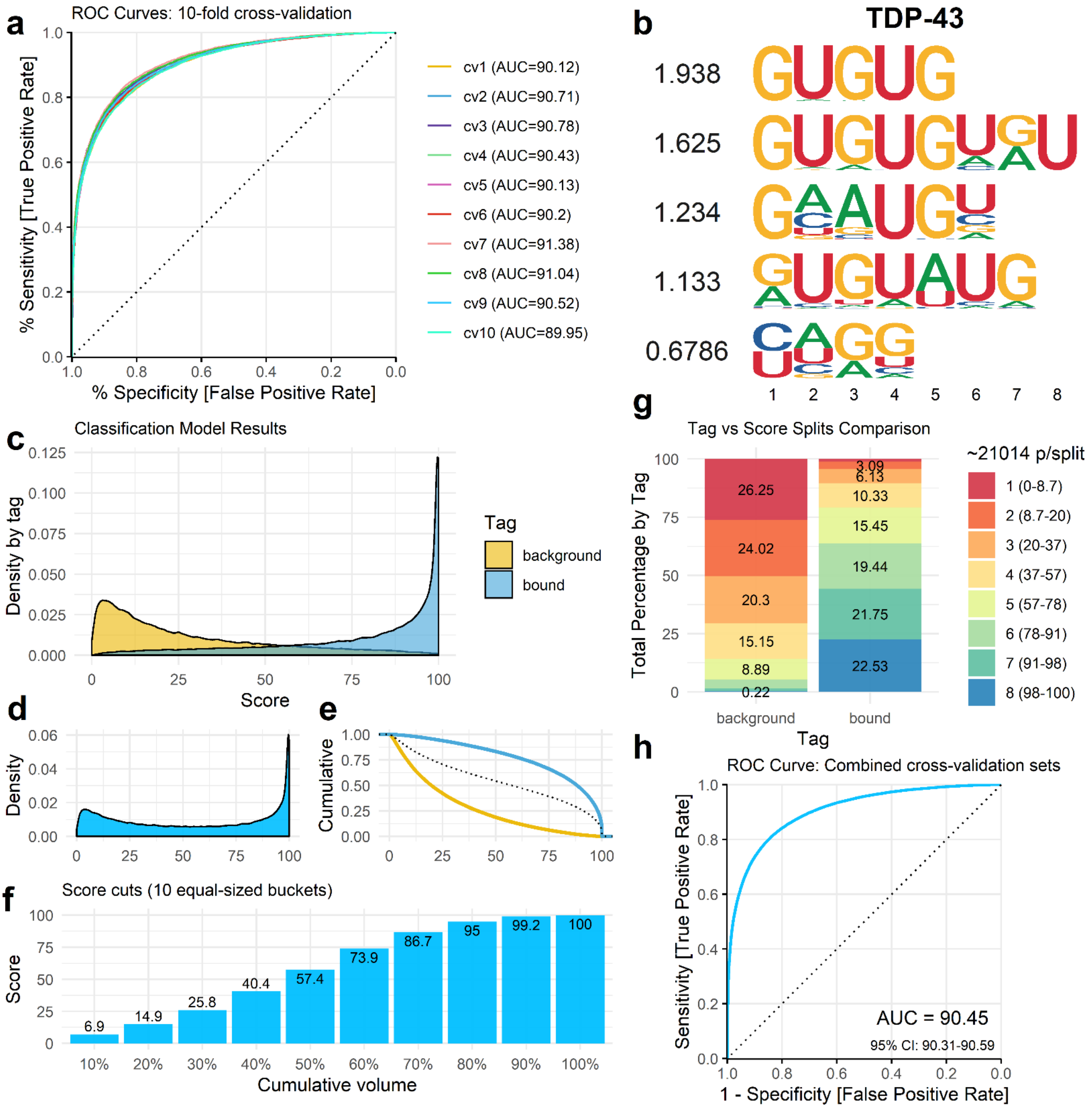

a
ROC Curves: 10-fold cross-validation
% Sensitivity [True Positive Rate]
% Specificity [False Positive Rate]
cv1 (AUC=90.12)
cv2 (AUC=90.71)
cv3 (AUC=90.78)
cv4 (AUC=90.43)
cv5 (AUC=90.13)
cv6 (AUC=90.2)
cv7 (AUC=91.38)
cv8 (AUC=91.04)
cv9 (AUC=90.52)
cv10 (AUC=89.95)
b
TDP-43
1.938
1.625
1.234
1.133
0.6786
c
Classification Model Results
Density by tag
Score
Tag
background
bound
d
Density
e
Cumulative
f
Score cuts (10 equal-sized buckets)
Score
Cumulative volume
6.9
14.9
25.8
40.4
57.4
73.9
86.7
95
99.2
100
g
Tag vs Score Splits Comparison
Total Percentage by Tag
~21014 p/split
1 (0-8.7)
2 (8.7-20)
3 (20-37)
4 (37-57)
5 (57-78)
6 (78-91)
7 (91-98)
8 (98-100)
26.25
24.02
20.3
15.15
8.89
0.22
3.09
6.13
10.33
15.45
19.44
21.75
22.53
background
bound
Tag
h
ROC Curve: Combined cross-validation sets
Sensitivity [True Positive Rate]
1 - Specificity [False Positive Rate]
AUC = 90.45
95% CI: 90.31-90.59

Fig. S26

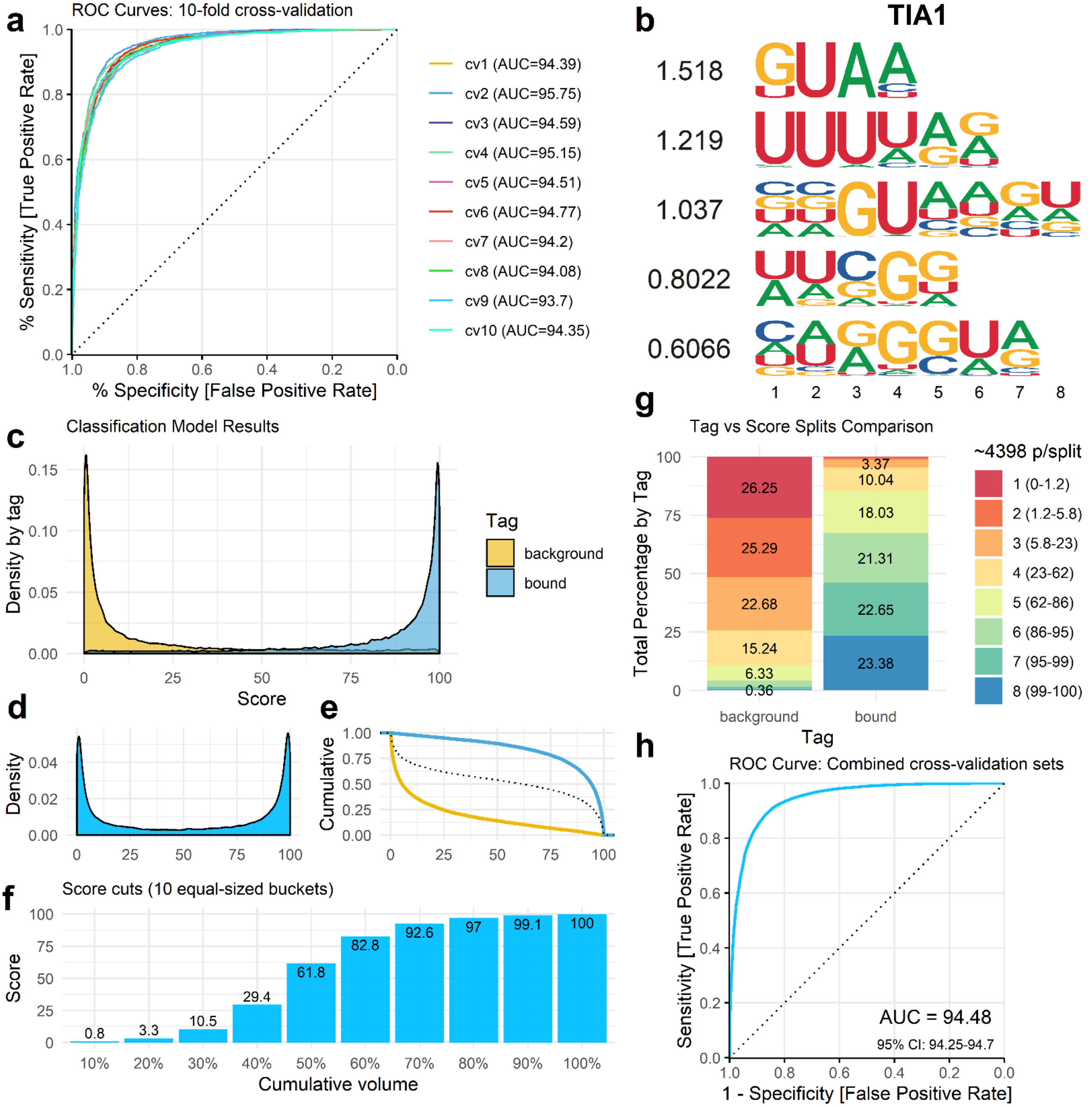
a
ROC Curves: 10-fold cross-validation
% Sensitivity [True Positive Rate]
% Specificity [False Positive Rate]
cv1 (AUC=94.39)
cv2 (AUC=95.75)
cv3 (AUC=94.59)
cv4 (AUC=95.15)
cv5 (AUC=94.51)
cv6 (AUC=94.77)
cv7 (AUC=94.2)
cv8 (AUC=94.08)
cv9 (AUC=93.7)
cv10 (AUC=94.35)
b
TIA1
1.518
1.219
1.037
0.8022
0.6066
c
Classification Model Results
Density by tag
Score
Tag
background
bound
d
Density
e
Cumulative
f
Score cuts (10 equal-sized buckets)
Score
Cumulative volume
0.8
3.3
10.5
29.4
61.8
82.8
92.6
97
99.1
100
g
Tag vs Score Splits Comparison
Total Percentage by Tag
~4398 p/split
1 (0-1.2)
2 (1.2-5.8)
3 (5.8-23)
4 (23-62)
5 (62-86)
6 (86-95)
7 (95-99)
8 (99-100)
26.25
25.29
22.68
15.24
6.33
0.36
3.37
10.04
18.03
21.31
22.65
23.38
background
bound
Tag
h
ROC Curve: Combined cross-validation sets
Sensitivity [True Positive Rate]
1 - Specificity [False Positive Rate]
AUC = 94.48
95% CI: 94.25-94.7

Fig. S27

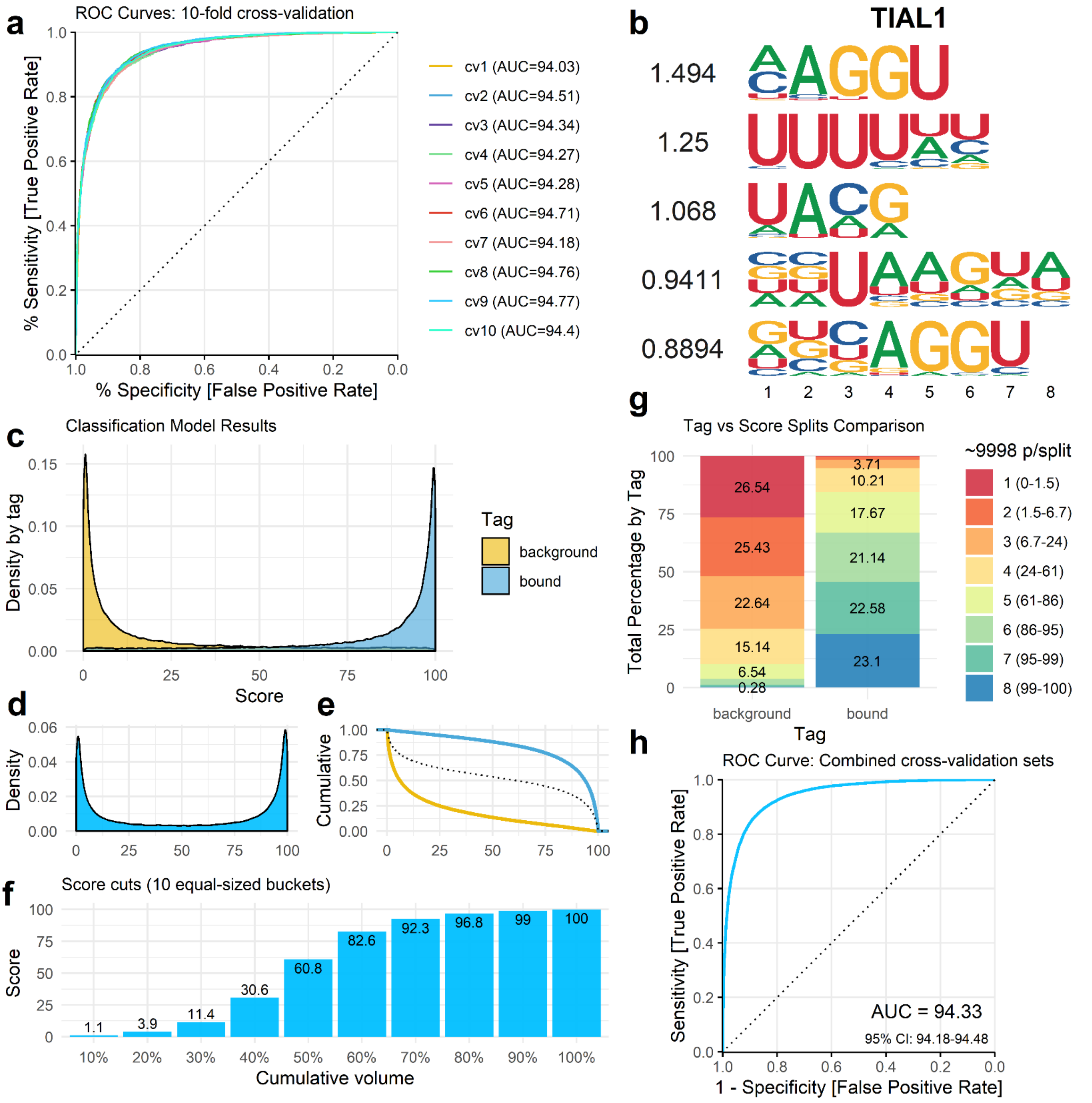
a
ROC Curves: 10-fold cross-validation
% Sensitivity [True Positive Rate]
% Specificity [False Positive Rate]
cv1 (AUC=94.03)
cv2 (AUC=94.51)
cv3 (AUC=94.34)
cv4 (AUC=94.27)
cv5 (AUC=94.28)
cv6 (AUC=94.71)
cv7 (AUC=94.18)
cv8 (AUC=94.76)
cv9 (AUC=94.77)
cv10 (AUC=94.4)
b
TIAL1
1.494
1.25
1.068
0.9411
0.8894
c
Classification Model Results
Density by tag
Score
Tag
background
bound
d
Density
e
Cumulative
f
Score cuts (10 equal-sized buckets)
Score
Cumulative volume
1.1
3.9
11.4
30.6
60.8
82.6
92.3
96.8
99
100
g
Tag vs Score Splits Comparison
Total Percentage by Tag
~9998 p/split
1 (0-1.5)
2 (1.5-6.7)
3 (6.7-24)
4 (24-61)
5 (61-86)
6 (86-95)
7 (95-99)
8 (99-100)
26.54
25.43
22.64
15.14
6.54
0.28
3.71
10.21
17.67
21.14
22.58
23.1
background
bound
Tag
h
ROC Curve: Combined cross-validation sets
Sensitivity [True Positive Rate]
1 - Specificity [False Positive Rate]
AUC = 94.33
95% CI: 94.18-94.48

Fig. S28

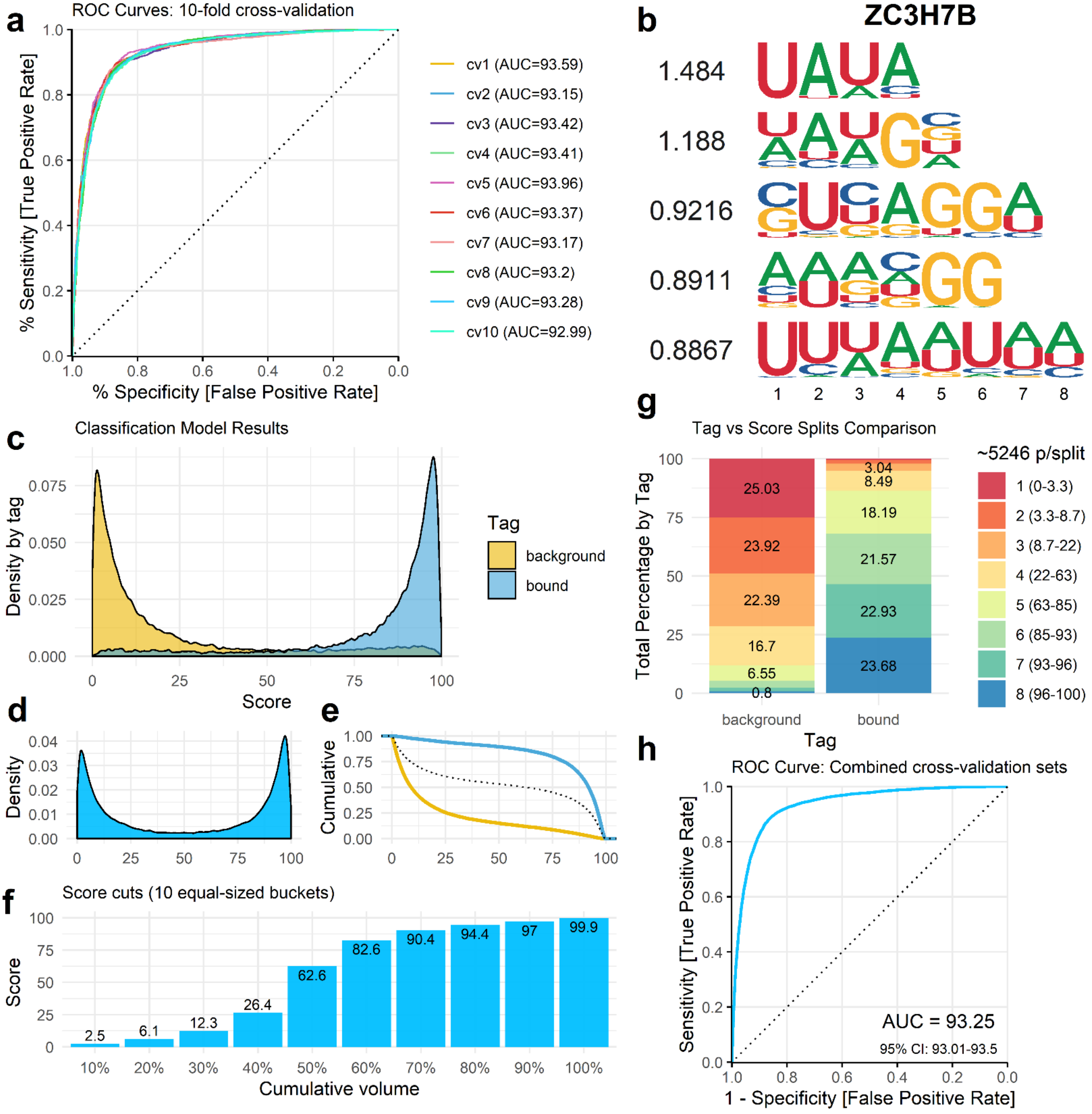
a
ROC Curves: 10-fold cross-validation
% Sensitivity [True Positive Rate]
% Specificity [False Positive Rate]
cv1 (AUC=93.59)
cv2 (AUC=93.15)
cv3 (AUC=93.42)
cv4 (AUC=93.41)
cv5 (AUC=93.96)
cv6 (AUC=93.37)
cv7 (AUC=93.17)
cv8 (AUC=93.2)
cv9 (AUC=93.28)
cv10 (AUC=92.99)
b
ZC3H7B
1.484
1.188
0.9216
0.8911
0.8867
c
Classification Model Results
Density by tag
Score
Tag
background
bound
d
Density
e
Cumulative
f
Score cuts (10 equal-sized buckets)
Score
Cumulative volume
2.5
6.1
12.3
26.4
62.6
82.6
90.4
94.4
97
99.9
g
Tag vs Score Splits Comparison
Total Percentage by Tag
~5246 p/split
1 (0-3.3)
2 (3.3-8.7)
3 (8.7-22)
4 (22-63)
5 (63-85)
6 (85-93)
7 (93-96)
8 (96-100)
25.03
23.92
22.39
16.7
6.55
0.8
3.04
8.49
18.19
21.57
22.93
23.68
background
bound
Tag
h
ROC Curve: Combined cross-validation sets
Sensitivity [True Positive Rate]
1 - Specificity [False Positive Rate]
AUC = 93.25
95% CI: 93.01-93.5

Fig. S29

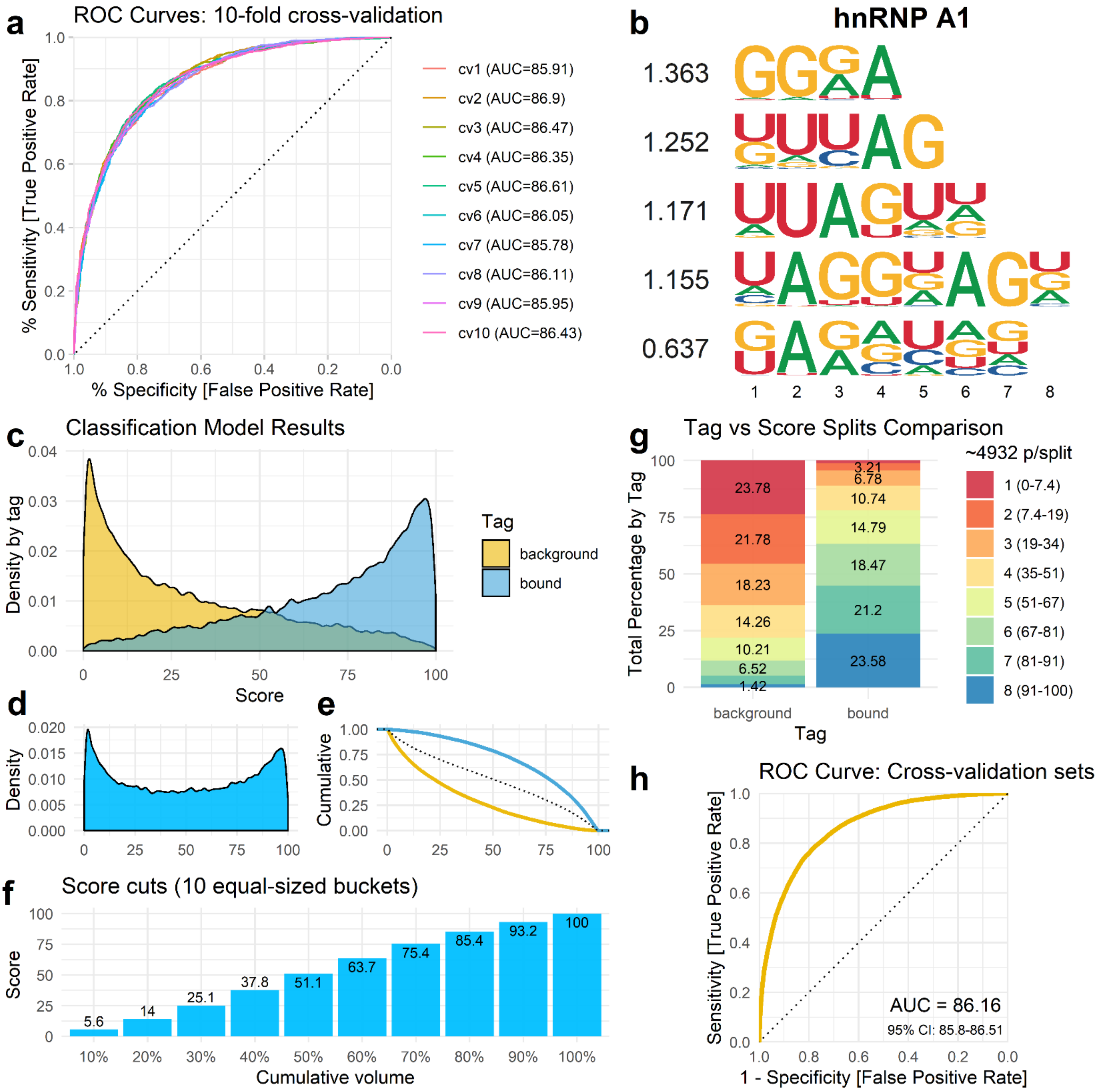

a
ROC Curves: 10-fold cross-validation
% Sensitivity [True Positive Rate]
% Specificity [False Positive Rate]
cv1 (AUC=85.91)
cv2 (AUC=86.9)
cv3 (AUC=86.47)
cv4 (AUC=86.35)
cv5 (AUC=86.61)
cv6 (AUC=86.05)
cv7 (AUC=85.78)
cv8 (AUC=86.11)
cv9 (AUC=85.95)
cv10 (AUC=86.43)
b
hnRNP A1
1.363
1.252
1.171
1.155
0.637
c
Classification Model Results
Density by tag
Score
Tag
background
bound
d
Density
e
Cumulative
f
Score cuts (10 equal-sized buckets)
Score
Cumulative volume
5.6
14
25.1
37.8
51.1
63.7
75.4
85.4
93.2
100
g
Tag vs Score Splits Comparison
Total Percentage by Tag
Tag
~4932 p/split
1 (0-7.4)
2 (7.4-19)
3 (19-34)
4 (35-51)
5 (51-67)
6 (67-81)
7 (81-91)
8 (91-100)
23.78
21.78
18.23
14.26
10.21
6.52
1.42
3.21
6.78
10.74
14.79
18.47
21.2
23.58
h
ROC Curve: Cross-validation sets
Sensitivity [True Positive Rate]
1 - Specificity [False Positive Rate]
AUC = 86.16
95% CI: 85.8-86.51

Fig. S30

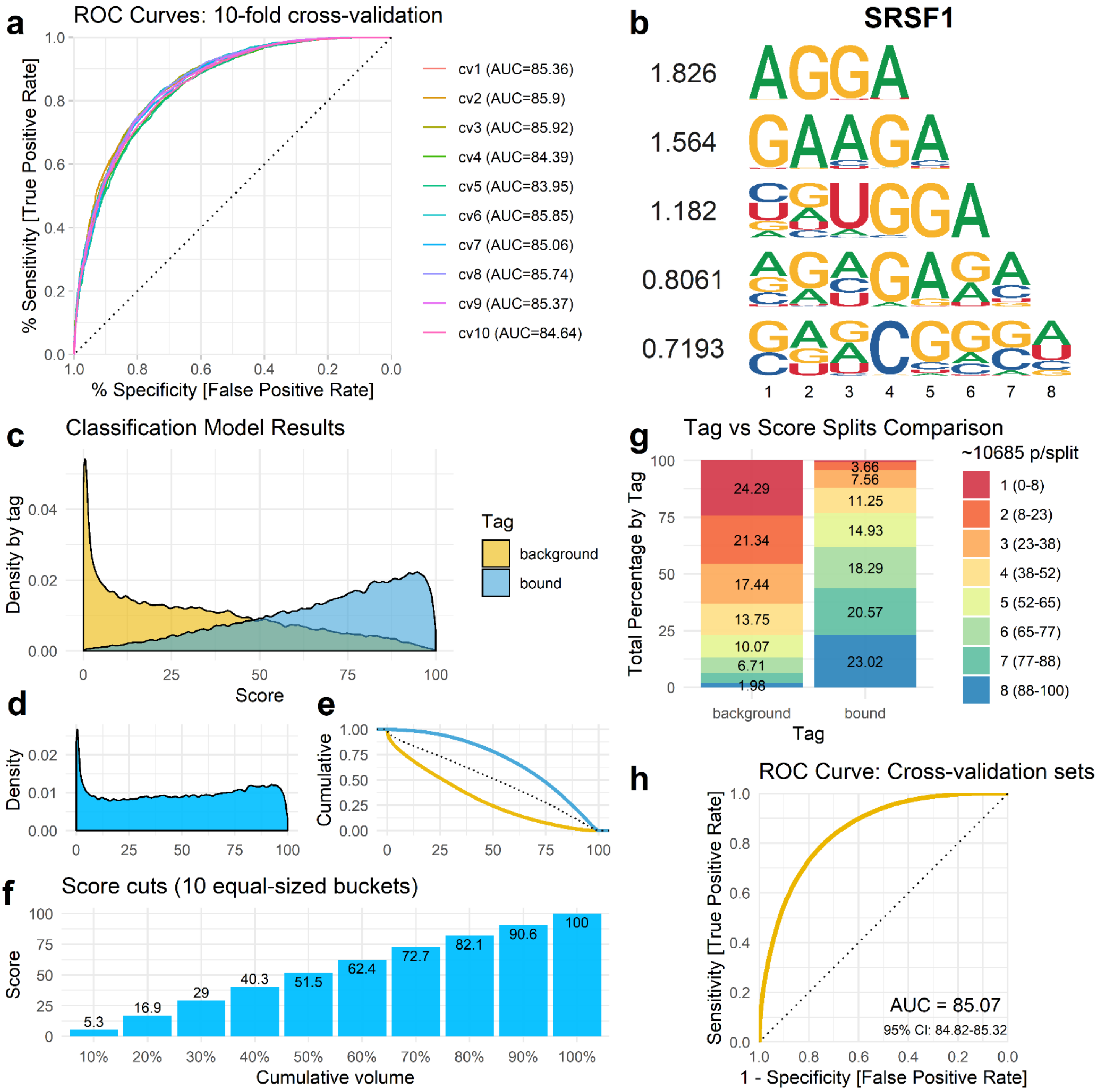
a
ROC Curves: 10-fold cross-validation
% Sensitivity [True Positive Rate]
% Specificity [False Positive Rate]
cv1 (AUC=85.36)
cv2 (AUC=85.9)
cv3 (AUC=85.92)
cv4 (AUC=84.39)
cv5 (AUC=83.95)
cv6 (AUC=85.85)
cv7 (AUC=85.06)
cv8 (AUC=85.74)
cv9 (AUC=85.37)
cv10 (AUC=84.64)
b
SRSF1
1.826
1.564
1.182
0.8061
0.7193
c
Classification Model Results
Density by tag
Score
Tag
background
bound
d
Density
e
Cumulative
f
Score cuts (10 equal-sized buckets)
Score
Cumulative volume
5.3
16.9
29
40.3
51.5
62.4
72.7
82.1
90.6
100
g
Tag vs Score Splits Comparison
Total Percentage by Tag
Tag
~10685 p/split
1 (0-8)
2 (8-23)
3 (23-38)
4 (38-52)
5 (52-65)
6 (65-77)
7 (77-88)
8 (88-100)
background
24.29
21.34
17.44
13.75
10.07
6.71
1.98
bound
3.66
7.56
11.25
14.93
18.29
20.57
23.02
h
ROC Curve: Cross-validation sets
Sensitivity [True Positive Rate]
1 - Specificity [False Positive Rate]
AUC = 85.07
95% CI: 84.82-85.32

Fig. S31

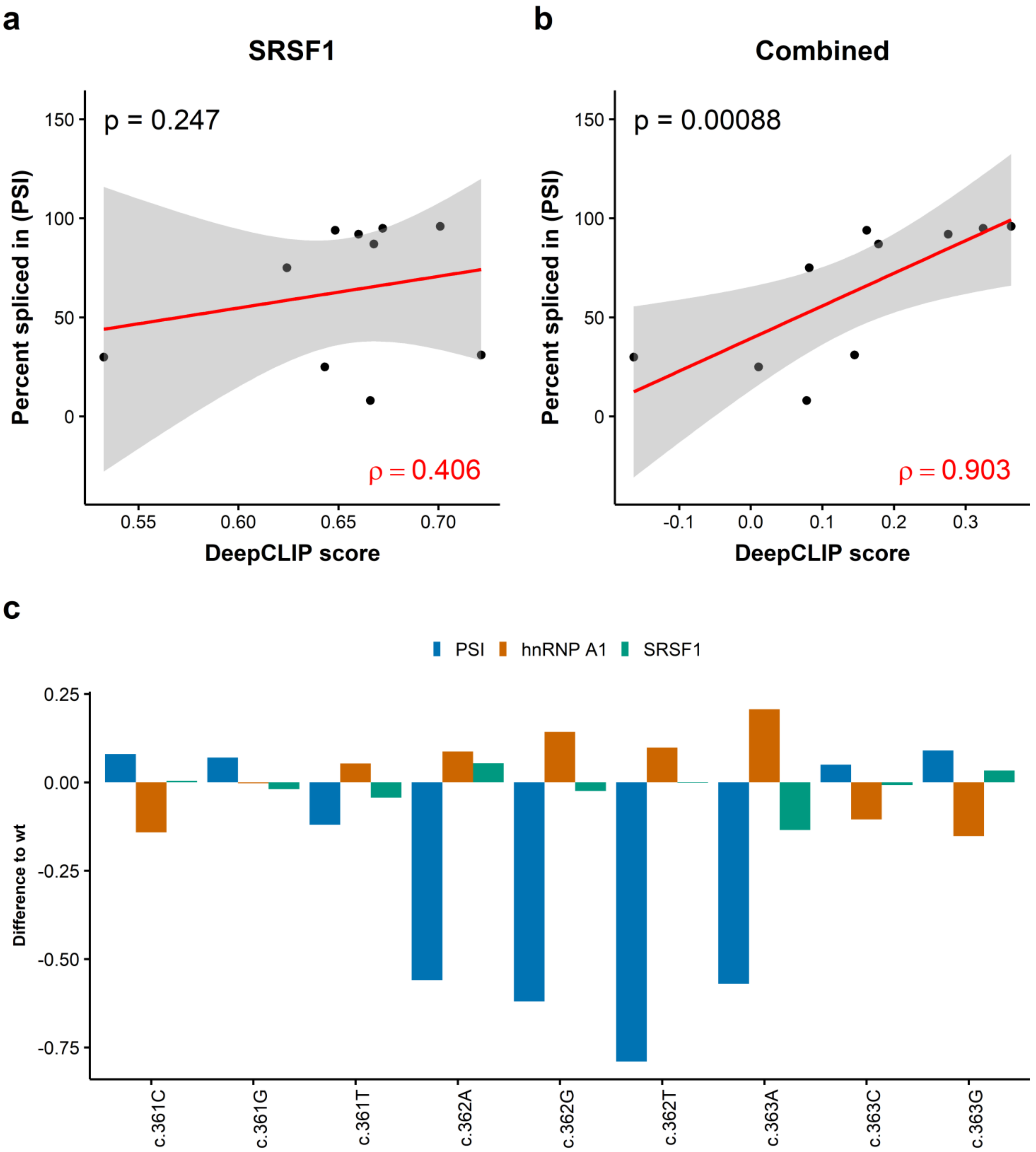
a
SRSF1
p = 0.247
ρ = 0.406
Percent spliced in (PSI)
DeepCLIP score
150
100
50
0
0.55
0.60
0.65
0.70
b
Combined
p = 0.00088
ρ = 0.903
Percent spliced in (PSI)
DeepCLIP score
-0.1
0.0
0.1
0.2
0.3
c
PSI
hnRNP A1
SRSF1
Difference to wt
0.25
0.00
-0.25
-0.50
-0.75
c.361C
c.361G
c.361T
c.362A
c.362G
c.362T
c.363A
c.363C
c.363G

Fig. S32

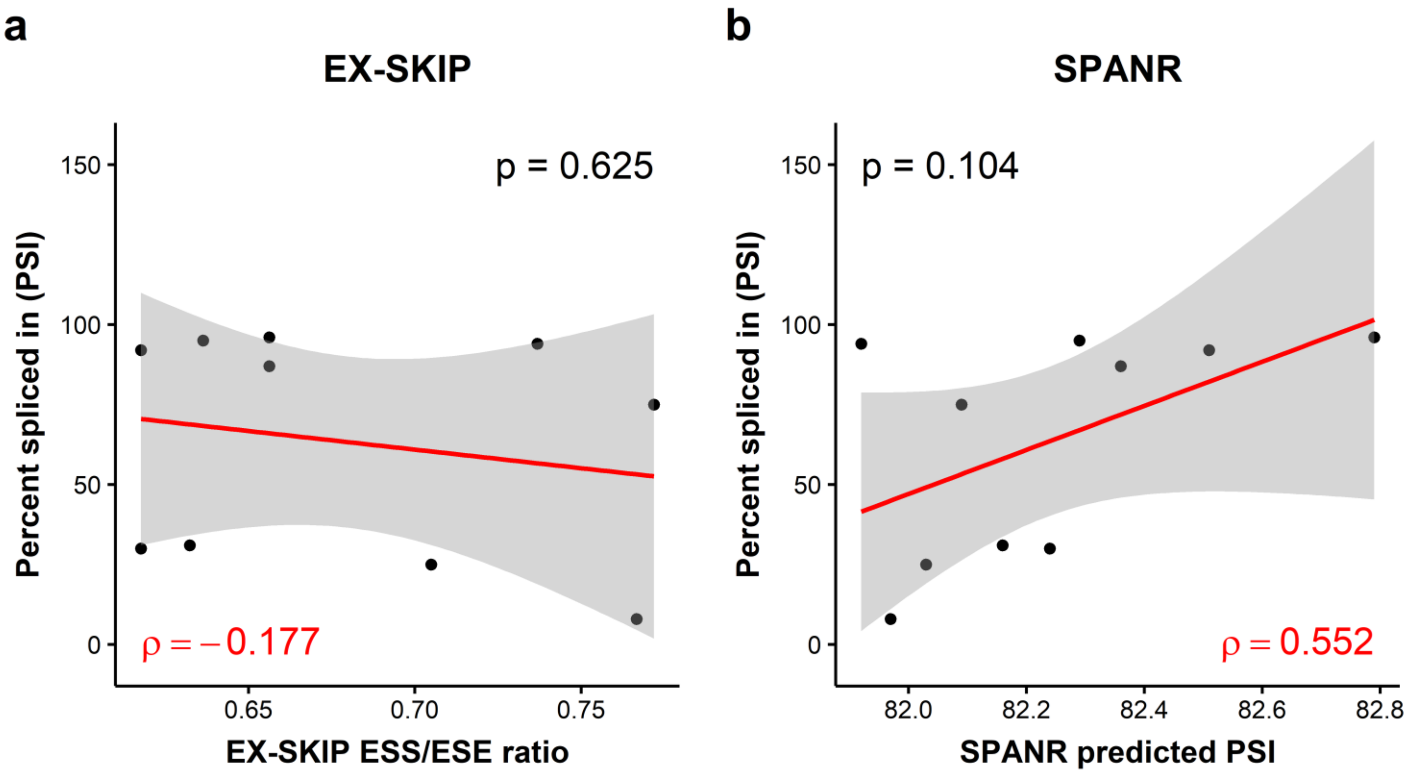

a
EX-SKIP
p = 0.625
ρ = −0.177
Percent spliced in (PSI)
EX-SKIP ESS/ESE ratio
0
50
100
150
0.65
0.70
0.75
b
SPANR
p = 0.104
ρ = 0.552
Percent spliced in (PSI)
SPANR predicted PSI
0
50
100
150
82.0
82.2
82.4
82.6
82.8

Fig. S33

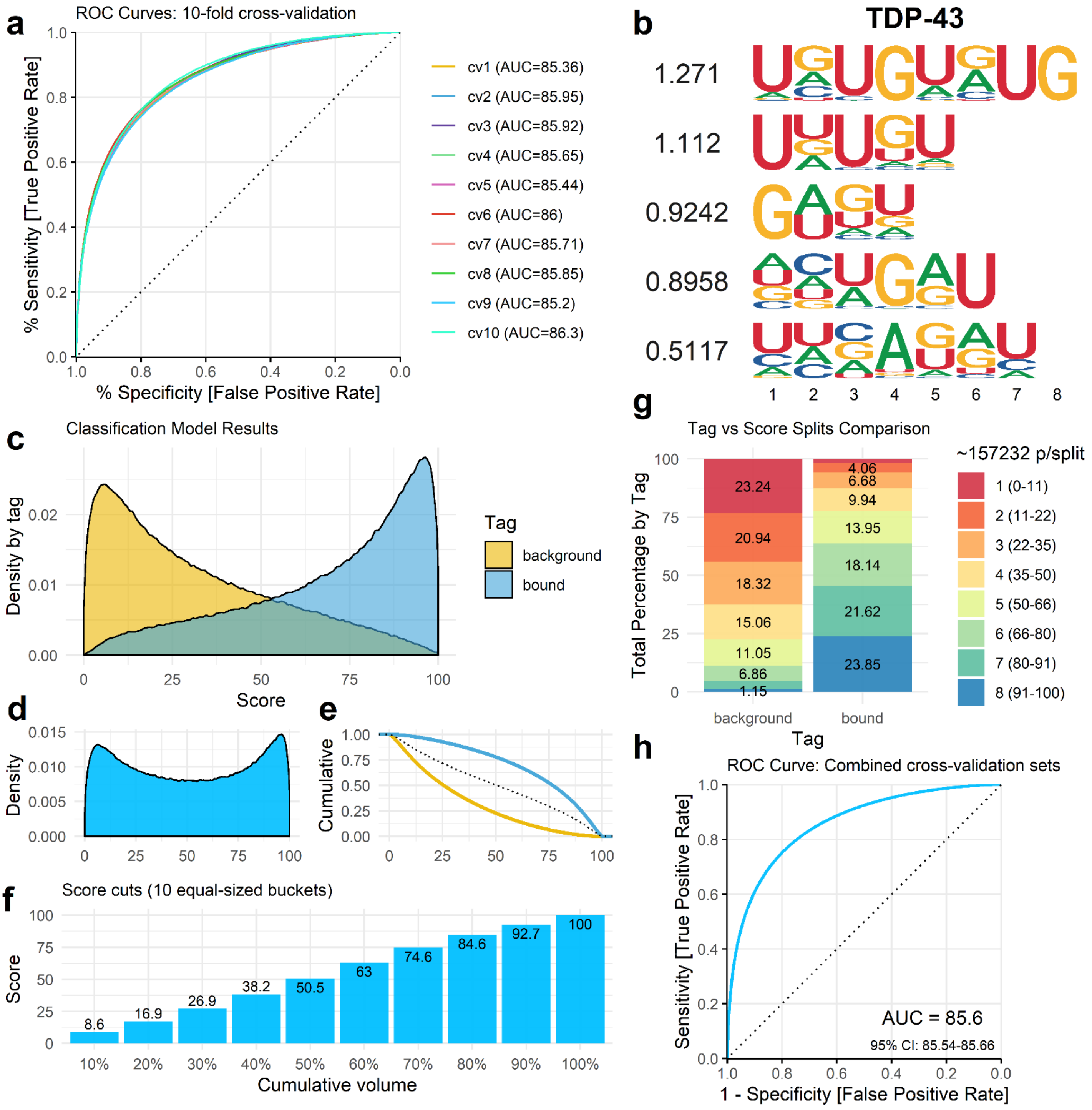

a
ROC Curves: 10-fold cross-validation
% Sensitivity [True Positive Rate]
% Specificity [False Positive Rate]
cv1 (AUC=85.36)
cv2 (AUC=85.95)
cv3 (AUC=85.92)
cv4 (AUC=85.65)
cv5 (AUC=85.44)
cv6 (AUC=86)
cv7 (AUC=85.71)
cv8 (AUC=85.85)
cv9 (AUC=85.2)
cv10 (AUC=86.3)
b
TDP-43
1.271
1.112
0.9242
0.8958
0.5117
c
Classification Model Results
Density by tag
Score
Tag
background
bound
d
Density
e
Cumulative
f
Score cuts (10 equal-sized buckets)
Score
Cumulative volume
8.6
16.9
26.9
38.2
50.5
63
74.6
84.6
92.7
100
g
Tag vs Score Splits Comparison
Total Percentage by Tag
~157232 p/split
1 (0-11)
2 (11-22)
3 (22-35)
4 (35-50)
5 (50-66)
6 (66-80)
7 (80-91)
8 (91-100)
23.24
20.94
18.32
15.06
11.05
6.86
1.15
4.06
6.68
9.94
13.95
18.14
21.62
23.85
background
bound
Tag
h
ROC Curve: Combined cross-validation sets
Sensitivity [True Positive Rate]
1 - Specificity [False Positive Rate]
AUC = 85.6
95% CI: 85.54-85.66

Fig. S34

hnRNP A1 SPRi binding plots with CLAMP model data

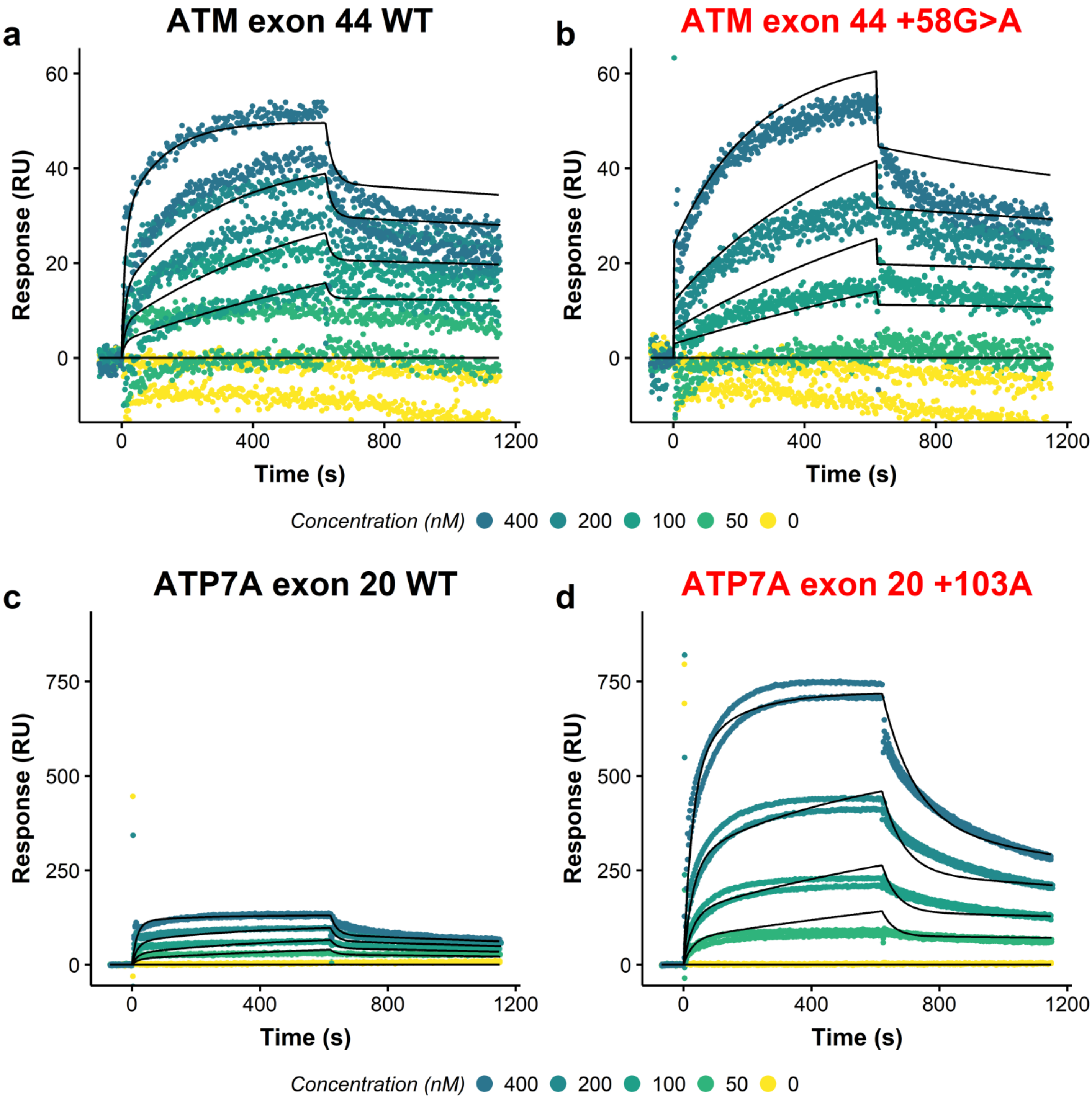

Fig. S35

hnRNP A1 SPRi binding plots with CLAMP model data

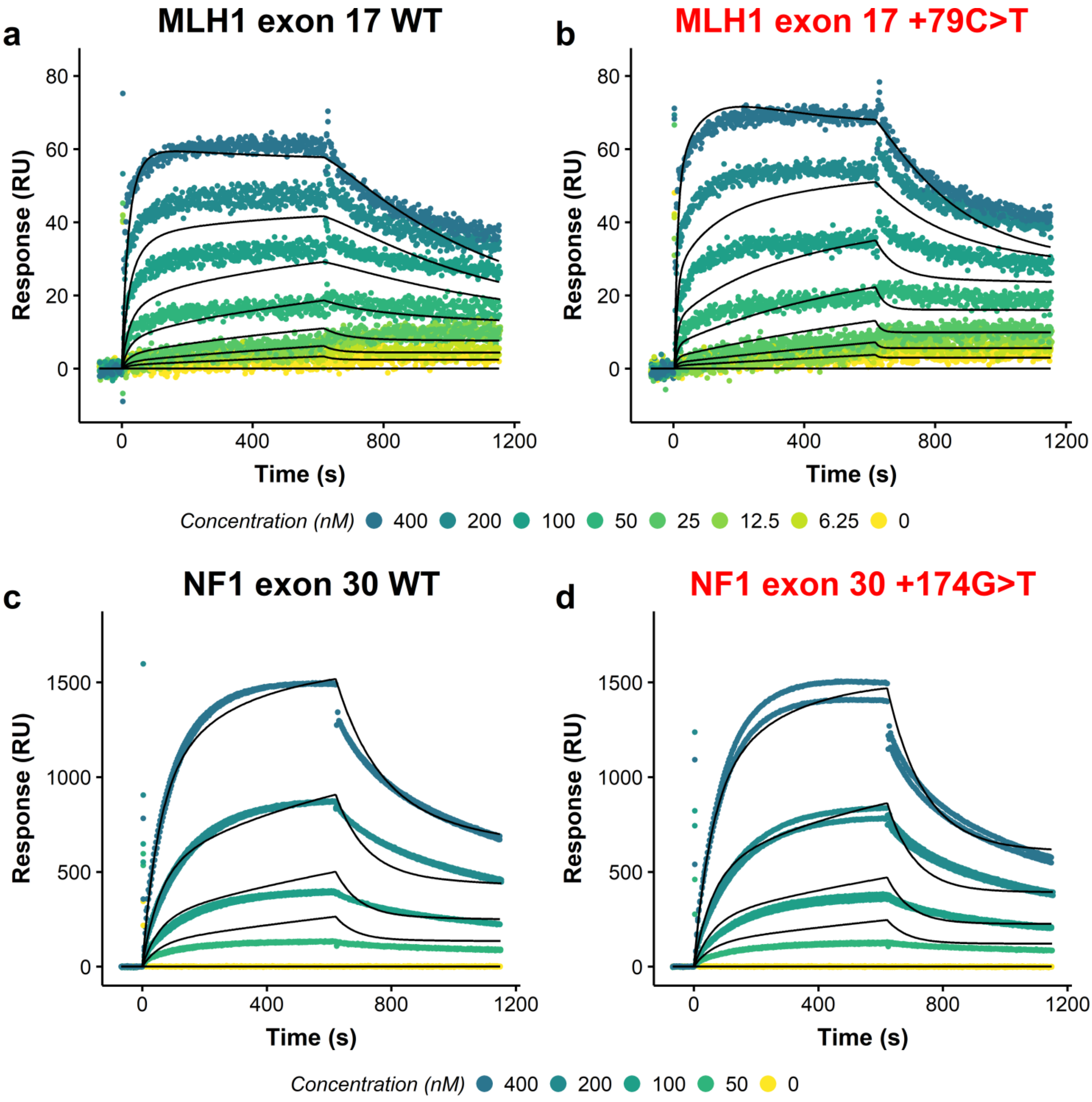

Fig. S36

hnRNP A1 SPRi binding plots with CLAMP model data

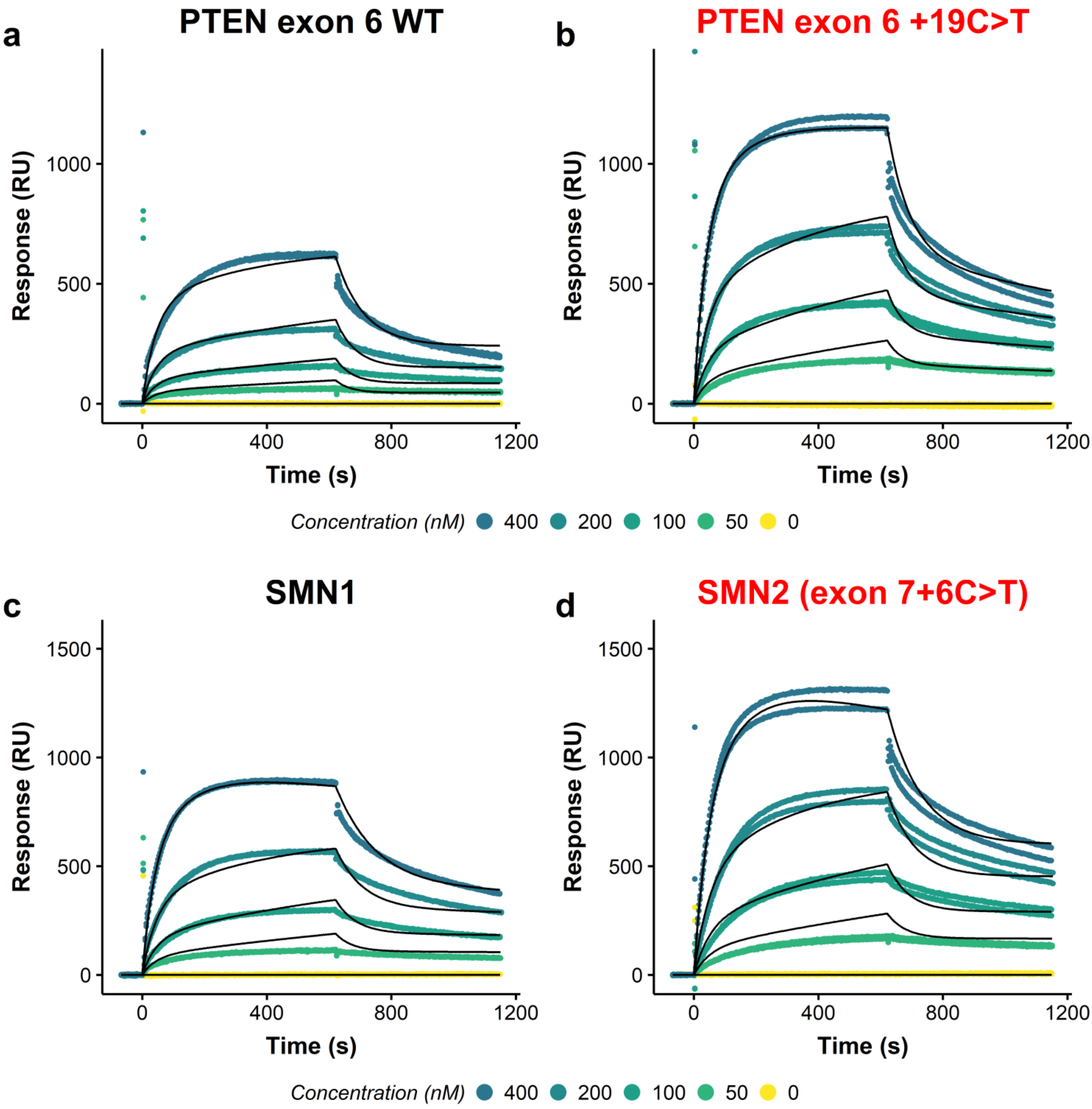

Fig. S37

SRSF1 SPRi binding plots with CLAMP model data

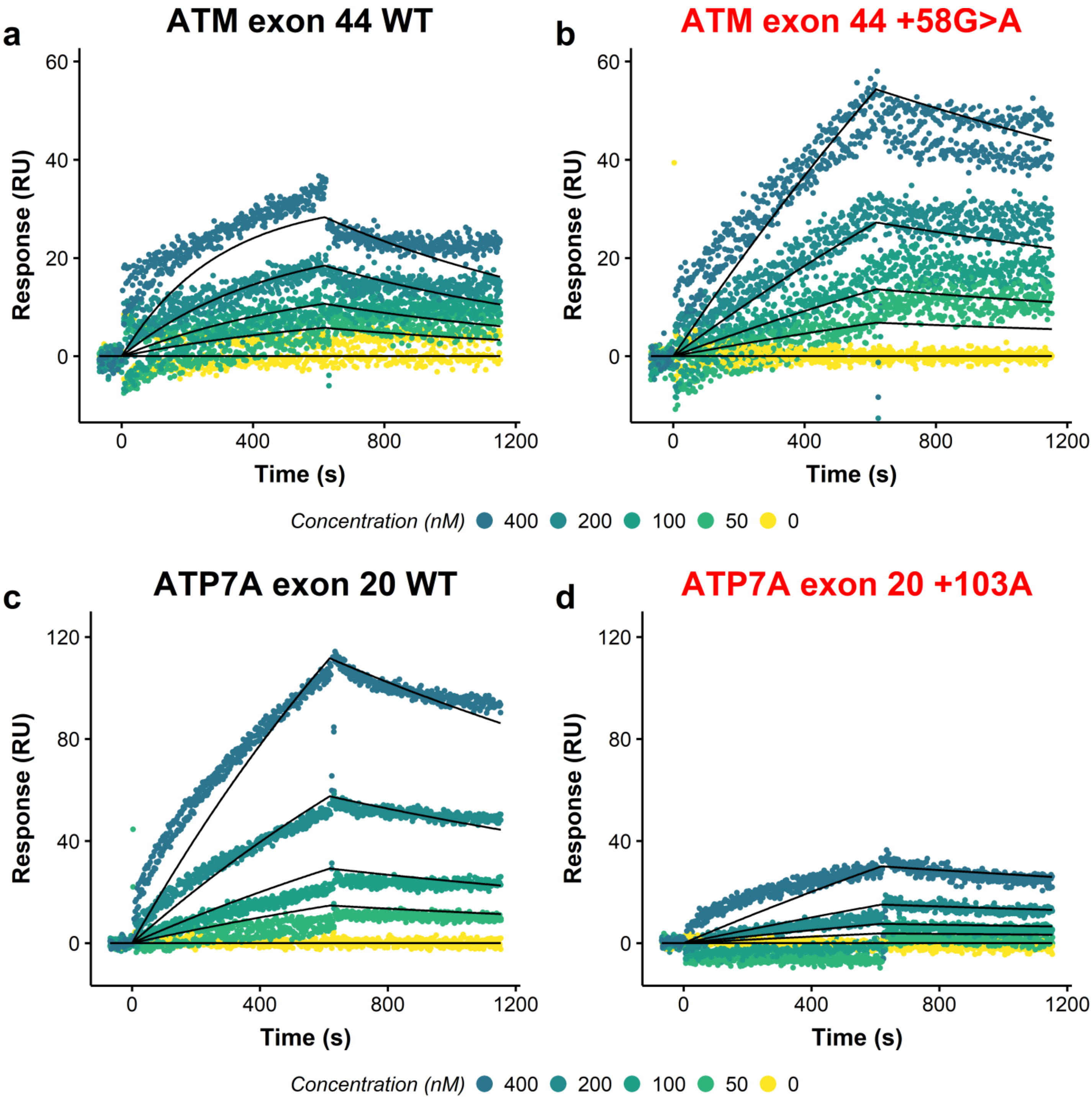

Fig. S38

SRSF1 SPRi binding plots with CLAMP model data

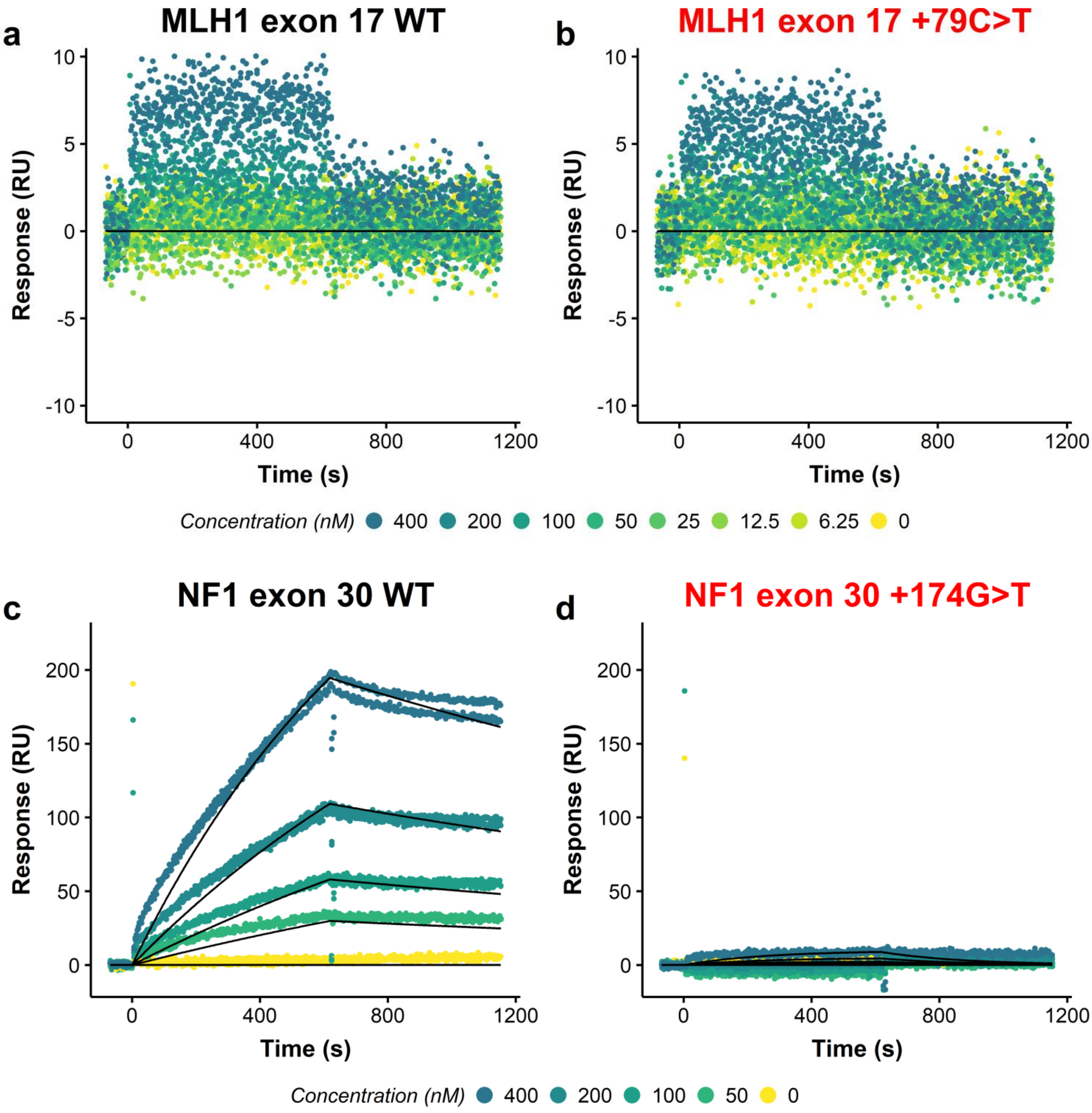

Fig. S39

SRSF1 SPRi binding plots with CLAMP model data

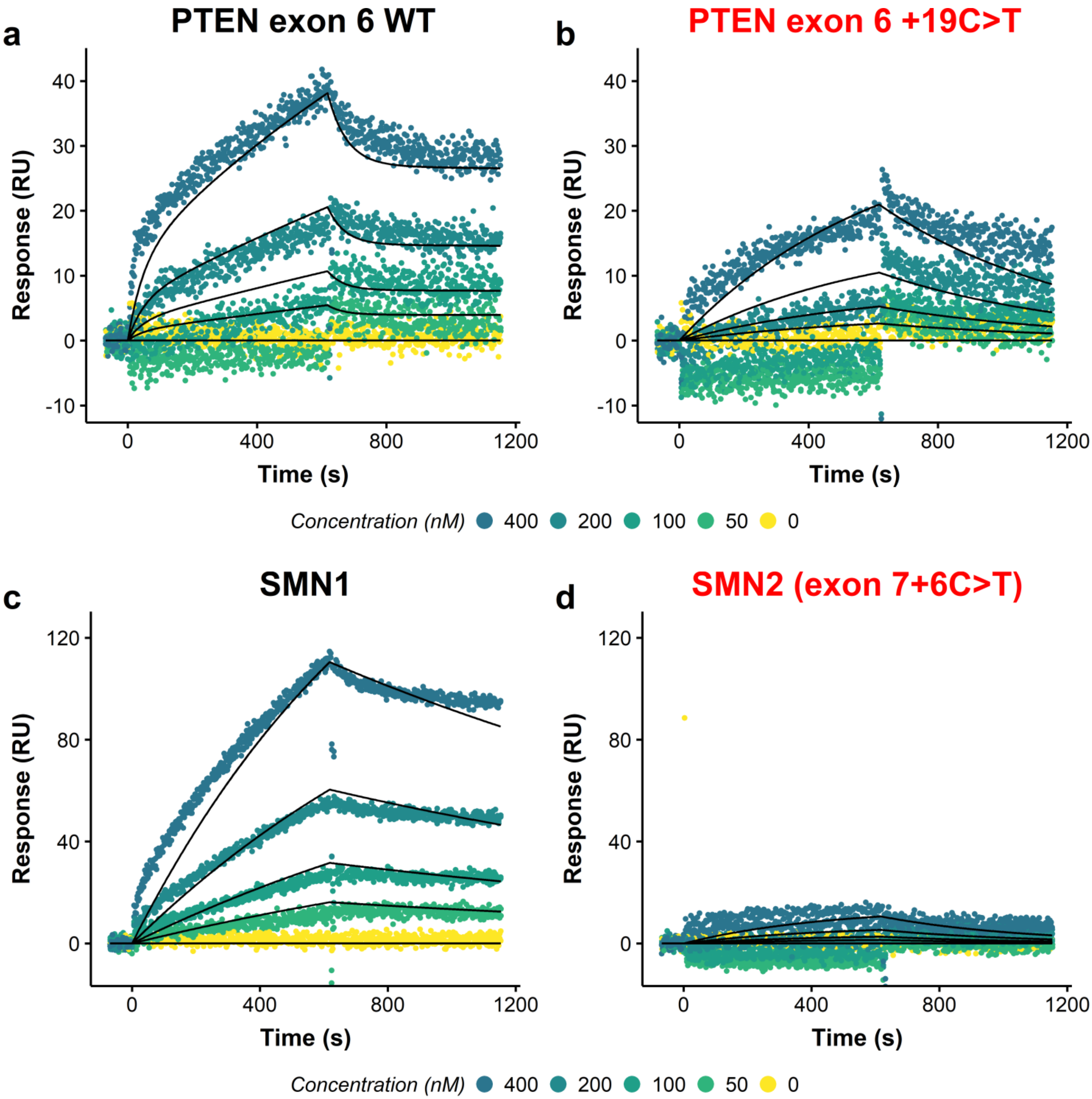

Fig. S40

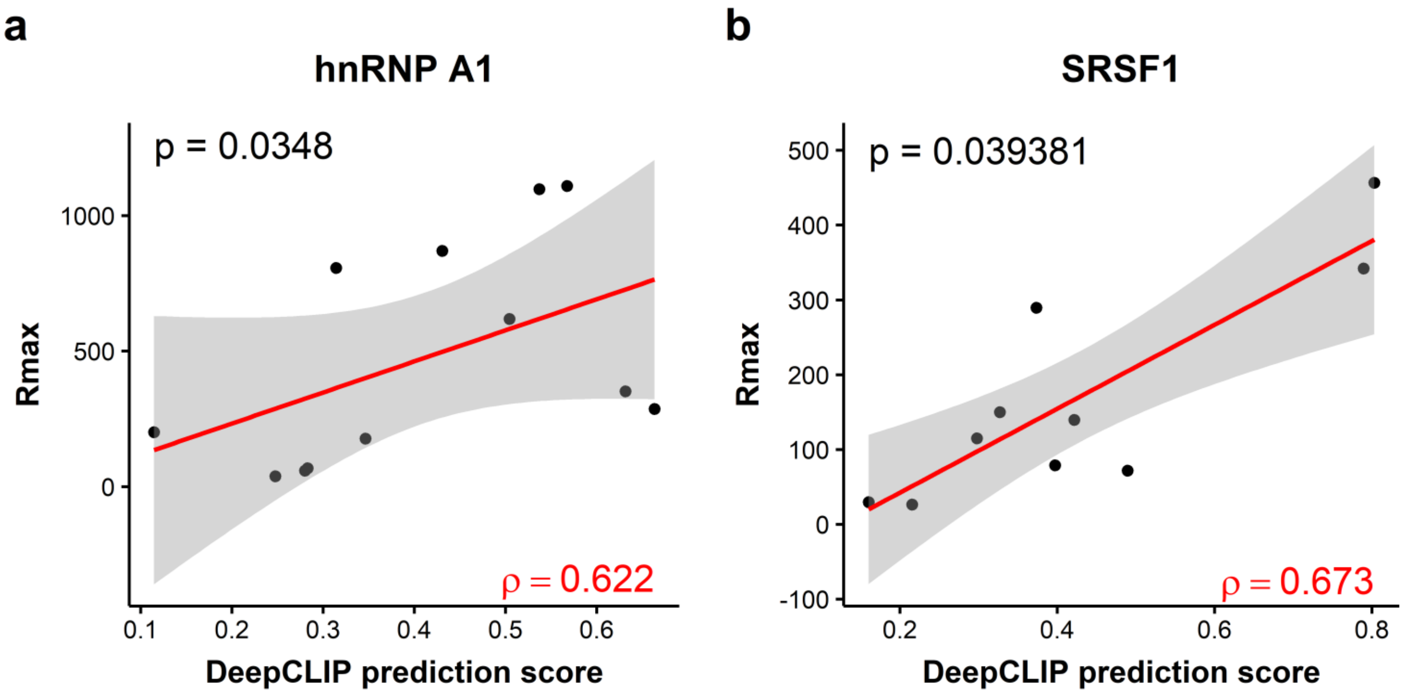

a
hnRNP A1
p = 0.0348
ρ = 0.622
Rmax
DeepCLIP prediction score
b
SRSF1
p = 0.039381
ρ = 0.673
Rmax
DeepCLIP prediction score

Fig. S41

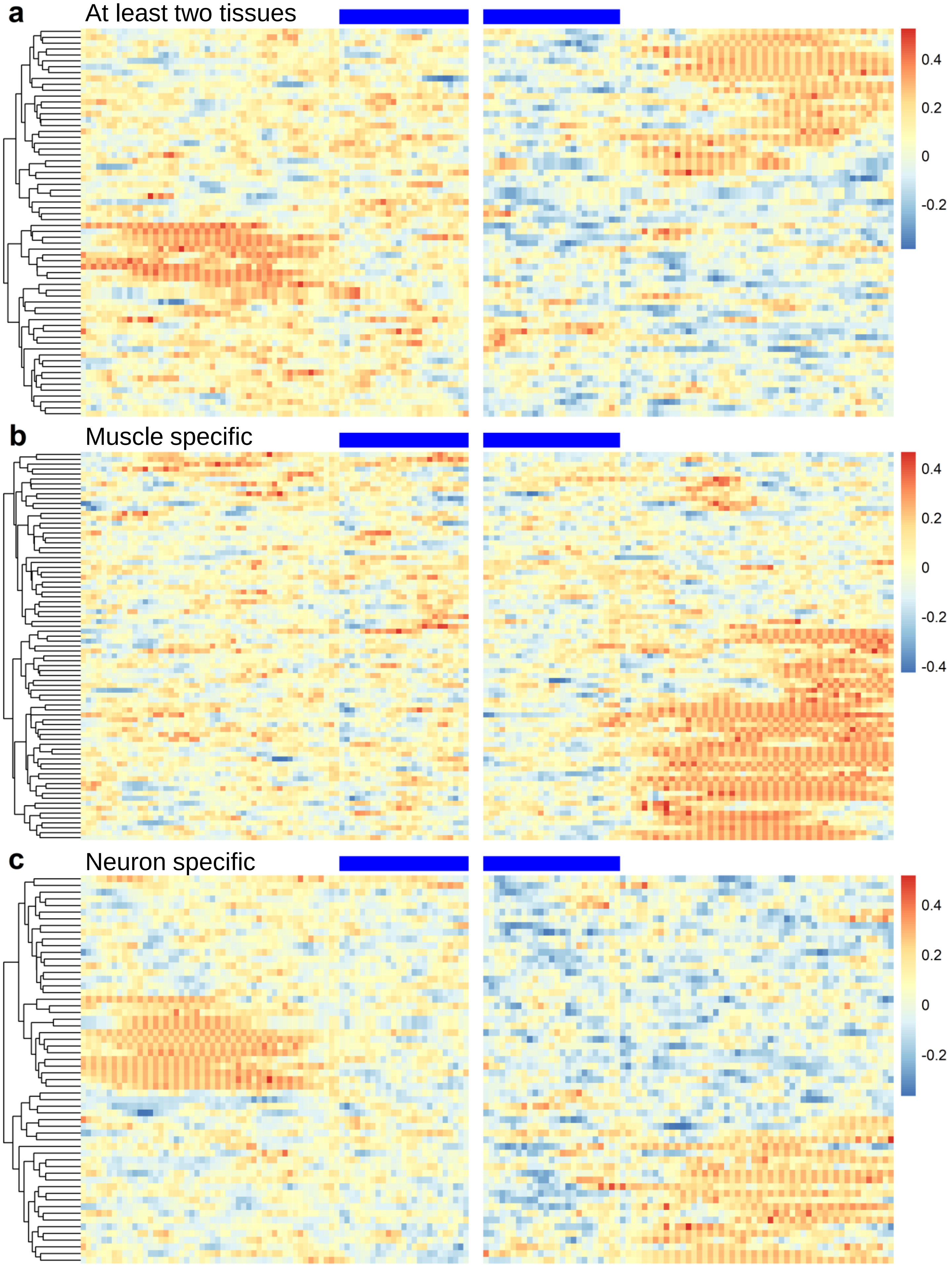
a
At least two tissues
0.4
0.2
0
-0.2
b
Muscle specific
0.4
0.2
0
-0.2
-0.4
c
Neuron specific
0.4
0.2
0
-0.2
